# Delineating mouse β-cell identity during lifetime and in diabetes with a single cell atlas

**DOI:** 10.1101/2022.12.22.521557

**Authors:** Karin Hrovatin, Aimée Bastidas-Ponce, Mostafa Bakhti, Luke Zappia, Maren Büttner, Ciro Sallino, Michael Sterr, Anika Böttcher, Adriana Migliorini, Heiko Lickert, Fabian J. Theis

## Abstract

Multiple pancreatic islet single-cell RNA sequencing (scRNA-seq) datasets have been generated to study development, homeostasis, and diabetes. However, there is no consensus on cell states and pathways across conditions as well as the value of preclinical mouse models. Since these challenges can only be resolved by jointly analyzing multiple datasets, we present a scRNA-seq cross-condition mouse islet atlas (MIA). We integrated over 300,000 cells from nine datasets with 56 samples, varying in age, sex, and diabetes models, including an autoimmune type 1 diabetes (T1D) model (NOD), a gluco-/lipotoxicity T2D model (db/db), and a chemical streptozotocin (STZ) β-cell ablation model. MIA is a curated resource for interactive exploration and computational querying, providing new insights inaccessible from individual datasets. The β-cell landscape of MIA revealed new disease progression cell states and cross-publication differences between previously suggested marker genes. We show that in the STZ model β-cells transcriptionally correlate to human T2D and mouse db/db, but are less similar to human T1D and mouse NOD. We observe different pathways shared between immature, aged, and diabetes model β-cells. In conclusion, our work presents the first comprehensive analysis of β-cell responses to different stressors, providing a roadmap for the understanding of β-cell plasticity, compensation, and demise.

## Introduction

The major hallmark of diabetes mellitus is impaired glucose homeostasis. Blood glucose is regulated by multiple hormones secreted from pancreatic islets of Langerhans that are comprised of insulin-producing β-cells, which are main acters in diabetes, as well as glucagon-producing α-cells, somatostatin-producing δ-cells, pancreatic polypeptide-producing γ-cells, and ghrelin-producing ε-cells^1–3^. Type 1 (T1D) and type 2 diabetes (T2D) arise due to the loss or progressive dysfunction of β-cells, respectively. Current anti-diabetic medications do not lead to remission^4,5^, while more effective treatments, such as bariatric surgery and islet transplantation, are highly invasive or can be only offered to a small number of patients^6,7^. The central role of β-cells in diabetes development urges the establishment of new therapies that focus on restoring β-cell mass and function^8–13^. Achieving such strategies requires deeper understanding of β-cell heterogeneity, maturation, function, and failure^14–16^.

Shortly after birth, β-cells are immature, defined by poor glucose-stimulated insulin secretion (GSIS) ^17^. Immature β-cells gain functional maturation, as defined by expression of several markers, including Urocortin-3, Flattop, MafA, and glucose transporter 2 (Glut2), and accurate GSIS in the first weeks after birth and again after weaning^17–20^. Adult β-cells also differ within and across phenotypes and conditions^15,19^. For instance, insulin production and secretion of β-cells is changed due to healthy aging or stress-induced senescence^21–26^. Function also differs between sexes, with male β-cells having transcriptomic signatures more akin to T2D^27^.

Different stressors can lead to β-cell failure, which is often studied with mouse models^28,29^. T2D is marked by gluco-/lipotoxicity leading to β-cell dedifferentiation, compensatory insulin production, and resulting endoplasmic reticulum (ER) stress^30,31^, all of which are also present in the hyperphagic mouse db/db model^32,33^. In contrast, T1D is caused by autoimmune attack against β-cells^34,35^ that is mirrored by the mouse non-obese diabetic (NOD) model, which was also used to show the importance of β-cell stress-induced senescence and senescence-associated secretory phenotype (SASP) in T1D^29,36,37^. β-cell identity can be also disrupted due to chemical stress^38^ and the streptozotocin (STZ) induced ablation of β-cells was previously used to study both T1D and T2D^39–41^. Yet, due to failed clinical translation of treatments showing promise on animal models, it is important to decipher to which extent models resemble human diabetes^34^.

Implication of single-cell RNA sequencing (scRNA-seq) has greatly enhanced our understanding of β-cell maturation, heterogeneity and function in health and disease^2,12,40,42–48^. Nevertheless, there is no consensus on which β-cell populations exist^14,16,49^ and which pathways lead to β-cell dysfunction in different conditions. For example, for T2D progression alone, previous studies used different systems and individually identified various molecular changes, associated with energy metabolism, compensatory insulin secretion, apoptosis, inflammation, dedifferentiation, and disrupted islet communication^43,48,50,51^. This ambiguity can be attributed to heterogeneous cellular states, joint action of multiple molecular mechanisms, different stressors, and confounding of unknown environmental factors^43,47,51–53^. Such complexity cannot be fully captured in datasets of individual studies. Hence, a combined analysis of multiple datasets is needed to comprehensively describe β-cell heterogeneity in health and disease and to disentangle molecular pathways contributing to deterioration of glucose homeostasis in various dysfunction conditions.

Direct comparison of multiple scRNA-seq datasets generated by different scientific groups is often not possible due to batch effects. To circumvent this, multiple scRNA-seq data analysis and integration^54–57^ approaches have been proposed. This also enabled the creation of so-called “integrated atlases” that provide an expertly curated resource with a high-quality embedding optimized to retain biological variation while removing batch effects. Atlases have become an invaluable tool as they provide new insights beyond individual datasets^58–62^, such as description of cellular landscape in health and disease^60,61,63–69^ and comparison across animal or *in vitro* models and corresponding human datasets^70–72^. Whilst previous efforts have been made to compare the results of multiple islet scRNA-seq studies^27,47,73^, a comprehensive integrated atlas of mouse pancreatic islet cells across biological conditions and datasets with sufficient power to identify cell states is still missing. Therefore, we present an integrated mouse islet atlas (MIA) of scRNA-seq datasets across conditions (**Figure 1a**). The analysis of MIA provided insights that could not be obtained from individual datasets (**Figure 1c**), including a holistic description of the β-cell landscape across datasets and conditions, identification of similarities and differences between diabetes models, and disentanglement of molecular pathways involved in different types of β-cell dysfunction (**Figure 1b**). To empower future studies we also made MIA available for both interactive and computational analyses (**Figure 1d**, https://github.com/theislab/mouse_cross-condition_pancreatic_islet_atlas).

**Figure 1:**
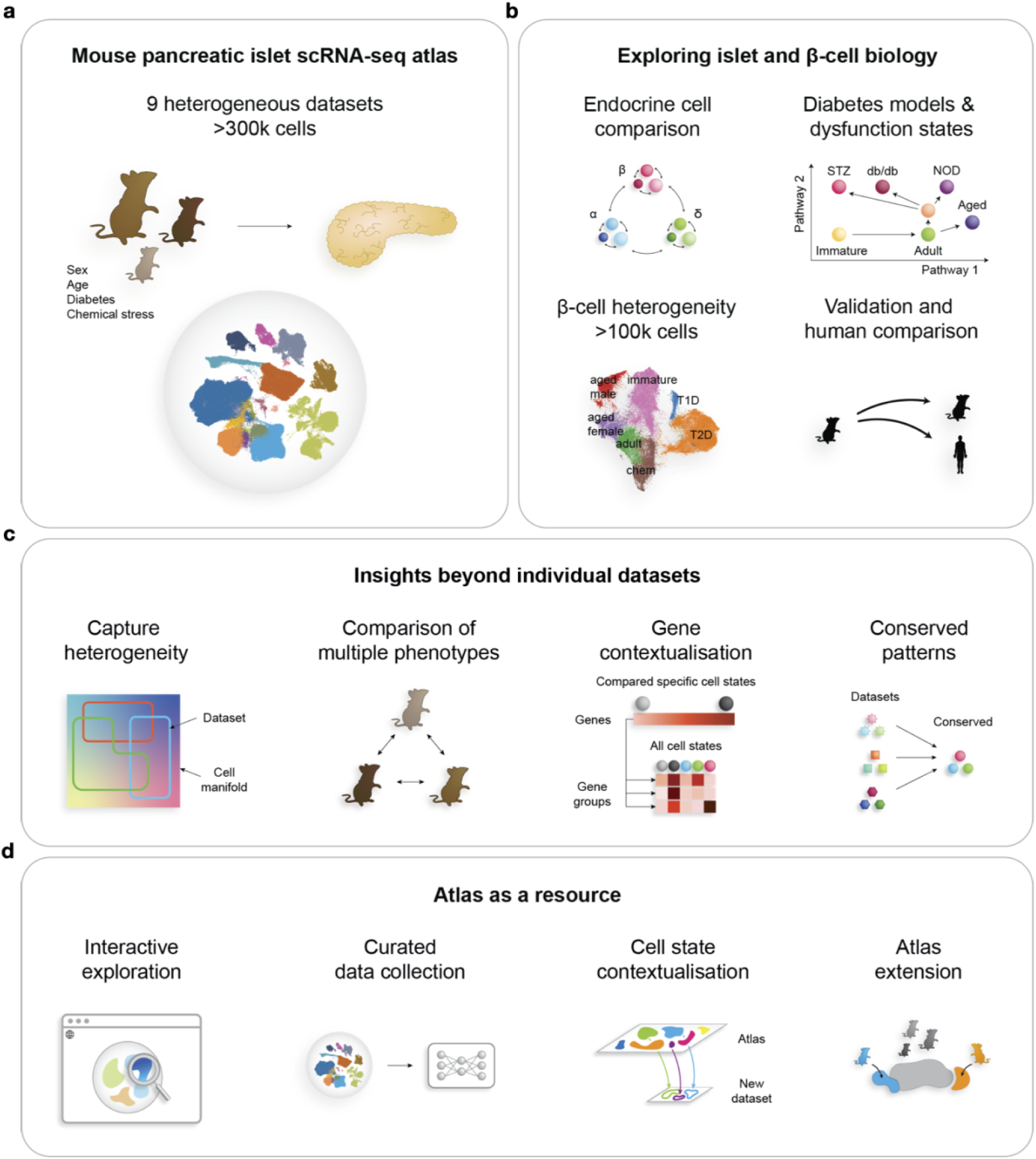
The mouse islet atlas (MIA) of scRNA-seq datasets across conditions offers new insights into islet and β-cell biology. Highlighted are (a) MIA content, including different conditions: sex, age, diabetes models (STZ, db/db, NOD) and anti-diabetic treatments, and chemical stress (application of different chemicals such as FoxO inhibitor), (b) putative novel biological insights, (c) analyses enabled by MIA that would not have been possible on individual datasets, and (d) potential use cases of MIA as a resource for future studies.

## Results and discussion

### An integrated atlas of mouse pancreatic islet cells compiles biological variability across conditions

To better understand how the transcriptome of individual healthy pancreatic islet cells looks like and how it changes across life-time and upon various forms of diabetogenic stress, we integrated nine mouse datasets. We comprehensively collected seven previously published datasets (see **Methods** for data inclusion criteria), as well as generated two new datasets (**Table 1**). MIA contains 301,796 pancreatic islet cells from 56 samples (**Figure 2a,c**, **Table 1** and **Supplementary Table S1**). We use the term dataset for collection of samples that were generated for the same purpose (e.g., published together) and the term sample for jointly processed cells with shared biology, which may originate from a single animal, sequenced individually or demultiplexed, or are pooled across multiple animals sequenced on the same lane without demultiplexing. The samples within MIA vary in sex, age (ranging from embryonic, to postnatal, to adult, to aged), application of chemical stressors implicated in loss of cellular identity (FoxO inhibitor, artemether), and disease status (diabetes models: NOD, db/db, and multiple-low dose STZ (mSTZ) together with different anti-diabetic treatments (vertical sleeve gastrectomy (VSG), insulin, glucagon-like peptide 1 (GLP-1), estrogen) (**Figure 2a**). To cover a wide range of developmental stages we extended the available scRNA-seq data (embryo to adult) with a newly generated scRNA-seq of aged mice (above two years) across sexes (17,361 cells). To identify characteristics of mature cells conserved across datasets we sampled islet cells from adult (four months) male mice (17,353 cells), thus complementing two other publicly available datasets.

**Figure 2:**
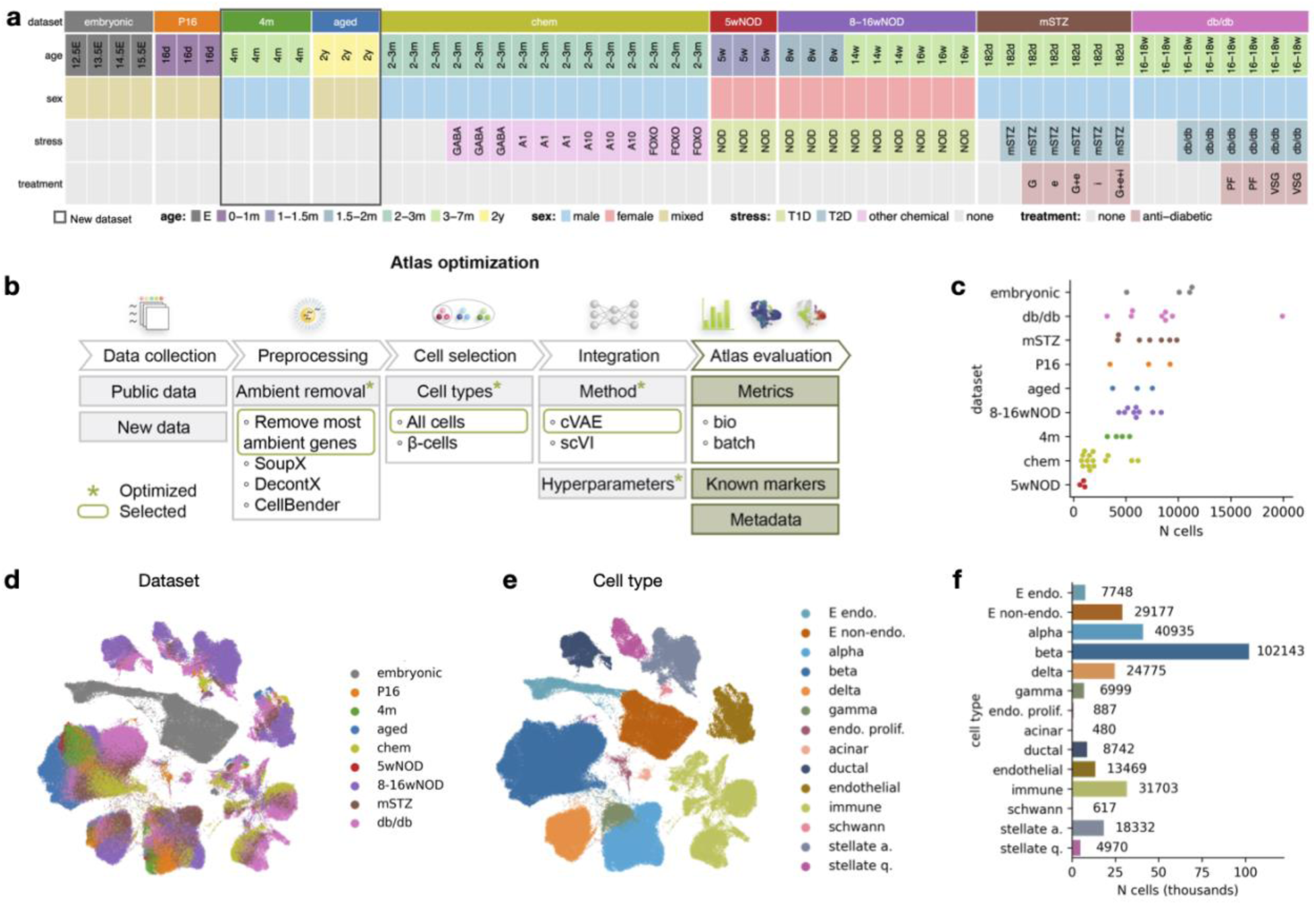
The integrated mouse islet atlas (MIA) captures cell types and states across lifetime, sexes, and multiple stressed or diabetic conditions from different scRNA-seq datasets. (a) Metadata of datasets and samples used in MIA. (b) Overview of atlas integration evaluation. We tested multiple integration approaches and used the circled ones for the final atlas. (c) Number of cells per sample (dots) within each dataset. (d) Dataset distribution within the integrated atlas (excluding low quality cells) shown on a UMAP. Datasets are described in **Table 1**. (e) Atlas-level cell-type re-annotation (excluding low quality cells) shown on a UMAP. (f) Number of cells per cell type from atlas-level re-annotation, excluding low quality cells. Abbreviations: E - embryonic, P - postnatal, d - days, w - weeks, m - months, y - years, A1/A10 - artemether (1 or 10 μM), FOXO - FoxO inhibitor, G - GLP-1, e - estrogen, i - insulin, PF - pair-fed, VSG - vertical sleeve gastrectomy, endo. - endocrine, prolif. - proliferative, stellate a. - stellate activated, stellate q. - stellate quiescent.

**Table 1:** Summary of datasets used for the atlas and their availability.

| Name | Description | N samples | GEO accession | Reference | Source | Ensembl release |
| --- | --- | --- | --- | --- | --- | --- |
| embryonic | Embryo progression from E12.5 to E15.5 | 4 | GSE132188 | <sup>85</sup> | in-house | 100 |
| P16 | Healthy young (P16) islets sorted according to the Fltp lineage tracing model | 3 | GSE161966 | <sup>104</sup> | in-house | 94 |
| 4m | Healthy adult (4m) islets from pancreas head and tail sorted according to the Fltp Venus reporter to isolate FVR+ and FVR- cells | 4 | GSE211796 | previously unpublished | in-house | 94 |
| aged | Healthy aged (2y) islets sorted according to the Fltp lineage tracing model | 3 | GSE211795 | previously unpublished | in-house | 94 |
| mSTZ | Healthy adult control, mSTZ-induced T2D model, and mSTZ model with different anti-T2D treatments | 7 | GSE128565 | <sup>40</sup> | in-house | 100 |
| db/db | Healthy adult control, db/db-induced T2D model, and db/db model with different | 8 | GSE174194 | <sup>32</sup> | in-house | 94 |
|  | T2D treatments |  |  |  |  |  |
| 5wNOD | NOD model of T1D before T1D onset (5w) | 3 | GSE144471 | <sup>178</sup> | external | 100 |
| 8-16wNOD | NOD model of T1D during T1D development (8-16w) | 9 | GSE117770 | <sup>36</sup> | external | 100 |
| chem | Healthy young adult control or with applied chemical stress; sequencing with spike-in cells | 15 | GSE142465 (GSM4228185 - GSM4228199) | <sup>38</sup> | external | 100 |

To enable joint analysis of all datasets we performed data integration, creating a joint embedding space. We ensured optimal trade-off between batch correction and biological preservation on the level of cell types and cell states by evaluating different integration approaches, including preprocessing and data selection, integration tools, and hyperparameter selection (**Figure 2b**), as discussed in **Supplementary Note S1**. The integrated atlas, as depicted in **Figure 2e**, shows clear separation into clusters that correspond to distinct cell types (**Figure 2e**, **Supplementary Figure S2a,b,c**) that co-localize across datasets (**Figure 2d**).

As the available cell type annotation was incomplete and inconsistent across datasets (**Supplementary Figure S2c,d**) we manually re-annotated the integrated embedding (**Figure 2e,f**, **Supplementary Figure S2a**). This enabled us to resolve cell populations that were not annotated in some of the original studies, potentially because low cell numbers hamper annotation^60^. For example, we found that Schwann cells (617 out of 301,796 atlas cells) were present across the studies (**Supplementary Figure S3**), although they were not annotated in any individual dataset (**Supplementary Figure S2d**). Similarly, none of the original annotations distinguished between activated and quiescent stellate cells and some of the studies did not annotate stellate cells at all (**Supplementary Figure S2d**, **Supplementary Figure S3**).

Additionally, we also observed populations influenced by technical artifacts that co-localized across datasets, namely a low quality cluster (lowQ: 853 cells, as well as low quality cells identified based on more detailed analysis of individual cell type clusters: 2782 cells within β-cell cluster and 377 cells within α-cell cluster) and mixed (doublet) clusters (altogether 9966 cells) (**Supplementary Figure S2a, Supplementary Table S2**). They may be useful in the future in automatic annotation transfer to identify residual low quality populations in new datasets, such as doublets that are often hard to identify.

#### Endocrine cell type markers partially overlap in embryonic and postnatal stages

Pancreatic islet profiling and stem cell differentiation highly depend on reliable endocrine cell type markers^74^. However, markers of individual cell types may differ across developmental stages. For example, in embryonic and postnatal stages different cell types are present, meaning that different markers will be specific for an individual cell type against all other present cell types. Furthermore, our integrated embedding revealed molecularly distinct cells states within cell types across development (**Figure 2d, Supplementary Figure S2**). Thus, we provide cell type specific markers separately for embryonic and postnatal mice (**Supplementary Table S3**). We did not compute postnatal ε-cell and embryonic γ-cell markers due to the lack of these cell types at the respective stages.

The newly identified embryonic and postnatal markers only partially overlapped (**Supplementary Figure S4a**), confirming that distinct marker sets are needed at different developmental stages. For example, while the expression of *Cer1* is higher in embryonic compared to postnatal δ-cells, it is a potential δ-cell marker only in postnatal and not in embryonic samples. This is due to the high expression of *Cer1* also in ε-cells and high-level *Ngn3*-expressing endocrine precursor cells (**Supplementary Figure S4b**) that are present only in the embryo (**Supplementary Figure S4b**).

Some of the markers were shared with human endocrine markers reported in a recent scRNA-seq metaanalysis^74^ (mouse homologues *Ttr, Gcg*, *Irx2*, and *Slc7a2* for α-cells; *Ins1*, *Ins2*, *G6pc2*, and *Iapp* for β-cells; *Sst* and *Rbp4* for δ-cell; *Ppy* for γ-cell; **Figure 3a**) and in other publications (*Ghrl* and *Irs4* for ε-cells)^75,76^. Furthermore, we detected several novel cell type-specific genes at different developmental stages (e.g., *Wnk3* and *Nxph1* for α-cells; *Cytip* and *Spock2* for β-cells; *Slc2a3*, *Nrsn1*, and *Spock3* for δ-cells; *Vsig1* for γ-cells; **Figure 3a**). Among these, *Spock3* has been reported multiple times as a human α-cell, rather than δ-cell, marker^74,77,78^. However, in mice we observed consistent upregulation in δ-cells across datasets, which is further supported by a prior study reporting this gene as a δ-cell marker in zebrafish^79^.

**Figure 3:**
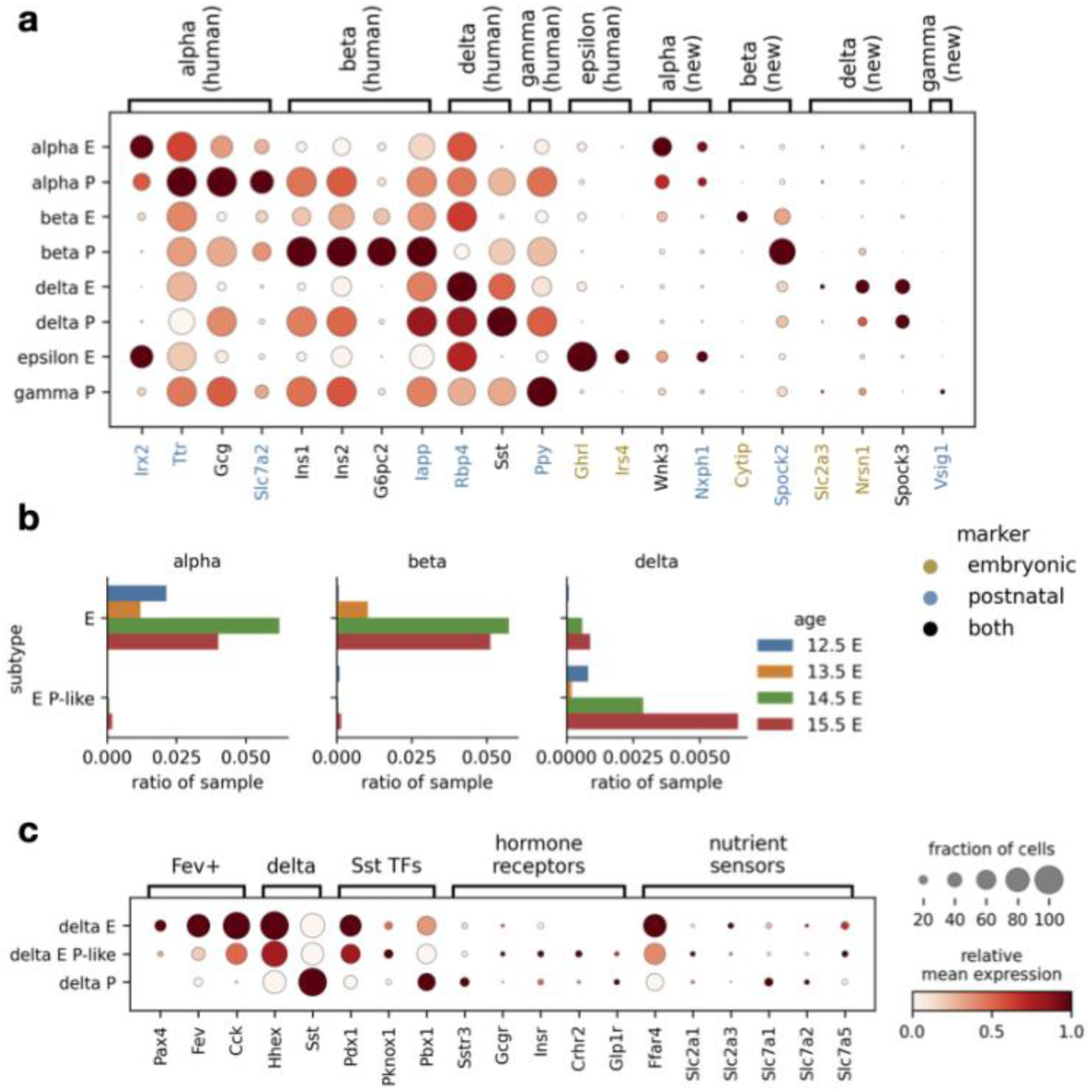
The integrated atlas embedding shows differences between embryonic and postnatal endocrine cells. (a) Expression of endocrine markers shown across postnatal (P) and embryonic (E) endocrine cell types, including known markers shared with human (human) and newly identified markers (new). (b) Number of cells in each embryonic endocrine cell group within individual embryonic samples, expressed as a fraction of cells within a sample. Cell groups: E - embryonic cells mapping to the embryonic cluster, E P-like - embryonic cells mapping to the postnatal cluster. (c) Expression of known maturity and δ-cell function markers across embryonic δ-cells groups. Groups are as in panel b; P - postnatal cells mapping to the postnatal cluster. In the panels a and c relative expression is computed as the average of cell groups normalized to [0,1] for each gene feature.

We analyzed protein expression of two transcriptome-based markers (*Ttr* in α-cells and *Rbp4* in δ-cells) with immunohistochemistry in mouse islets (**Supplementary Figure S4c**). As anticipated, expression of Ttr protein, which is involved in regulation of Gcg expression and glucose homeostasis^80^, was specific to α-cells. In contrast, Rbp4 protein, which was previously reported to be a marker of δ-cells^74,81^, is expressed across the whole islet and could thus not be used to reliably distinguish δ-cells in immunohistochemistry (**Figure 3a, Supplementary Figure S4c**). Its relatively high protein levels in β-cells may be further explained by the young developmental stage (P9) of the used islets and hence β-cell immaturity, which is known to be associated with high Rbp4 expression^82,83^.

#### Embryonic δ-cells cluster with postnatal δ-cells

One of the key questions in islet biology is when and how endocrine cells become functionally mature, which is of relevance for developing functional cell types from pluripotent stem cells^11^. As MIA provides a shared embedding of different biological conditions from multiple datasets that would otherwise not have been comparable due to confounding batch effects, we leveraged it to analyze cell populations during endocrine maturation. As expected, most embryonic cells (termed E group) generally did not overlap with postnatal cells (termed P group), but surprisingly we observed that a large proportion of embryonic δ-cells mapped to the postnatal δ-cell cluster (termed E P-like group, **Figure 3b**, **Supplementary Figure S2d**).

To understand this overlap, we evaluated the expression of endocrine development and δ-cell function related genes. The E P-like δ-cells had, in comparison to the E group, lower expression of δ-cell lineage determinant *Hhex^84^* and lower expression of gene markers enriched in Fev-positive population^85^, from which δ-cells arise^85–88^ (**Figure 3c**, **Supplementary Figure S5a**). Among known δ-cell functional genes, somatostatin was highly expressed already in the E group, likely because *Sst* has been used for δ-cell annotation, therefore not capturing earlier δ-cell developmental stages^75^. Other functional genes include transcription factors involved in *Sst* expression^89^ and sensors required for appropriate paracrine regulation, namely neurotransmitters, hormone receptors, including the somatostatin receptor (*Sstr3*) (autocrine feedback), and nutrient sensors, including sensors for milk-based high-fat weaning diet (fatty acids: *Ffar4*; amino acids: SLC7 family)^81,90–94^ (**Figure 3c**). They were relatively higher expressed in all cell groups. This indicates that δ-cells already possess the machinery for regulating somatostatin expression at the embryonic stage and that they quickly downregulate expression of developmental genes, explaining the mapping of embryonic δ-cells to the postnatal cluster. However, we must note that genes potentially involved in somatostatin regulation could also be related to other cellular functions at this developmental stage. Thus, further validation of δ-cell physiology during development would be required.

### β-cells show heterogeneity across and within conditions compiled within the atlas

Extensive research has shown that β-cells are heterogeneous^15,17,19^, however, there is a lack of knowledge on how these states relate^14,16^. Hence, we aimed to use MIA to comprehensively describe β-cell states alongside their molecular characteristics in different sexes, ages, and stress conditions (**Table 1**).

To test whether the integration is adequate for downstream analyses of β-cell states we assessed a MIA subset consisting of 102,143 β-cells. Cells separated based on biological covariates, such as age and disease status, and overlapped between samples with similar biological covariates from different datasets (**Figure 4a**, **Supplementary Figure S6**). For example, healthy control β-cells mapped together regardless of their dataset of origin (mSTZ, db/db, and 8-16wNOD), while the cells from diabetic samples from these datasets mapped away from the healthy clusters. This is in accordance with previously reported β-cell changes in aging and diabetic dysfunction^14,45,95,96^. Furthermore, we assessed the expression patterns of known immaturity (*Rbp4*), maturity (*Mafa*), stress (*Gast*), aging/senescence (*Cdkn2a*), and inflammatory (*B2m*) β-cell markers (**Figure 4b**), showing complementary patterns when considering opposite activity of β-cell functional maturation (*Mafa*) and dedifferentiation (*Gast*) markers. Altogether, this indicates successful integration of the multiple data sets both on the cell-type and cell-state level.

**Figure 4:**
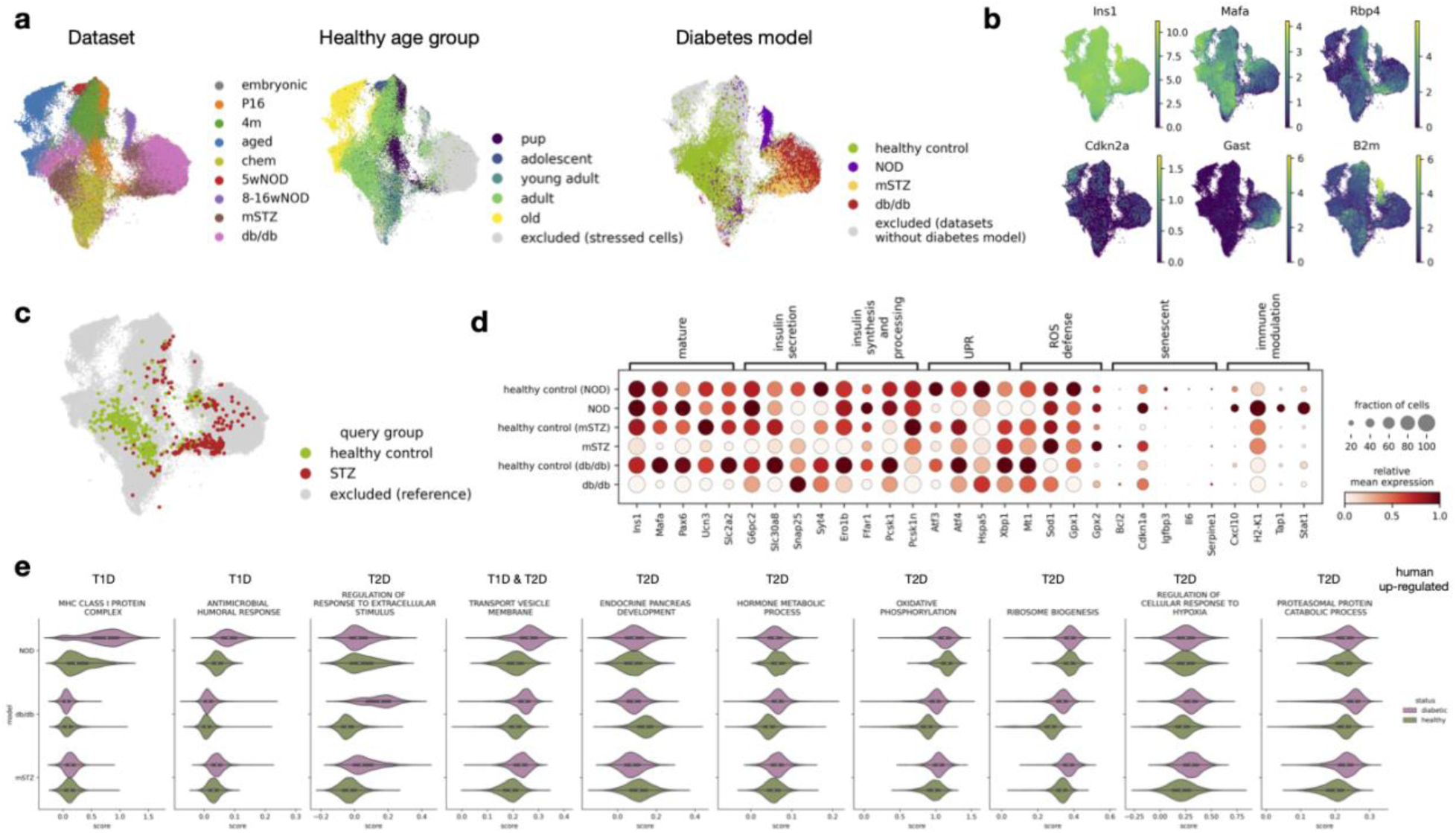
The integrated atlas embedding reveals similarities between mSTZ and db/db diabetes models. (a) Distribution of technical (dataset) and biological (age, disease status) covariates on a UMAP of the β-cell MIA subset. The age subplot shows only cells from healthy, non-stressed samples. The disease subplot shows only cells from samples belonging to datasets that contain both healthy and diabetes model data. (b) Expression of selected β-cell heterogeneity markers on a UMAP of the β-cell MIA subset. (c) Joint UMAP embedding of the reference atlas (background) and the external (Feng) mouse dataset (query, foreground) indicating positioning of healthy control and STZ treated query cells. (d) Expression of known β-cell function genes across different diabetes models and corresponding healthy controls from individual datasets (the NOD model is from the 8-16wNOD dataset, other model names correspond to the dataset names). Relative expression is computed as the average of cell groups normalized to [0,1] for each gene feature. (e) Activity of gene sets up-regulated in T1D or T2D human samples shown for mouse diabetes models and corresponding healthy controls from individual datasets (as in d).

### Comparison of diabetes models identifies transcriptomic similarities in db/db and mSTZ diabetic β-cells

The usage of the appropriate mouse model is of utmost importance to studying β-cell function both in health and disease conditions^28^. Different models with unique phenotypes and disease mechanisms have been developed^29^, each of them with advantages and limitations to be considered^28^. In order to better understand the transcriptomic differences among the diabetes mouse models, we compared the commonly used genetic models of T1D (NOD, for which we used samples from early disease stages^29,36^) and T2D (db/db^33^) together with the β-cell ablation model (STZ) that was previously used to study both T1D and T2D^39,40^. The NOD model is characterized by autoimmune and cytokine-mediated destruction of β-cells as well as ER stress^97,98^. The Leptin-receptor-deficient db/db mice are obese, hyperglycemic, and dyslipidemic^99,100^, leading to β-cell failure and compensation, which are associated with metabolic stress, including ER stress^32,33^. The STZ treatment is used for specific destruction of β-cells due to its affinity for the Glut2^101^ expressed in β-cells. The stressor is applied either in a single high dose to resemble T1D or in multiple low doses to elicit partial β-cell loss reminiscent of T2D, but in the absence of insulin resistance^28^, with both strategies analyzed below.

Based on MIA embedding we found that β-cells from mSTZ-induced (multiple low doses) and db/db models mapped together, separately from NOD diabetic β-cells (**Figure 4a**). To further validate the similarity between the mSTZ and db/db models, we mapped onto MIA another mouse dataset (referred to as Feng dataset^41^, not part of MIA), containing samples treated with STZ (single high dose). Again, the healthy control cells from the Feng study mapped onto the healthy β-cell region of MIA, and STZ-treated cells mapped onto the region with mSTZ and db/db model samples (**Figure 4c**). Similarly, in the future mapping onto MIA may reveal relationships between other dysfunctional conditions.

To better understand molecular mechanisms underlying β-cell dysfunction within each of the models, we analyzed the expression of known β-cell function and stress genes (**Figure 4d**). In the mSTZ and db/db models multiple maturity and insulin-related genes were downregulated, whilst in the NOD model immune modulation genes were upregulated. In all three models we observed expression changes in several unfolded protein response (UPR), reactive oxygen species (ROS) defense, and senescence-related genes. This indicates the involvement of metabolic stress in db/db and mSTZ models and immune stress in the NOD model, in accordance with current views on T1D and T2D pathomechanisms^102^.

To elucidate which mouse models capture transcriptional signatures of human T1D or T2D, we assessed whether changes observed in human diabetes are also present in mice. We performed differential gene expression (DGE) analysis on β-cells from multiple human T1D and T2D datasets (**Table 2**), selected genes upregulated across multiple datasets per diabetes type (T1D 32 genes, T2D 59 genes), and identified enriched gene sets (**Supplementary Table S4**). We further complemented our list of newly identified gene sets with known human diabetes-associated gene sets from literature. Human T1D is marked by upregulation of immune gene sets^30^, which were much more strongly upregulated in NOD than db/db and mSTZ models (**Figure 4e**, for details of gene set activity analysis across mouse models see **Supplementary Note S2**). Conversely, human T2D is associated with changes in hormone metabolism and stress related to metabolic compensation^30,31,103^, which were upregulated in db/db and mSTZ but not in the NOD model. Thus, the mSTZ model reflects key molecular changes of human T2D, but not T1D. The presence of metabolic stress in the mSTZ model β-cells after clearance of the chemical stressor can be explained by the surviving population of β-cells being too small to prevent hyperglycemia and hence leading to compensatory insulin-production behavior and subsequent stress.

**Table 2:**
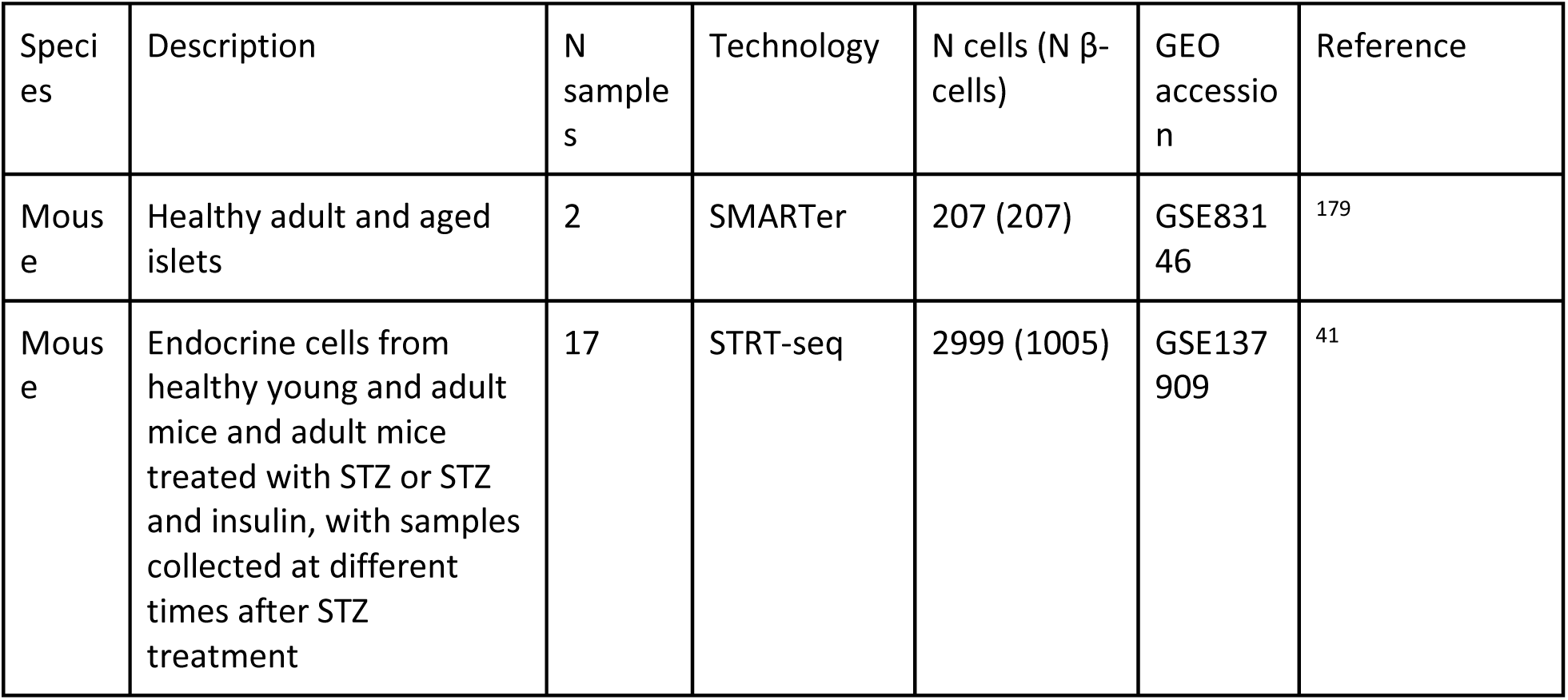
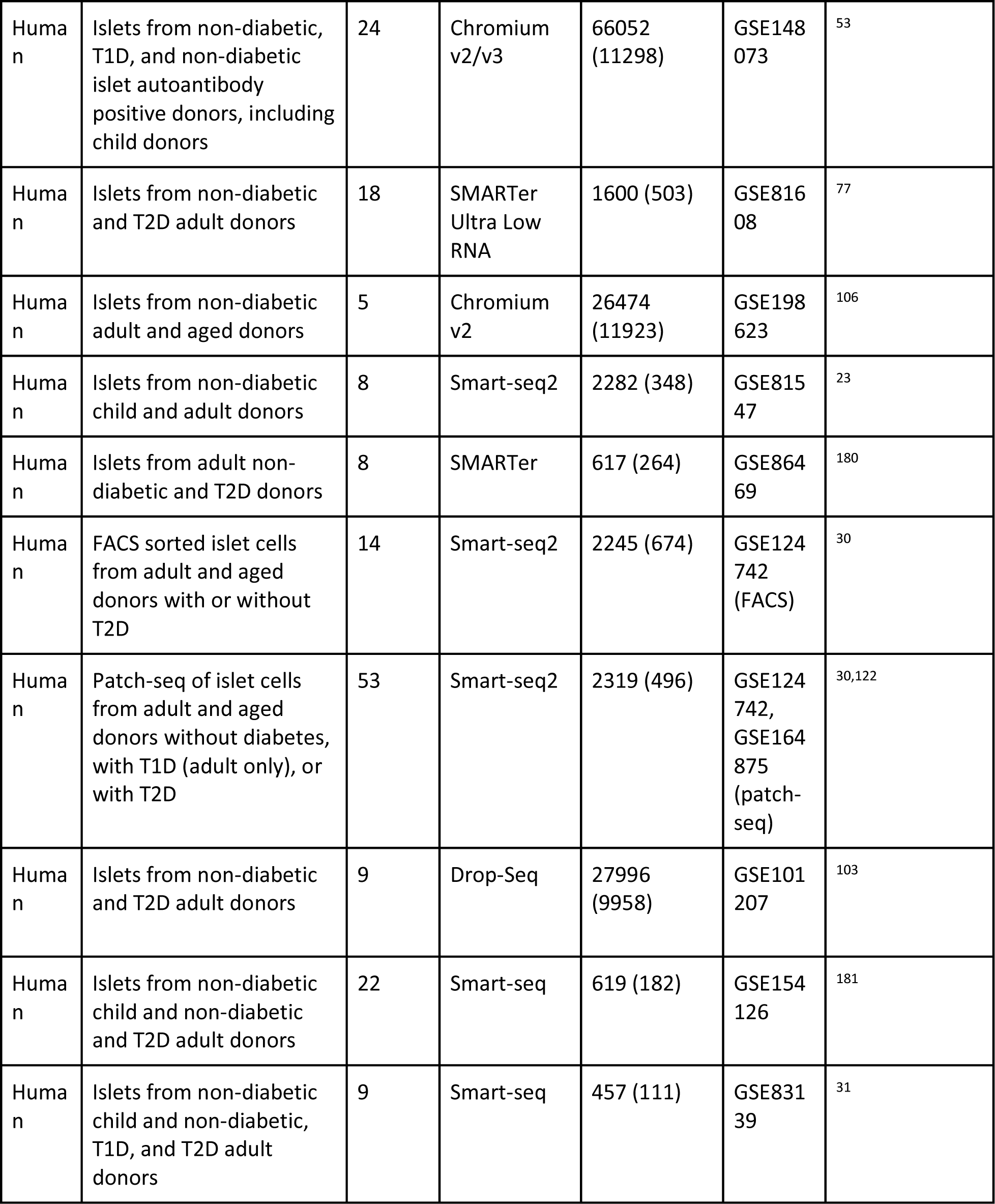
Datasets used for validation; not part of the atlas.

### Markers of β-cell states conserved across datasets

As it is unclear how newly reported β-cell states correspond across publications^14,15^ we next aimed to utilize the cross-dataset integrated conditions within MIA to describe β-cell heterogeneity in health and disease in an unified manner. We annotated states on postnatal non-proliferative β-cells (“beta” cluster in **Figure 2e**) and labeled them on the basis of the metadata (altogether referred to as “coarse states”, **Figure 5a**, **Supplementary Figure S7a**). We resolved populations of healthy adult, immature, aged (separated by sex), NOD diabetes model, mixed db/db and mSTZ diabetes models, and cells from the dataset with chemical perturbations in cultured islets (referred to as chem), that likely separate due to strong differences in sample handling. For detailed description of states see **Supplementary Note S4**.

**Figure 5:**
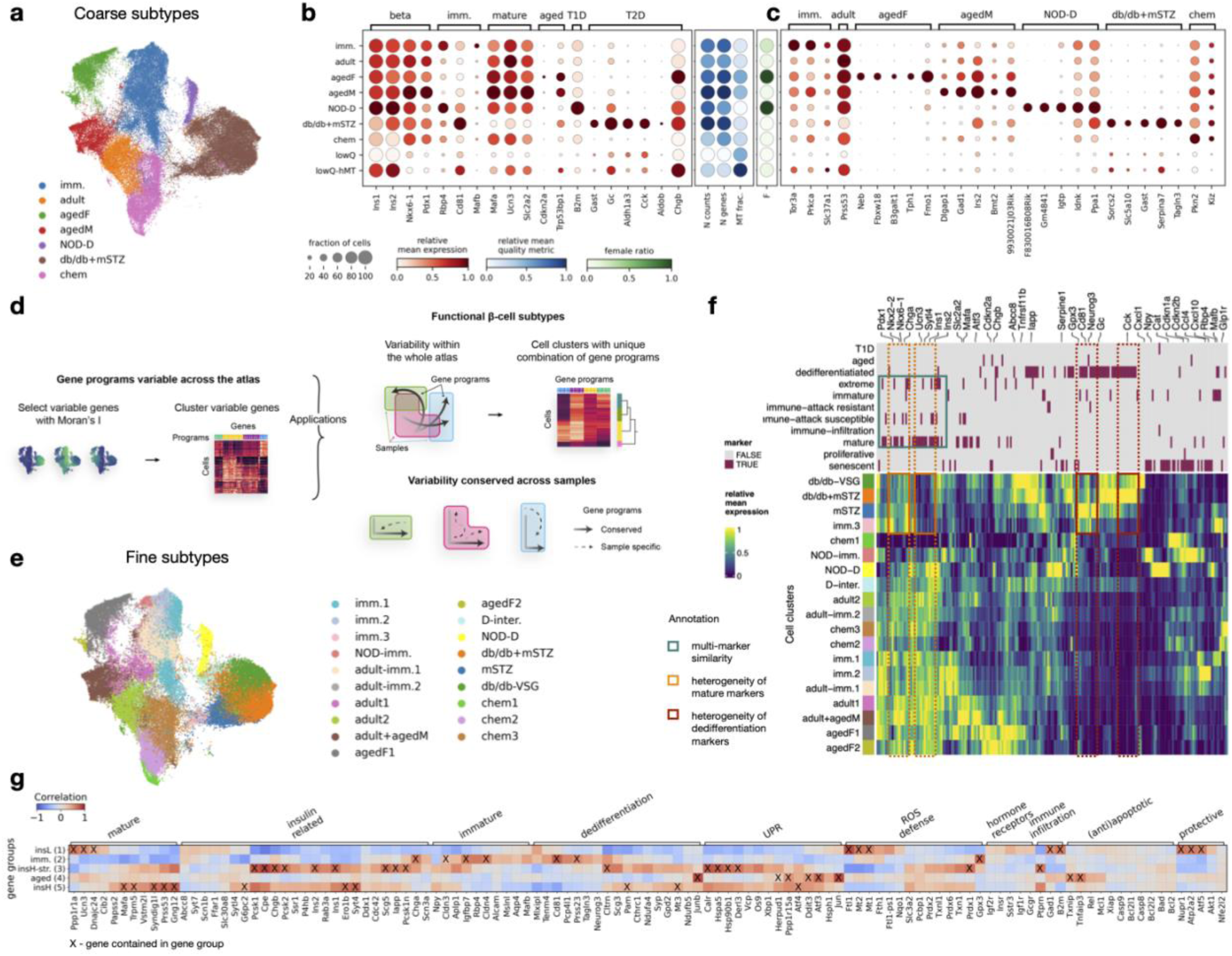
MIA encompasses β-cells heterogeneity across and within biological conditions. (a) Coarse β-cell states labeled based on sample metadata (excluding low quality clusters) shown as a UMAP. (b) Expression of known markers (marker groups are specified on the top of the plot), quality control metrics, and sex ratios across coarse β-cell states displayed in separate dotplot panels. In the marker expression panel the dot size indicates the fraction of cells expressing a gene, while in other panels it is set to a fixed size. (c) Expression of newly defined markers of coarse β-cell states. (d) Overview of the method used for extraction of gene programs (GPs) and subsequent cell clustering resolution selection or definition of consistently variable GPs across samples. (e) Fine β-cell states defined based on the presence of a unique combination of GPs (excluding low quality clusters) shown as a UMAP. (f) Expression of known β-cell heterogeneity markers across fine β-cell states. The upper panel shows phenotypes associated with individual genes. Dotted boxes represent two distinct sets of maturity (orange) and dedifferentiation or diabetes markers (red); the solid cyan box shows overlap and expression similarity between maturity, immune-attack susceptibility, and extreme insulin producer markers. (g) Correlation between gene groups variable in all healthy samples and known β-cell heterogeneity markers on the healthy β-cell subset. Markers present within a specific gene group are annotated with an X. Abbreviations: imm. - immature, M - male, F - female, NOD-D - NOD diabetic, D.-inter - diabetic intermediate, insL/H - insulin low/high, str. - stressed. In the panels b, c, and f relative expression is computed as the average of cell groups normalized to [0,1] for each gene feature.

We support the annotation of coarse states with known β-cell state markers depicted in **Figure 5b**. Some known markers were not state-specific, such as certain immature markers that were also highly expressed in the db/db+mSTZ state (e.g., *Cd81*, **Figure 5b**), in accordance with β-cell dedifferentiation in mouse diabetes models^32,40,104^. Thus, the identification of new state-specific markers could improve monitoring of β-cells in specific states in order to study their function. We identified new markers specific for an individual β-cell state and conserved across all datasets mapping to that state, with top markers highlighted in **Figure 5c** (**Supplementary Table S5**, more detailed description is in **Supplementary Note S4**). For example, we identified a new marker of healthy adult state *Prss53*, associated with mitochondrial function^105^.

To test the robustness of our markers we analyzed their expression on the Feng mouse dataset that is not part of the atlas^41^. This dataset consists of healthy young and adult mice, with multiple samples spanning the ages of 0.1-4 months, as well as STZ-treated diabetic samples (**Supplementary Figure S8c,d**). The proposed T2D-model state (db/db+mSTZ) and adult state markers were expressed as expected in the Feng dataset, however, we did not observe specific expression of immature markers in the young samples. We next evaluated whether this difference arises due to a different immature cell state present in the Feng dataset or due to technical issues in marker identification. Thus, we mapped Feng dataset cells to MIA. Indeed, we observed differences in the two immature cell states, as young samples from the Feng dataset did not map to MIA immature state (**Supplementary Figure S9a,b**). The Feng postnatal day 3 (P3) β-cells mapped between embryonic and postnatal β-cells of MIA and the young postnatal cells (postnatal days 12 (P12) and 21 (P21)) mapped between the immature, adult, and chem MIA states.

Additionally, we assessed whether known and newly identified markers could be directly translated to ten human datasets with differences in donor metadata (**Supplementary Figure S8e,f**). Only *B2m* (T1D marker)^36^ and *Rbp4* (immature marker)^104^ were significantly upregulated in all human samples associated with those phenotypes. This is in accordance with previous reports^106^ showing that not all mouse markers directly translate to human data.

### Increased β-cell clustering resolution reveals heterogeneity within biological conditions

β-cells are known to be heterogeneous within individuals^19,20,107^. However, our metadata-driven coarse states mainly did not reveal multiple populations per sample (**Supplementary Figure S7c**). Some markers were heterogeneously expressed within coarse states, such as *Rbp4* in young and db/db+mSTZ states and *Mafa* and *Gast* in the db/db+mSTZ state (**Figure 4b**), indicating that we could identify higher resolution states in MIA.

Annotation of cell states is challenging due to uncertainty about the number of distinct states^108^. To ensure that states can be always biologically interpreted, we based clustering on interpretable features (termed gene programs (GPs), **Figure 5d** and methods). GPs are data-driven groups of genes coexpressed across β-cells (27 GPs, size=14-228 genes, **Supplementary Figure S10a**, **Supplementary Table S6**). Most of the GPs were enriched for distinct molecular functions (**Supplementary Table S6**) and we show that they generalize to other datasets by explaining variance in two external mouse and ten human datasets (**Supplementary Figure S10f**).

We defined 19 fine β-cell states (**Figure 5e**), which mainly corresponded to subclusters of the coarse states (**Supplementary Figure S7e**) and described more subpopulations within samples, while still containing cells from multiple samples and datasets (**Supplementary Figure S7d, Supplementary Table S2**). Additionally, two clusters were characterized by low quality control metrics and were thus not regarded as true cell states (**Figure 5e**, **Supplementary Figure S7b**). We further discuss β-cell heterogeneity captured within MIA in relation to previous literature in **Supplementary Note S5**.

We observed two populations of β-cells in the mSTZ model (states mSTZ and db/db+mSTZ; **Figure 5e**, **Supplementary Figure S7d**). We used biologically interpretable GP differences to ease the comparison of these two states (**Supplementary Figure S10b,d**, for validation of this approach see **Supplementary Note S6**). The db/db+mSTZ state had higher activity of multiple GPs that contained known diabetes markers or were associated with ER stress (GP2, GP3, GP4) and cell state mSTZ had higher activity of GPs associated with immaturity (GP8, GP23). Both increased ER stress and immaturity were reported in the paper publishing the mSTZ dataset^40^, however they did not describe dysfunctional populations differing in the two processes. While the more immature state (mSTZ state) was specific for the mSTZ model, the more stressed state (db/db+mSTZ state) also contained db/db model cells. This may be explained by either mSTZ diabetes model having a milder hyperglycemia than the db/db model^32,40^, leading to a lower β-cell compensatory response and thus reduced stress, or by a different mechanism of β-cell damage due to the use of STZ. As these two populations clearly differ in their metabolism, they may be of relevance for studying diabetes with the mSTZ model.

Publications based on individual datasets often do not agree on β-cell heterogeneity markers^47^. Thus, we used the wide range of β-cell phenotypes across dataset within MIA, encompassed by the fine β-cell states, to assess population markers manually extracted from the literature (**Figure 5f**, **Supplementary Table S7**). Some markers previously reported as marking the same β-cell population, such as markers of maturity or dedifferentiation (often related to T2D models), separated into multiple groups with distinct expression patterns across fine states (**Figure 5f**). This shows how MIA could be used to find specific and sensitive markers. Furthermore, we observed that different groups of markers reported across studies with different biological focuses share similar expression profiles, such as mature^18,32,109,110^, extreme insulin producing^32,110^, and immune-attack susceptible markers^111^. The immune-attack susceptible markers were extracted from Rui et al. (2017) who reported NOD subpopulations differing in immune attack susceptibility. They reported that the immune-attack susceptible population expressed β-cell maturity genes and indeed we observed that the population markers reported by Rui et al. (2017) co-localized with known maturity genes in MIA (**Figure 5f**). This demonstrates how the heterogeneous cell states within MIA can be used for gene contextualization by providing information on which β-cell states express a gene of interest and which known markers have similar expression patterns.

### β-cell dysfunction and aging associated heterogeneity is observed across healthy samples

In our GP analysis we observed that GPs that changed between healthy and T2D-model cells (GPs: 3, 4, 19, 20, **Supplementary Figure S10a,b**) were also among GPs explaining the largest proportion of cell-to-cell variability within healthy datasets and samples in both mouse and human (**Supplementary Figure S10g, Supplementary Table S6**). This motivated us to describe heterogeneity conserved across healthy adult samples.

We collected genes that are consistently variable within individual healthy samples and grouped them based on co-expression patterns conserved across samples, resulting in five gene groups (detailed description of groups is in **Supplementary Note S7**, **Supplementary Table S8**). Groups 3 and 5 were associated with β-cell maturity and insulin production, with group 3 having a stronger insulin-production related stress signature (**Figure 5g, Supplementary Table S8**). Group 1 contained genes implicated in β-cell metabolic stress recovery, such as ATP production related genes^107^ (**Figure 5g, Supplementary Table S8**). The negative correlation between expression of group 1 and groups 3/5 is in accordance with previously reported cycling of β-cells between insulin production and recovery in mouse and human^107,112,113^. As group 1 genes, including multiple mitochondrial genes, β-cell maturation and function genes (*Ucn3*, *Ftl1*, *Cd63*, Scg2)^73,114^, and protective genes (*Nupr1*, *Atp2a2*, *Atf5*)^115–117^, are involved in healthy metabolic stress recovery they may be of interest for T2D therapy. Indeed, group 1 showed the lowest activity in the diabetes model β-cells (**Supplementary Figure S11b**), indicating impaired stress recovery.

We also observed two gene groups indicating that cells within healthy adults differ in the degree of maturity and senescence. Group 4 contained senescence genes and healthy adult cells most highly expressing these genes co-localized with aged cells. Interestingly, while group 2 contained immaturity genes the healthy adult cells with high expression of this group partially co-localized with the immature subset of mSTZ-model cells (fine β-cell states imm.3 and mSTZ) (**Supplementary Note S7**, **Figure 5g, Supplementary Figure S11**, **Supplementary Table S8**).

Comparison to a meta-analysis of human healthy heterogeneity markers^47^, revealed shared genes *Tm4sf4* and *Clu* from group 3 (insulin production and metabolic stress) and genes *Fos*, *Herpud1*, and *Rgs4* from group 4 (aging). While these orthologues likely share function across species, Mawla and Huising (2019) did not specifically state which β-cell states they are associated with.

### Diabetes response of β-cells is highly complex

While β-cells are the primary cell type affected in diabetes, the disease also has broader effects on the whole islet^118,119^. To investigate these effects, we performed DGE analysis between healthy and T1D model or T2D model samples in α-, β-, γ-, and δ-cells. All cell types had a large number of differentially expressed genes (DEG) in both diabetes types (**Supplementary Figure S12**, **Supplementary Table S9**). DEGs in β-cell T1D model and T2D model had a relatively low overlap and were also distinct from DEGs in other cell types (**Figure 6b**). This is in accordance with different mechanisms that lead to loss or dysfunction of β-cells in T1D and T2D^102^. In contrast, DEGs overlapped more strongly between T1D model and T2D model within α-, γ-, and δ-cells and also showed relatively high overlap across these cell types. This is likely due to β-cells being the primary cell type affected in diabetes, further leading to islet disruption and causing residual stress in other endocrine cells^120,121^.

**Figure 6:**
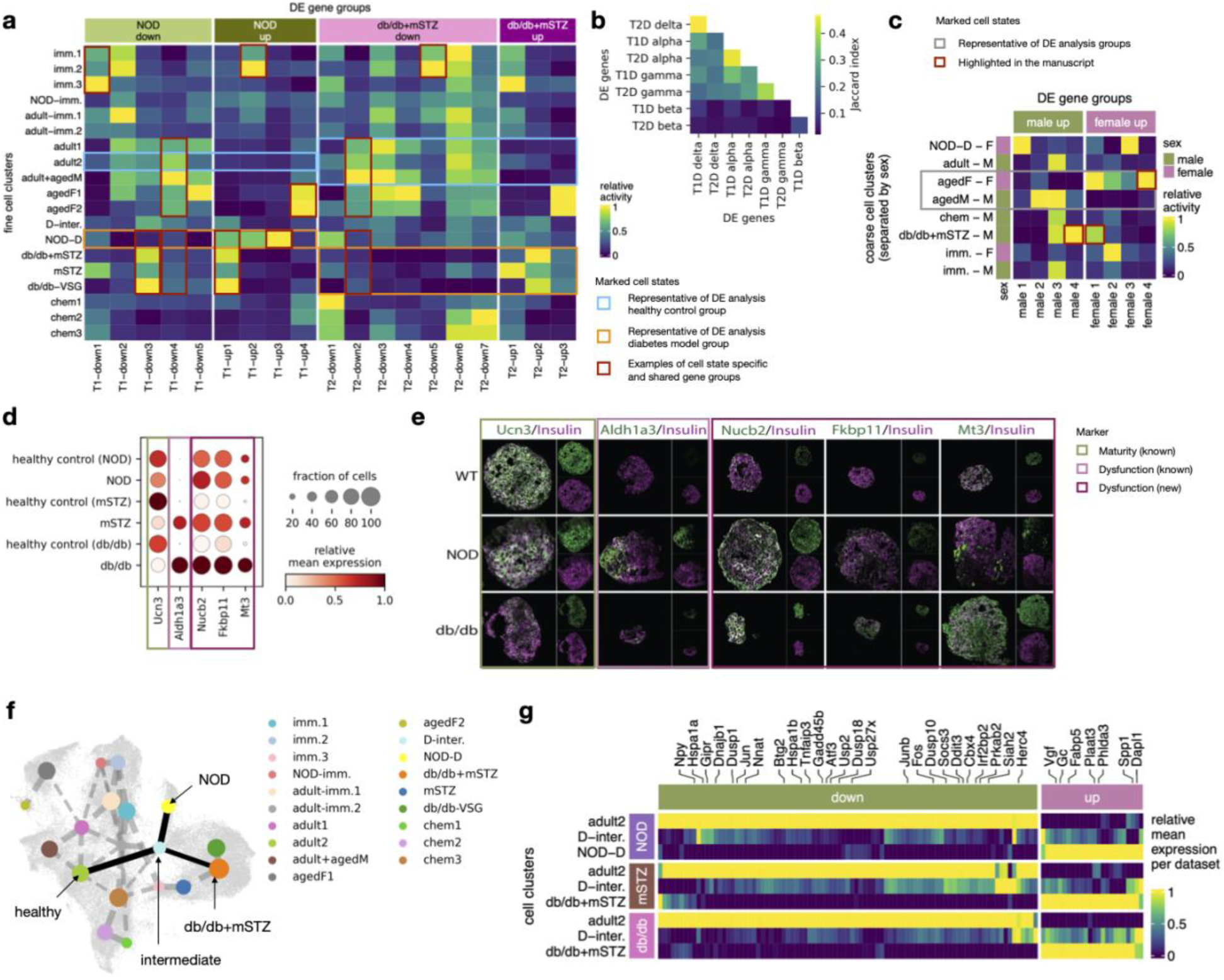
β-cell diabetes dysfunction involves different molecular patterns that are unique or shared with other conditions, including different diabetes models and aging. (a) Activity of β-cell diabetes-trajectories (NOD and db/db+mSTZ) associated DEG groups across fine β-cell states shows diabetes specific and cross-condition shared gene groups (examples are marked with red rectangles). Cell groups that are representative of healthy and diabetic states within the DGE analysis trajectory are marked with rectangles (blue and orange, respectively). (b) Overlap of DEGs across diabetes model types (T1D - NOD, T2D - db/db+mSTZ) and endocrine cell types. (c) Expression of DEG groups between aged males and females across coarse β-cell states, split up by sex. Marked are cell groups discussed in more detail in the text as well as cell groups that are representative of healthy and diabetes model cells used in the DGE analysis. (d) Gene expression of diabetes markers that were validated on protein level. Expression is shown for diabetes models and controls from corresponding datasets. (e) Validation of selected diabetes model β-cell DEGs on protein level with quantification of fluorescence intensity in antibody staining. (f) PAGA graph showing connectivity (lines) between fine β-cell states (dots) imposed on top of the β-cell UMAP. The connections between healthy, intermediate, and diabetes model states are marked in solid lines. (g) Same-direction DEGs across the NOD and db/db+mSTZ diabetes model trajectories already change expression from healthy to intermediate state, with the changes further increasing from healthy to diseased states. The heatmap shows expression across healthy, intermediate, and diabetes model states for each dataset (dataset 8-16wNOD is abbreviated as NOD). Expression is normalized per gene and dataset. Abbreviations: imm. - immature, M - male, F - female, NOD-D - NOD diabetic, D.-inter - diabetic intermediate. In the panels a, c, d, and g relative expression is computed as the average of cell groups normalized to [0,1] for each gene feature.

To characterize the residual stress within endocrine cell types other than β-cells we examined shared DEGs in both diabetes types. Upregulated genes were enriched for ER stress, whilst downregulated genes were enriched for gene sets related to membrane depolarization and ion transport (**Supplementary Table S9**) and contained hormone genes (*Gcg* in α-cells, *Ppy* in γ, *Sst* in δ) (**Supplementary Table S9**). This indicates that diabetes also affects endocrine hormone production and secretion in endocrine cell types beyond β-cells. In support of this, a recent human α-cell patch-seq study reported a loss of electrophysiological identity in T2D^122^ and electrophysiology of δ-cells was likewise reported to be disrupted in prediabetic mice^123^. However, in further analyses we decided to focus on β-cells due to their importance in diabetes development^119^.

#### Diabetes signatures in β-cells are comprised of both unique and cross-condition dysfunction genes

To find genes dysregulated in the T1D NOD model and T2D db/db and mSTZ model β-cells, a DGE analysis was performed for each model group. As cells within individual subjects can be heterogeneously dysfunctional, leading to reduced power in DGE analysis^103^, we leveraged MIA embedding to assign cells from healthy controls and disease models along a healthy-dysfunctional trajectory (**Supplementary Figure S13a**, **Supplementary Note S8**). This is of special importance for NOD mice, as in the original study the authors observed incomplete penetrance ^36^dysfunctional β-cell phenotype ^36^.

As the DGE analysis resulted in hundreds of DEGs that are expected to be heterogeneous in terms of their molecular function, we clustered them using their expression across all β-cells within MIA (sizes 12-349 genes, **Figure 6a**, **Supplementary Table S10**). The groups are described in more detail in **Supplementary Table S10** in terms of gene set enrichment, gene membership, and cell states with high expression. In the text they are referred to as T1 groups for NOD and T2 groups for db/db+mSTZ.

First of all, we used the DEG groups to disentangle dysfunction patterns of interest from confounding effects. In the original NOD dataset study by Thompson et al. (2019) the authors observed confounding of dysfunction progression and age differences between samples containing healthy (8 weeks) and dysfunctional cells (14 and 16 weeks), impairing the interpretation of diabetes associated changes. Indeed, we also observed, among NOD downregulated genes, one group (T1-down1), which was highly expressed across multiple immature states (**Figure 6a**) and contained genes associated with immaturity (*Pyy*, *Npy*)^124,125^ thus likely representing a confounding effect of age. Other gene groups did not seem to be associated with known batch effects.

With our DEG clustering approach we disentangled two NOD-upregulated immune processes (groups T1-up2 and T1-up3) that showed differences in expression across β-cell states. Group T1-up3 was NOD diabetic cells (state 14-16wNOD) specific and more strongly enriched for antigen processing genes (containing genes *B2m*, *Tap2*, and histocompatibility II group members), while T1-up2 was, in addition to NOD diabetic cells, also highly expressed in immature cells (**Figure 6a**) and more strongly enriched for innate immune response genes (containing genes *Stat1*, *Stat2*, *Gbp7*, and immunoproteasome group members), potentially representing the regulation of β-cells by the immune system that is not restricted to diabetes^126^. Upregulation of both T1-up3 and T1-up2 in NOD is in accordance with active involvement of β-cells in T1D-related immune response by means of antigen presentation and immune infiltration in the islets^36,127^, respectively. Furthermore, in the NOD diabetes model we also observed upregulation of senescence related genes (group T1-up4) that were shared with aged females (**Figure 6a**). Indeed, senescence genes have been previously reported in association with NOD model dysfunction and aging individually^36,128^ and we here show their relationship.

As expected, in db/db+mSTZ cellular metabolism that is necessary for normal β-cell function^102^ was disrupted. A group of genes (T2-down3) was downregulated across all T2D model cell states and was higher across healthy cell states (**Figure 6a**), with enrichment for insulin secretion and steroid metabolism. Additionally, we observed groups supporting mSTZ subpopulations associated with immaturity or metabolic stress, which we observed above based on GP differences (see **Supplementary Note S9**).

Multiple parallels can be drawn between NOD and db/db+mSTZ dysregulation. For example, NOD group T1-up1 also showed high expression in cell states from db/db and mSTZ datasets (**Figure 6a**) and partially overlapped with db/db+mSTZ upregulated genes (**Supplementary Figure S13d**), with the overlap containing multiple genes previously associated with diabetes (Gc, Fabp5, Spp1, and Vgf)^129–132^. NOD and db/db+mSTZ also shared similarities in downregulated genes (T1-down4 and T2-down2, **Supplementary Figure S13d**) that were in turn highly expressed in healthy mature cells (**Figure 6b**). These groups contained multiple cross-species conserved β-cell genes (*Atf3*, *Btg2*, *Ddit3*, *Egr4*, *Fosb*, and *Jun*)^133^, targets of β-cell expression program regulator CREB (*Per1*, *C2cd4b*, *Nr4a2*, *Fos*, *Dusp1*)^133,134^, and genes involved in management of metabolic stress involved in insulin production and secretion in non-diabetic β-cells (*Egr1*, *Hspa1b*, *Ddit3*, and *Dnajb1*)^107,135^. This indicates that β-cell phenotype is compromised across diabetes models. In contrast, some gene groups were conversely expressed in NOD and db/db+mSTZ analyses. For example, NOD group T1-down3, containing some genes involved in adaptive stress response (*Txnip*, *Herpud1*)^44,136^, was, in addition to healthy cells, also highly expressed in db/db and mSTZ model cells.

As it has been previously reported that diabetes results in dedifferentiation of β-cells towards less mature states in both mouse and human^31,32,40,137^ we compared the expression of upregulated genes across postnatal β-cell states and embryonic cell types, including endocrine cells and their progenitors. Among both the NOD and db/db+mSTZ upregulated genes we found genes that were strongly expressed in embryonic data or were specific to diabetes model cells (**Supplementary Figure S13c**). This shows that changes in diabetes-model involve both dedifferentiation as well as diabetes-model specific responses.

In order to validate our findings, we further examined whether DEGs are translatable to other datasets. In the Feng dataset, which is not part of the atlas and contains STZ-treated samples^41^, most T2-groups had the expected expression direction in the STZ model cells (**Supplementary Figure S13b**). However, two gene groups (T2-down1 and T2-down5) did not show different expression activity between diabetic-model and healthy Feng cells. For group T2-down5 the discrepancy could be explained by the gene group being most highly expressed in immature healthy cell states from MIA (**Figure 6a**), which, as discussed above, are absent in the Feng dataset (**Supplementary Figure S9a,b**). In contrast, group T2-down1 had relatively low expression difference between diabetic and healthy MIA cell states (**Figure 6a**). For both gene groups, the observed expression patterns in MIA already indicate that they may not generalize to other datasets that have somewhat different healthy and diseased cell state composition. The dissection of DEGs based on MIA β-cell states enabled us to explain why a subset of DEGs may not be translatable to other datasets, which is a common, usually unexplained, problem in scRNA-seq studies.

To support RNA-level DGE results (**Supplementary Table S10**) at protein-level, we selected relatively highly expressed DEGs (**Figure 6d,e**) and stained them with specific antibodies in islets from healthy and diabetes model (NOD and db/db) mice. First, we validated that islets contain expected healthy and dysfunctional β-cell states by profiling the protein expression of insulin, an established maturation marker Ucn3^17^, and a dedifferentiation marker Aldh1a3^138,139^ (**Figure 6d,e**, **Supplementary Note S10**). We next profiled three novel markers of T2D model: Nucb2, which is involved in insulin secretion^140,141^ and whose mutations were reported to be associated with diabetes risk^142^, Fkbp11, an ER-located chaperone previously reported to be upregulated in certain mouse T2D models^143,144^, and Mt3, which was reported to be associated with β-cell death^145^. Protein and RNA levels of Nucb2 were upregulated in both NOD and db/db islets and Fkbp11 and Mt3 in the db/db islets. This validation supports the observations from our DGE analysis and proposes novel dysfunction markers on both the RNA and protein level.

When comparing NOD and db/db+mSTZ genes to multiple human datasets we did not observe the expected DEG group activity differences between healthy and diabetic samples in a consistent manner (**Supplementary Figure S13b**). However, certain diabetes hallmark genes translate across the species. For example, *DGKB*, which is associated with human T2D^146^, was upregulated in our db/db+mSTZ analysis. Thus, future studies could use our diabetes DGE results to query for molecular changes shared with humans and thus assess if pathways of interest could be further profiled with NOD, db/db, or mSTZ models.

#### Diabetes progression in β-cells reveals a shared intermediate state in type 1 and 2 diabetes models

One of the key goals of diabetes research is to understand the transition from pre-diabetes to diabetes and back upon treatment in order to identify disease states where remission is still possible. To decipher the relationships between healthy and diseased states we calculated a partition-based graph abstraction (PAGA) on the fine β-cell states (**Figure 6f**). The connection from the main healthy state (adult2, containing healthy adult cells across dataset) to the T1D model state (14-16wNOD) or the T2D model state (db/db+mSTZ) led in both cases via an intermediate state (D-inter.). Indeed, it has been suggested previously that both T1D and T2D may share some molecular stress patterns in β-cells, but diverge in the final outcome due to a persistent immune or metabolic challenge, respectively^36,37,147–149^. However, we did not find a report of a shared intermediate state in T1D and T2D models.

The intermediate state contained both stressed healthy and diabetic cells (**Supplementary Figure S7d, Supplementary Note S7**), including cells from the Feng dataset mapped onto MIA **(Figure 4c**). However, the sample with the largest cell proportion localizing in this state was the mSTZ diabetes model sample with regenerative anti-diabetic treatment^40^ (GLP-1+estrogen+insulin; **Supplementary Figure S7d**). This indicates that the intermediate state may be related to either treatment effects or diabetes progression and β-cell stress.

Molecular differences between the healthy and the intermediate state resembled those observed in the diabetic states (14-16wNOD, db/db+mSTZ) (**Supplementary Figure S10c,e**), as described in **Supplementary Note S11**. As the intermediate state may be related to both T1D and T2D models we profiled expression of diabetes DEGs shared between T1D model and T2D model DGE analyses (described above). Most of these genes already exhibited expression differences between the healthy and the intermediate state and further changed from the intermediate to the diabetes model states (**Figure 6g**, **Supplementary Note S11**). Interestingly, shared downregulated genes (89 genes) were strongly enriched for response to extracellular stimuli and transcription factor regulation of gene expression due to genes of activator protein-1 (AP1) complex, which are involved in cell survival and death^150^. This indicates that regulatory mechanisms are disrupted between the healthy and intermediate states.

Our analysis suggests that the intermediate state presents a snapshot of the transition between healthy and dysfunctional cells in different diabetes models. However, it is unclear whether this is part of disease progression or a result of treatment and further investigations are required to clarify this state.

#### Sex differences in β-cells of aged mice involve diabetes-associated genes

Sex differences affect normal β-cell function and subsequent development of diabetes^151–154^. Therefore, we assessed sex differences at across ages and their relationships to diabetes models. Two datasets from early postnatal (P16) and aged (2 years) mice with a mixture of male and female cells were used. In P16 mice we did not observe any DEGs, except for sex-linked Y-chromosome genes (*Ddx3y*, *Eif2s3y*, and *Uty*), which were also used during data preprocessing for sex-annotation of cells. More DEGs were observed in aged mice (26 male and 116 female upregulated genes, **Supplementary Table S11**), which is also reflected in the clear separation of these cells into two distinct states (**Figure 5a**). To further dissect the aged DEGs we clustered them based on expression across all β-cells of MIA, resulting in four female and four male groups (female1-4 and male1-4, **Figure 6c**, **Supplementary Figure S14**, **Supplementary Table S11**).

Females are known to have higher insulin production and are less prone to develop T2D^27,155^. Indeed, we observed some DEG groups explaining these phenotypes. Group male-4, which was highly expressed in T2D model state (**Figure 6c**), contained multiple genes related to dedifferentiation, immaturity, and other endocrine cell types^74,138,156–158^ (**Supplementary Table S11**). In contrast, the female-1 group, which was likewise expressed in T2D model state (**Figure 6c**), contained multiple genes previously reported to be upregulated in pregnancy^32,159^ (**Supplementary Table S11**) as well as genes related to insulin secretion (*Chgb*)^160^ and stress response (*Mapk4* and *Gpx3*)^161,162^. Furthermore, a group expressed specifically in old female cells (female-4, 78 genes, **Figure 6c**), contained some genes involved in insulin regulation^163–165^ and glucose metabolism^166,167^ (**Supplementary Table S11**). Altogether, this indicates that female β-cells are more inclined to diabetes-associated compensation and male β-cells to loss of identity.

## Conclusion and perspectives

Here we present a high-quality mouse islet atlas (MIA) that compiles multiple developmental stages and disease conditions from 56 samples with transcriptomics readouts of over 300,000 cells. The exploration of MIA provides new insights into islet biology and diabetes research that could not have been obtained from individual datasets. Our key discoveries are the description of β-cell landscape from diverse datasets, proposition that mSTZ diabetes model molecularly resembles T2D rather than T1D, and the identification of molecular pathways involved in different types of β-cell dysfunction. While this paper is focused on β-cells, we also showcased that MIA can be used for studying other cell types, presenting an opportunity for future studies.

We used MIA to comprehensively describe β-cell landscape across datasets and conditions. We identified molecular variation conserved across healthy adult β-cells. This included pathways of immaturity and aging as well as pathways potentially involved in cycling between insulin-production and metabolic stress, followed by regeneration. We further proposed the use of gene programs to identify and characterize molecularly distinct cell states in the β-cell landscape. This led to the identification of a new intermediate β-cell state between healthy controls and different diabetes models that may be involved in diabetes progression or treatment-induced remission. We also observed two novel populations within the mSTZ model differing in immaturity and compensatory phenotype, which may be of relevance when using the STZ model in future diabetes studies. Importantly, when comparing different diabetes models, we observed that β-cells in the STZ model exhibited gene expression profile akin to the db/db model and not the NOD model. This was again reflected in comparison to human data, where mSTZ β-cells showed upregulation of T2D-related metabolic stress pathways, while lacking upregulation of T1D-related immune pathways.

For future studies, MIA enables automatic cell type and state transfer as well as cross-study and cross-condition comparison by embedding cells into a shared reference space. We have demonstrated this with the Feng dataset, which is not part of MIA, resulting in the expected mapping of healthy control and STZ diabetes model β-cells to the corresponding MIA regions. This also showed that the immature populations present in MIA and the Feng dataset differ, indicating that the reason for them not sharing markers is likely of biological nature, attributed to different cell states. Our vision is that future studies can similarly map their datasets on top of MIA and publicly provide the generated embeddings in order to further extend the conditions compiled in MIA. As an example, we showed this for a young (P3) sample from the Feng dataset, for which we do not have a matched developmental stage in MIA, with its embedding filling the gap between our embryonic and older postnatal samples.

The heterogeneity compiled within MIA also enables contextualization at the gene-level. For example, known β-cell maturity and dysfunction markers are more heterogeneous than expected, showing distinct expression subgroups across β-cells states of MIA. Similarly, researchers could use the interactive cellxgene instance of MIA to analyze expression of their genes of interest across cell types and diverse biological conditions within MIA.

Our next aim was to describe which pathways are involved in different β-cell dysfunction phenotypes. Therefore, we used MIA to group DEGs and contextualize them based on expression across other conditions. For diabetes-model DEGs this approach revealed phenotype specific as well as shared molecular changes across diabetes models, aging, and immaturity. Grouping of DEGs also identified distinct dysfunction-associated changes across sexes, explaining lower susceptibility of females for diabetes due to upregulation of compensatory rather than loss of identity pathways that were observed in males. In the future, the dissection of dysfunction patterns based on multiple phenotypes may provide valuable insights for personalized medicine, which is based on the knowledge about different disease-associated molecular patterns. It may also be useful for drug repurposing, which relies on pathways shared across diseases^168–170^. For example, it was previously shown that removal of senescent cell populations in NOD mice and models of aging improves overall regulation of glucose levels^36,128^. Indeed, in our analysis we observed upregulation of senescence-associated genes in both aged and T1D model cells.

We show that our results are reproducible in independent mouse transcriptomic data and in newly generated antibody-stainings, proposing new markers of T2D-model associated dysfunction (Nucb2, Fkbp11, Mt3). Comparison to human datasets revealed some similarities to mice, however, novel methods will be required to improve cross-species comparison and translation.

In conclusion, MIA provides a useful tool for making new discoveries in islet biology and diabetes research. It is available as a curated resource in formats that enable interactive exploration via cellxgene and computational analyses (https://github.com/theislab/mouse_cross-condition_pancreatic_islet_atlas), including access to the cellxgene curated dataset via Sfaira^171^). Our novel discoveries in β-cell biology showcase how MIA can be used both as a reference of cell states as well as for further querying of gene expression across conditions.

## Methods

### Data

#### Generation of new mouse samples included in the atlas

Animal breeding and sampling of material were carried out in compliance with the applicable legislation and the guidelines of the Society of Laboratory Animals (GV-SOLAS) and of the Federation of Laboratory Animal Science Associations (FELASA). Mice were housed in groups of two to four animals and maintained at 23 ± 1 ⁰C and 45-65 % humidity on a 12-hour dark/light cycle with ad libitum access to diet and water.

Islets of Langerhans have been isolated using a standard protocol^172,173^. The aged dataset was generated from islets of Langerhans isolated from the Fltp lineage tracing mouse model (Fltp iCre mTmG)^174^ in mice older than 2 years. Two male and two female mice were pooled together after islet isolation and prior to fluorescence-activated cell sorting (FACS) into Fltp negative (tomato positive), Fltp lineage positive (GFP positive), and Fltp transient (double positive) populations (**Supplementary Figure S15**). Separate libraries were generated for each sorted population after pooling across sexes. For the 4m dataset, we used the Fltp reporter mouse line Fltp^ZV175^. Pancreas head and tail were anatomically separated prior to islet isolation. Islets from six Fltp^ZV/+^ male mice were pooled. Subsequently, Fltp Venus reporter positive and negative cells were sorted (**Supplementary Figure S15**), thus generating four libraries. Metadata of all samples is collected in the **Supplementary Table S1**.

Libraries of single cells were produced using the Chromium Single-Cell 3′ library and 10XGenomics gel bead kit v3.1 (PN #1000121) in the aged dataset and with v2 (PN #120237) in the 4m dataset. Briefly, 10,000 cells were loaded per channel of a 10x chip to produce gel bead-in-emulsions (GEMs). Then the samples underwent reverse transcription to barcoded RNA followed by cleanup, cDNA amplification, enzymatic fragmentation, 5′ adaptor, and sample index attachment. The samples of the aged dataset were sequenced using a NovaSeq6000 (Illumina) with 100 bp paired-end sequencing and the samples of 4m dataset were sequenced using a HiSeq4000 (Illumina) with 150 bp paired-end sequencing of read 2.

#### Datasets included in the atlas

We used nine mouse pancreatic islet scRNA-seq datasets previously generated with 10X Genomics Chromium technology. Data availability is described in **Table 1**. Public data was obtained from GEO in July 2020 by comprehensively searching for mouse pancreatic islet scRNA seq datasets. From the collected datasets we excluded datasets that would not be applicable for analysis of β-cell heterogeneity, such as cancer and reprogramming datasets as well as datasets with low endocrine cell counts, including embryonic datasets, with the exception of an in-house embryonic dataset. We also excluded datasets that were not generated with Chromium (namely Smart-seq2) as most of them had low cell counts and could lead to strong cross-technology batch effects due to differences in sensitivity and bias in the type of captured genes^176^. Furthermore, some of the integration methods are not designed for full-length reads, such as Smart-seq2^55^. Altogether, using additional sequencing technologies would make the integration more challenging.

All computational analyses of scRNA-seq data were performed with Scanpy (v1.6 - 1.8.1)^177^, except where noted elsewhere.

#### Datasets for atlas validation

For validation we collected public mouse and human scRNA-seq datasets (**Table 2**, **Supplementary Table S12**) and downloaded their expression count matrices and metadata from GEO and paper supplements. If raw counts were available, re-normalization was performed with the Scanpy normalize_total function, otherwise, the available pre-normalized data was used. For downstream analyses *log*(*expr* + 1) transformed normalized expression was used. We manually unified cell type annotation from original studies to a shared set of cell type names by renaming existing labels. No further preprocessing was performed on these datasets. These datasets were not included in the atlas and were always analyzed individually. In the text we refer to the GSE137909 dataset as the Feng dataset.

Where necessary, we mapped genes across species based on orthologue information from BioMart^182^ with Ensembl Genes v103.

### Preprocessing of datasets integrated in the atlas

Gene expression counts were calculated based on genome versions described in **Table 1** with 10x Genomics Cell Ranger (v2.2.1-v3.1.0)^183^. Each dataset was separately preprocessed with the below described steps, except when we note that a processing step was performed per-sample, and filtering thresholds were determined on a per-dataset level.

#### Ambient gene selection

To reduce the effect of ambient expression on embedding calculation we removed most prominent ambient genes, which were identified as described here. We selected likely empty droplets that contained only ambient RNA based on having less than 100 counts. Gene proportions within empty droplets were computed on raw counts per sample, representing gene proportions within the ambient RNA. Genes with highest ambient proportion were selected with a dataset-specific ambient proportion threshold, selecting genes as the union across samples, generating a set of approximately 20 genes per dataset. Due to the proportional nature of expression measurements a relatively high ambient proportion of some genes leads to lower proportions in other ambient genes. Thus, we reduced the ambient threshold when some genes had a relatively high ambient proportion in order to also capture less ambient genes that are nevertheless known to strongly affect ambient profiles, such as endocrine hormone genes. Additionally, a larger set of approximately 100 genes was generated with a more permissive threshold that aimed to include top ambient genes so that selecting more genes would no longer evidently increase the captured cumulative ambient proportion given by the sum of the per-gene ambient proportions.

#### Quality control

Empty droplet score was computed per sample with DropletUtils (v1.10.3)^184^ *emptyDrops* function using *LogProb* output for downstream visual QC assessment purposes. Cell-containing droplets as determined by the Cell Ranger pipeline were used in downstream analyses. Cell filtering was performed based on guidelines from^185^, excluding cells with a low number of expressed genes, low total counts or high mitochondrial proportion and outliers with a very high number of total counts or expressed genes. Genes expressed in a very small number of cells and top ambient genes were excluded for the purpose of annotation and integration. Doublets were filtered out with Scrublet (v0.2.1)^186^ scores computed per-sample using a manually set threshold to separate the scores into cross-cell type doublet and potential non-doublet populations as proposed in the tutorial^186^, while ensuring that selected doublet cells mainly mapped into discrete cluster locations on the UMAP embedding. The choice of the threshold was set permissively, as indicated by the presence of some residual doublet populations in the final atlas version.

#### Annotation

To perform cell annotation within individual datasets normalization was performed per dataset with scran (v1.16.0-1.18.7) pooled size factors^187,188^, data was *log*(*expr* + 1) transformed, and 2000 highly variable genes (HVG) were selected with Scanpy using the *cell_ranger* selection flavor and samples as batches. The cell cycle stage of each cell was annotated using the Cyclone method^189^ as implemented in scran. For datasets without per-cell sex information the sex was annotated based on Y-chromosome located HVGs with high expression. We assigned cells into insulin, glucagon, somatostatin, and pancreatic polypeptide high or low groups per-sample based on scores from the Scanpy *score_genes* function. Cell types were annotated in the following datasets: P16, 4m, aged, mSTZ (healthy sample), db/db (healthy samples), based on known pancreatic cell type markers followed by recursive sub-clustering until homogenous clusters were reached. Rare cell types that did not form a separate cluster were annotated based on per-cell marker scores (e.g., ε-cells in the P16 dataset). Here and in the below re-annotation of the integrated data we relied on the following cell type markers across multiple datasets, although on the per-dataset level we also used other markers, expressed in cell subpopulations present in only some of the datasets. Marker list: acinar: *Cpa1*, *Prss2*; α: *Gcg*; β: *Ins1*, *Ins2*; δ: *Sst*; ductal: *Krt19*, *Muc1*, *Sox9*; endothelial: *Pecam1*, *Plvap*; ε: *Ghrl*; γ: *Ppy*; immune: *Cd52*, *Lyz2*, *Ptprc*; stellate activated: *Col1a2*, *Bicc1*, *Pdgfra*; stellate quiescent: *Ndufa4l2*, *Acta2*, *Cspg4*, *Rgs5*; Schwann: *Cryab*, *Plp1*, *Sox10*. Expected multiplet rates were computed and together with Scrublet scores used to determine which annotated multiplet cell types present true cells or residual multiplets. We annotated β-cell states based on expression of known β-cell heterogeneity markers.

### Integration

#### Preprocessing and ambient correction for integration

We tested different methods for ambient expression correction: CellBender (v0.2.0)^190^, SoupX (v1.5.0)^191^, and DecontX (from celda v1.5)^192^. We did not use CellBender preprocessed data further as we observed non-homogeneous correction within clusters, namely some genes known to be cell-type specific, such as β-cell specific *Ins1* and *Ins2*, were removed partially and at different levels across cells within other cell types. For other methods different ambient correction strengths were used and one or more were selected for integration per method. Non-ambient-corrected data was also used. Top ambient genes were excluded, also in ambient corrected datasets (using the smaller ambient gene set). The ambient correction method selected for final integration is described in the integration selection section. Genes previously marked as too lowly expressed on a per-dataset level were also removed. To enable integration with samples as batches and future mapping of new samples onto the reference the data was per-sample scran normalized and transformed with *log*(*expr* + 1). We re-normalized data on the sample-level as we observed that when performing separate scran normalizations across different data subsets (e.g., samples or datasets) biologically related cells have different total number of counts across the data subsets (not shown). By performing batch-wise normalization (here batch is sample) we ensure that the integration model can account for this effect when removing batch effects. For scVI integration non-normalized data was used. Expression matrices of all samples were merged, retaining intersection of genes. The 2000 HVGs obtained with the scIB (developmental version, last update on 17. 1. 2022)^55^ *hvg_batch* function was used.

#### Integration selection

For integration we used scVI v0.7.0a5^54^ with hyperopt hyperparameter optimization and scArches v0.1.5^56^ with manual parameter optimization. First, we performed integration on the annotated data only to select scVI parameters with hyperopt (number of network layers and their size, number of latent dimensions, reconstruction loss, dropout rate, learning rate, gene dispersion, and number of epochs) and scArches parameters based on visual evaluation (different HVG selection, integration strength regulated by the weight between reconstruction and Kullback–Leibler divergence loss, number of network layers, and reconstruction metrics), to ensure that selected parameters lead to a reasonable integration. Afterwards, integration was performed on all data. Different integration methods and preprocessing combinations were evaluated with scIB metrics. We added a new biological conservation metric named Moran’s I conservation, which does not require cell type annotation. For biological conservation evaluation we excluded unannotated and multiplet cells, except for Moran’s I, which could be run on all cells. As annotation was available only for a subset of cells the batch correction metrics were run both on all data, using clusters instead of cell type labels, and on the annotated data subset. We also performed evaluation on β-cells only, using β-cell states as cell labels, with different integration strengths. Top selected integrations were run multiple times to better distinguish between random-initialization and true performance variation. The best method (removed top ambient genes and scArches-cVAE) was selected based on summarized biological conservation and batch correction scores, as described in scIB, with special focus on β-cell state conservation.

We also tested β-cell specific integration, using β-cells defined based on an integrated annotation (see below) with the same integration settings as for the whole atlas, but with multiple different integration strengths in scArches-cVAE. Batch correction evaluation was run on all cells, using clusters instead of cell type labels, and biological preservation evaluation on cells that had state annotation. The results were compared to metrics computed on the same set of cells from the whole atlas integration.

For comparison we also show unintegrated embedding, that was computed using the same set of genes as the final atlas integration. We normalized expression using the Scanpy *normalize_total* function as Scran normalization performed on individual samples, as used for integration, leads to less comparable normalization factors across samples. Data was *log*(*expr* + 1) transformed and scaled, followed by PCA embedding computation that was used as the basis for UMAP.

#### Integration evaluation with Moran’s I conservation

We proposed a new biological conservation metric for comparison across integration runs without the need for cell type annotation that determines how strongly genes are variable across the integrated embedding. Namely, if embedding captures biological variation at a finer scale, e.g., within cell types, then the expression variation of genes that are potential determinants of cell state differences (e.g., HVGs) should be non-random across the embedding. The method first computes HVGs (*g*, 1000 genes) on the expression data with Scanpy *highly_variable_genes* function using *cell_ranger flavor* and *batch_key* parameters. Moran’s I for these HVGs is then computed on the integrated embedding (*i*) with Scanpy *morans_i* function. This function uses information about each cell’s k-nearest neighbors graph computed with Scanpy *neighbors* function on the integrated embedding with Euclidean distance metric. The final score is computed as the mean of per-gene scores. This score is rescaled to fall within range [0,1], matching other scIB scores. This can be formulated as:

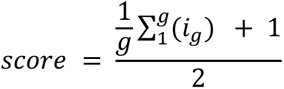

### Analysis of the integrated atlas

#### Clustering and annotation

We defined new cell types on the integrated atlas by consecutive Leiden^193^ subclustering with Scanpy, namely by manually selecting clusters to be subclustered as needed to separate cell types, relying on information about previously annotated cells, hormone expression high/low assignment, and quality metrics. Namely, empty droplets were identified based on low expression and high empty droplet probability and doublet clusters based on higher doublet scores and expression of markers of multiple cell types. We compared the re-annotation to the annotation from original publications, for which we manually unified cell type labels by renaming the labels to a shared set of names.

As scran normalization performed per-sample is not comparable across samples (described above) scran size factors were recalculated on the integrated cell clusters and the atlas was jointly re-normalized. In downstream analyses we used this normalized data, except for the methods that require raw counts.

To disentangle biologically relevant differentially active genes from genes whose expression is likely a result of ambient expression differences in the downstream analyses we defined genes that may be predominately ambiently expressed in a given cell type. Top ambient genes likely not coming from β-cells were defined as follows. For each sample, genes with high expression in empty droplets, containing less than 100 counts, were selected with a single threshold across all samples, and the genes were pooled across samples. These ambient genes were clustered based on expression across integrated cell clusters. Ambient gene clusters were assigned to non-β-cell originating ambient genes if they had relatively low expression across all β-cell clusters compared to cell clusters coming from other cell types. Besides making the set of likely non -β-cell ambient genes, we used during interpretation a per-gene metric that can indicate ambient gene origin, namely relative gene expression in a cell type compared to other cell types, with higher scoring genes being less likely ambient. As this metric was used for postnatal endocrine analyzes the embryonic clusters were excluded as they are not expected to contribute to ambience in postnatal samples. The atlas subset was then sub-clustered using Leiden clustering with resolution=2. Mean expression in cell clusters was maxabs-scaled across clusters, representing relative expression in each cluster. To determine relative expression of a gene in a cell type we used the highest relative expression obtained across all cell clusters containing predominantly that cell type.

In all further analyses where we needed to reduce the number of cells due to computational constraints we prepared pseudobulk data (here termed “fine pseudobulk”) by Leiden clustering with high resolution (such as resolution=20) in order to create tens or hundreds of clusters (depending on data size) that should capture the majority of heterogeneity within the data. This is akin to recently proposed methods that aim at creating so called “metacells” that group together cells without biological differences^194,195^. Pseudobulk expression was computed as the mean of *log*(*expr* + 1) transformed normalized expression within each cluster. For DGE analysis on pseudobulk (here termed “metadata-based pseudobulk”) we grouped cells based on their metadata, such as sample and cell type, as before suggested for single cell DGE analysis^196^. Here normalized counts were summed across cells and *log*(*expr* + 1) transformation was not applied.

#### Identification of endocrine cell type markers

For the identification of endocrine cell type markers one-vs-one DGE analyzes were performed with edgeR (v3.32.1)^197^. For the postnatal markers metadata-based pseudobulks of postnatal datasets per cell type, sample, and sex were created. We excluded embryonic, doublet, and endocrine proliferative cell types. The former cell type was excluded as a minute number of postnatal cells mapped to the embryonic clusters (**Supplementary Figure S2**). The latter two cell type groups were excluded as they share gene expression with matched non-doublet and non-proliferative cell types, which would prevent identification of these genes as DGE markers. Lowly expressed genes were removed with edgeR and a single DGE test was fitted, using GLM with robust dispersion, with sample and sex as covariates, and two-sided likelihood-ratio significance testing. To obtain one-vs-rest upregulated genes for each endocrine cell type the factors across cell types were compared. Marker genes were selected based on false discovery rate (FDR) below 0.05 and log fold change (lFC) above 1.5 against all other cell types. In the supplementary tables we reported maximal adjusted p-values across compared cell types and for lFC we reported 0 if lFCs across comparisons had both negative and positive values and otherwise signed minimal lFC based on absolute value sorting. For embryonic markers the embryonic dataset with cell type annotation from the original study^85^ was used. The Fev+ cluster was excluded as it contained precursors of individual endocrine cell types with similar expressions as in the descendant cell types, which would prevent identification of markers. Metadata-based pseudobulks were created per cell type and sample while sex was not used as a covariate, as at this age strong sex differences were not expected. Endocrine cell type markers were identified as for the postnatal datasets. In the postnatal dataset we used 52 samples and in the embryonic dataset 4 samples, with some cell types being represented in fewer samples and some samples containing data pooled across multiple animals.

#### Comparison of embryonic and postnatal endocrine cells

We grouped α-, β-, and δ-cells into three groups per cell type: embryo - cells that were annotated as a certain endocrine cell type in the original embryo study and mapped into the embryo endocrine atlas cluster, embryo postnatal-like - cells from the embryo dataset that mapped into one of the postnatal endocrine atlas clusters, and postnatal - cells from postnatal datasets that mapped into one of the postnatal endocrine atlas clusters. For embryo and embryo postnatal-like cell types we computed what proportion of embryonic cells per sample-specific age group they represent.

#### Reference mapping of the external mouse dataset

The Feng dataset (query) was re-normalized per-sample with scran and *log*(*expr* + 1) transformation to match atlas (reference) datasets preprocessing. The reference scArches model was used to compute the query embedding, using samples as batches. For query β-cell mapping analysis the cell type annotations from the original study^41^ were used. A joint UMAP embedding of query and reference β-cells was computed, as well as a UMAP with added reference embryonic β-cells, using β-cells from the original study annotation^85^ that mapped into the atlas embryo endocrine cluster, and reference proliferative β-cells, defined as endocrine proliferative cells that were previously annotated as highly expressing insulin, but not other hormones. Query β-cell states were predicted based on atlas coarse β-cell states with the addition of embryonic and proliferative β-cell groups. For cell type transfer a weighted kNN classifier adapted from scArches manuscript^56^ was used with an uncertainty threshold of 0.75.

#### Comparison of diabetes models to human T1D and T2D

To obtain T1D and T2D gene sets conserved across human datasets the T1D or T2D cells were compared against cells from non-diabetic samples in each human dataset (the number of samples in each group varied across datasets, see **Supplementary Table S12** for sample group sizes). Only genes expressed in at least 10% of diabetic or healthy cells per dataset were used. Genes with a FDR below 0.25 and lFC above 0.5 in at least half of the datasets based on the Scanpy *rank_genes_groups* t-test function (two-sided Welch’s test on cell level) were selected.

Gene set enrichment was computed with hypeR (v1.6.0)^198^ at the FDR threshold of 0.25 using GO, KEGG, and Reactome gene sets from MSigDB (v7.4.1). Before enrichment, each gene set was subsetted to genes present in the background that consisted of all genes used for the analysis (here genes tested for DGE) and gene sets containing less than five or more than 500 genes were removed. From enriched gene sets with shared genes, we manually selected representative gene sets to be highlighted in the text.

Mouse diabetes model β-cells were scored for both the newly defined and literature-based gene sets with Scanpy *score_genes* function on each dataset. Comparisons were performed between the following groups: in the 8-16wNOD dataset the 8-week (healthy) vs 14- and 16-week samples (diabetic); in the mSTZ dataset control (healthy) vs the mSTZ treated sample (diabetic), and in the db/db dataset control (health) vs db/db sham-operated samples (diabetic). Gene set score distributions in healthy and diabetic groups within each dataset (sample numbers for healthy: mSTZ =1, db/db = 2, 8-16wNOD = 3, and diabetic: mSTZ =1, db/db = 2, 8-16wNOD = 6, with some samples containing pooled animals) were compared with a two-sided Mann-Whitney U test on cell level and a natural-logarithm based lFC was computed between distribution medians.

#### Coarse β-cell states and their markers

Clusters were computed with the Scanpy Leiden function and were thereafter added descriptive annotation based on sample ratios across clusters, relying on sample metadata, quality scores, and relationships between clusters determined with PAGA. Initial clustering was performed with a relatively high resolution so that we could later merge clusters that we could not interpret as separate based on the criteria described above while ensuring that we did not miss any unique clusters.

Cluster-specific markers conserved across datasets were computed as follows. Data was subsetted to exclude low quality clusters and the embryo dataset as it contained too few β-cells (less than 20 per sample across all β-cell clusters). Cell groups used for DGE were defined as a combination of cluster and dataset, using for each cluster only datasets with a high proportion of cells in that cluster in at least one sample. For each dataset-cluster group DGE analysis was performed with the Scanpy *rank_genes_groups* t-test function against all other cell groups, except the ones from the same cluster, excluding genes that were lowly expressed in both clusters before DGE analysis. The number of samples per group varied across cell states, with the total number of considered samples before grouping being 52, with some samples containing pooled animals. As markers we selected genes that were significantly upregulated (FDR below 0.1 and lFC above 0) in all datasets across all other cell groups and for plotting genes were prioritized based on the highest minimal lFC across all comparisons. Genes were further filtered to select likely non-ambient genes by keeping only genes with relatively high expression in β-cells (above 0.7). Hemoglobin genes were also removed as they were not caught by the relative expression filter since erythrocytes are absent from data, but the transcripts are still present in the ambient RNA.

Markers of adult, immature, and T2D model states were visually validated on the external mouse dataset. The healthy β-cells were grouped by age and the STZ treated cells based on the administration of insulin.

Translation of markers to the human data was tested based on all collected human datasets with per-dataset one vs rest one-sided t-tests on cell level and p-value based significance threshold of 0.05. We also report log2-based lFC between group means. The following cell groups were defined: T1D or T2D groups contained all cells annotated as T1D or T2D and were used to test both known T1D or T2D markers as well as our NOD or db/db+mSTZ markers, respectively, and for other marker groups only healthy donor cells were used, with the adult set used to test our adult mouse cluster and contained ages of 19-64 years, mature set used to test known maturity markers and contained ages of 19 years or more, aged male or female sets contained ages of 65 years or more and immature set ages of 18 years or less. Age groups were defined based on OLS HsapDv human life cycle stages definitions ^199^. The number of samples varied across groups and datasets (see **Supplementary Table S12** for more details).

#### Gene programs in β-cells

To define GPs we first identified genes variable across embedding and then clustered them based on coexpression (**Figure 5d**), as described below. To identify variable genes low quality coarse β-cell clusters were excluded prior to the analysis as they could lead to high spatial autocorrelation scores of genes associated with data quality. Lowly expressed genes and the non-β-cell ambient gene set were removed. Moran’s I was used to assess the autocorrelation of expression across the integrated embedding (all 15 dimensions). We observed a bias of genes expressed in fewer cells towards lower Moran’s I, which would lead to lowly expressed genes unjustly being less often selected as variable based on Moran’s I threshold. To account for this bias, we regressed out the effect of the number of cells expressing the gene on Moran’s I. For this regression we used genes likely not to be truly variable across the embedding, as explained below, to estimate the base-level effect of expression sparsity across cells on Moran’s I. Genes likely not to be truly variable were selected as follows: Most highly expressed genes (N cells >= 40,000 from total of 99,361 cells) were excluded as they were deviating from the trend towards higher Moran’s I-s, which was likely due to their importance in β-cell function and thus higher variability across the β-cell embedding. The remaining genes were binned (N bins = 20) based on the number of cells in which they were expressed and the five genes with lowest Moran’s I from each bin were selected for regression, representing the base-level (likely not biologically relevant variable) Moran’s I at certain expression strength. The regression was fitted on the selected genes and then the corrected Moran’s I score was computed as the residuals from regression for all genes for which the uncorrected Moran’s I score was initially computed. Finally, GPs were defined by selecting genes with the highest corrected Moran’s I and clustering them using fine pseudobulk cell clusters as features with hierarchical clustering and visually determined cutting threshold based on a heatmap of gene expression across pseudobulks. Gene set enrichment of GPs was computed as for the human T1D and T2D conserved genes. We supplemented GP gene set enrichment interpretation with marker-based domain knowledge to support β-cell specific functional annotation, which is not fully encompassed by the more generic gene sets available in KEGG, GO, and Reactome.

The ratio of variance explained by GPs per dataset was computed based on principal component (PC) regression. For each dataset, lowly expressed genes were removed and 50 PCs were computed based on HVGs. Cells were scored for GP activities with the Scanpy *score_genes* function (excluding genes missing from each dataset from GPs) to analyze how well GP scores of all or individual GPs explain each PC component based on regression R^2^ (coefficient of determination) was computed. Total variance explained was computed as a sum of R^2^ across PCs weighted by ratio of variance explained by each corresponding PC. For comparison, the same procedure was used to evaluate variance explained by random gene groups of the same size as the GPs, repeating the procedure 10 times to estimate the random distribution. For the analysis of explained variance in healthy mouse and human samples, only samples with at least 100 β-cells were used and the explained variance was computed as described above, repeating calculation for random gene groups 100 times. The significance of the explained variance by GPs was computed as an one-sided empirical p-value compared to the distribution for the matched random gene group.

#### Fine β-cell states

Each cell was scored for each GP with the Scanpy *score_genes* function followed by averaging within the fine pseudobulk clusters to speed up further analysis. The GP scores were used as features to cluster pseudobulk clusters into β-cell state clusters using hierarchical clustering followed by visual selection of the cutting threshold based on GP activity purity within clusters and unique pattern of GPs across clusters. Each cell was assigned to the cluster of its pseudobulk group. The clusters were named based on metadata of the samples with a large proportion of cells within the cluster. The resulting β-cell state clusters were used to obtain a pruned PAGA graph, selecting a pruning threshold that separated between high and low connectivities.

We analyzed GP-based molecular differences for individual datasets between healthy and diseased states (adult2 vs db/db+mSTZ (for datasets db/db and mSTZ) and vs NOD-D (for dataset 8-16wNOD)) and two diseased states (db/db+mSTZ and mSTZ for dataset mSTZ). All β-cells were scored for GP activity with the Scanpy *score_genes* function and individual scores were normalized across cells to [0,1] with winsorizing by removing the highest and lowest 20 cells for setting the scaling range. The per-dataset differences between means of the normalized scores within clusters were then used for cluster comparison

We manually extracted known markers of β-cell heterogeneity from literature. For plotting across fine β-cell states we excluded markers expressed in less than 1% of β-cells and plotted mean expression per cell state. Heatmap was created with ComplexHeatmap (v2.11.1)^200,201^.

#### Conserved β-cell heterogeneity in healthy samples

Low quality coarse β-cell clusters were excluded as they could lead to high spatial autocorrelation scores of genes associated with data quality. Control samples from the chem dataset were not used as they showed lower integration of β-cells, indicating potential strong batch effects, which could negatively affect identification of variable gene groups conserved in healthy β-cells. Thus, healthy adult samples from db/db, STZ, and young_adult datasets were used. For each sample, lowly expressed genes were removed and a neighborhood graph was computed on per-sample PC embedding for Moran’s I computation, as described in the GPs section. Here, we adjusted the threshold for removing genes expressed in many cells from the Moran’s I score correction regression to expression in at least 30% of cells. Genes with high Moran’s I in all samples were selected. To ensure that gene clusters are conserved across samples the genes were clustered based on the highest distance on per-sample fine pseudobulks using hierarchical clustering. The cutting threshold was visually determined based on a heatmap of gene expression across per-sample pseudobulk. Gene group scores were compared to expression of known β-cell functional and phenotypic markers extracted from literature, with marker correlations computed on per-sample pseudobulks and summarized as a mean of per-dataset means across per-sample scores. Gene set enrichment was computed as for β-cell GPs.

To find the cells with the highest expression of each gene group we used Scanpy *score_genes* function on individual healthy adult samples, followed by selection of 50 cells with the highest score. As the Feng dataset had a low number of healthy adult β-cells we performed scoring on all control samples together and selected only top 20 cells per gene group.

#### Differential expression in T1D model and T2D model β-cells

We performed DGE analysis on all samples from 8-16wNOD (n=9) and from db/db and mSTZ (n=15, some samples contained pooled animals) datasets, excluding low quality coarse β-cell clusters. A continuous disease process (**Supplementary Figure S13a**) was computed with MELD (v1.0.0)^202^ on the integrated embedding as healthy sample densities normalized over healthy and diseased densities, using for healthy and diseased the same set of samples as in the diabetes model comparison to human diabetes-associated gene sets. In the db/db+mSTZ analysis the final MELD healthy and diseased scores were computed as a mean over datasets-specific scores. We observe that the resulting process corresponds to the gradient from the healthiest (highest healthy sample cell density within region) to the most diabetically stressed cells (highest diabetes model sample cell density within region), with the process value of individual cells being determined based on cell embedding location rather than just sample membership. Genes expressed in less than 5% of healthy or diabetic sample cells were removed. To assess linear change in gene expression along the disease process we used diffxpy (v0.7.4)^203^ two-sided Wald test that fits a negative binomial model to raw counts across cells using expression normalization size factors as exposure. Dataset information was used as a covariate in the db/db+mSTZ analysis. The DEGs were selected based on FDR below 0.05, lFC (binary logarithm of the relevant model coefficient representing linear change) above 1, and relative expression in β-cells above 0.2, to keep only genes that are less likely ambient, as described above. For comparison to the embryonic data the [0,1]-normalized expression of upregulated genes was plotted across fine β-cell states and embryonic clusters as annotated in the original study.

For both DGE analyses the up and down regulated genes were separately hierarchically clustered on the whole β-cell fine pseudbulk data. Cutting thresholds were selected visually based on heatmaps portraying gene expression grouped across fine pseudobulks. All β-cells were scored for DEG groups with the Scanpy *score_genes* function and the scores were averaged within β-cell clusters. Gene set enrichment was computed as described for human T1D and T2D genes. Gene membership across groups was compared as the relative overlap normalized by the size of the smaller group.

The DEGs in NOD and db/db+mSTZ were compared to three human datasets with T1D samples and one mouse and seven human datasets with T2D samples, respectively. We scored cells for each DEG group activity with the Scanpy *score_genes* function, followed by [0,1] normalization across cells, and separately plotted cells from healthy and diabetic samples.

For analysis of the DGE patterns in relationship to the D-inter. cluster the genes up or downregulated in both NOD and db/db+mSTZ were obtained. We plotted their expression per diabetes model datasets across the adult2, D-inter., and 14-16wNOD (for 8-16wNOD dataset) or db/db+mSTZ (for db/db and mSTZ datasets) clusters. We normalized gene expression across clusters in each dataset to [0,1]. We computed the gene set enrichment of the shared DEGs as for human T1D and T2D genes. The GP differences between adult2 and D-inter. clusters were computed for individual datasets (db/db, mSTZ, 8-16wNOD) as described in the section on the fine β-cell clusters.

#### Differential expression in T1D model and T2D model endocrine cells

To compare DEGs across diabetes models and endocrine cell types we fitted a joint model with edgeR. Cells from healthy adult (datasets: 4m, 8-16wNOD samples aged 8 weeks, db/db control, mSTZ control, n=10, some samples contained pooled animals), T1D model (dataset: NOD_progression samples aged 14 and 16 weeks, n=6), and T2D model (datasets: mSTZ and db/db, both without treatment, n=3) were used to compute metadata-based pseudobulks per disease status group, sample, dataset, sex, and endocrine cell type. Lowly expressed genes were removed with edgeR. A single expression model was fitted, using GLM with robust dispersion, with dataset and sex as covariates. A two-sided likelihood-ratio test was used to compare model factors for each T1D model or T2D model cell type to the corresponding healthy cell type to obtain the T1D model or T2D model effect per cell type. The DEGs were selected based on FDR below 0.05, absolute lFC above 1, and relative expression in individual cell types above 0.1 to focus on genes that are less likely to be ambiently expressed. Overlap between DEGs was computed accounting for DGE direction between the two groups. Same direction DEGs across α-, δ-, and γ-cells in both diabetes types were extracted and gene set enrichment was computed as for human T1D and T2D genes.

#### Sex differences in β-cells during aging

Two datasets that contained a mixture of male and female cells were used: P16 and aged. Each dataset was analyzed separately; both datasets had 3 samples per group with pooled animals within samples. Cells from low quality coarse β-cell clusters, genes expressed in less than 5% of cells, and non-β-cell ambient genes were removed. DGE analysis was performed with sex and samples as covariates using diffxpy two-sided Wald test. We removed genes that could not be fitted, as indicated by extremely small standard deviations of the regression coefficient (sd=2.2e-162). DEGs were selected based on FDR below 0.05 and absolute lFC above 1.

DEGs between sexes in the aged dataset were separated by DGE direction and hierarchically clustered on the whole β-cell fine pseudbulk data. Cutting thresholds were selected visually based on heatmap portraying gene expression across fine pseudobulks. All β-cells were scored for DEG groups with the Scanpy *score_genes* function.

### Laboratory validation of diabetes markers

For diabetes markers validation we used healthy adult mice from strains C57BL/6J (three males and three females, aged 2-4 months) and B6.BKS(D)-Leprdb/J (healthy db/db control), db/db T2D model mice (three males aged 8 weeks), and NOD T1D model mice (three females aged 8 weeks). For endocrine markers validation we used postnatal healthy mice from strain C57BL/6J (two males and one female, at P9 stage). Mice pancreases were dissected and fixed (4% PFA-PBS, 24 hrs at 4 °C). The organs were cryoprotected in a sequential gradient of 7.5, 15, and 30% sucrose-PBS solutions (each solution 2h at room temperature (RT)). Next, pancreases were incubated in 30% sucrose and tissue-freezing medium (Leica) (1:1, overnight at t 4 °C). Afterwards, they were embedded using a tissue-freezing medium. Sections of 20 μm thickness were cut from each sample mounted on a glass slide (Thermo Fisher Scientific).

Islet isolation was performed by collagenase P (Roche) digestion of the adult pancreas. We injected 3 mL of collagenase P (1 mg/mL) into the bile duct, the perfused pancreas was consequently dissected and placed into 3 mL of collagenase P for 15 min at 37 °C. 10 mL of G-solution (HBSS (Lonza) + 1% BSA (Sigma)) was added to the samples followed by centrifugation at 1600 rpm at 4 °C. After another washing step with G-solution, the pellets were resuspended in 5.5 mL of gradient preparation (5 mL 10% RPMI (Lonza) and 3 mL 40% Optiprep (Sigma) per sample), and placed on top of 2.5 mL of the same solution. To form a 3-layers gradient, 6 mL of G-solution was added on the top. Samples were then incubated for 10 min at RT before subjecting to centrifugation at 1700 rpm. Finally, the interphase between the upper and the middle layers of the gradient was harvested and filtered through a 70 μm Nylon filter and washed with G-solution. Islets were handpicked under the microscope, for fixation islets were incubated in 4% PFA-PBS for 15 min at RT.

For immunostaining the cryosections were rehydrated and then permeabilized (0.2% Triton X-100-H2O for 30 min RT). Then, the samples were blocked in a blocking solution (PBS, 0.1% Tween-20, 1% donkey serum, 5% FCS 1h RT). Primary antibodies (**Supplementary Table S13**) were incubated for at least 4 h at RT followed by 3 washes with PBX. The samples were then incubated with secondary antibodies (**Supplementary Table S13**) during 4-5 hrs of incubation. For the anti-Rbp4 antibody we performed antigen retrieval with a citric buffer (10 mM Sodium citrate, 0.05% Tween 20, pH = 6) in addition to the above described protocol. Finally, the pancreatic sections were stained with DAPI (1:500 in 1X PBS for 30 min). All images were obtained with a Leica microscope of the type DMI 6000 using the LAS AF software. Images were analyzed using the LAS AF and/or ImageJ software program.

## Data availability

Up-to-date data resource links are available from https://github.com/theislab/mouse_cross-condition_pancreatic_islet_atlas. The two newly generated scRNA-seq datasets, the integrated atlas, and the reference mapped embedding of the Feng dataset were deposited to GEO within super-series GSE211799. The atlas is also available as a cellxgene instance (https://cellxgene.cziscience.com/collections/296237e2-393d-4e31-b590-b03f74ac5070). The scArches model for reference mapping and an example code for reference mapping used for the Feng dataset are available in https://github.com/theislab/mouse_cross-condition_pancreatic_islet_atlas/tree/main/reference_mapping.

All figures are available in high resolution in: https://drive.google.com/file/d/1paBmvHNP_xzkOVTo104eiAj2Fzpuyknr/view?usp=sharing

All supplementary tables are available in: https://drive.google.com/drive/folders/1eJ6fIaA5hvlvOhsXHSX5L8A_P6dK7PXB?usp=sharing

## Code availability

All code is available at https://github.com/theislab/mouse_cross-condition_pancreatic_islet_atlas. This includes both reproducibility code and an example of how new datasets can be mapped onto the atlas.

## Supporting information

High-resolution figures

## Acknowledgements

We thank Thomas Walzthöni and Xavier Pastor Hostench for bioinformatics support in processing of raw scRNA-seq data provided at the Core Facility Genomics, Helmholtz Zentrum Munich, Munich, Germany. We are grateful to the members of Theis and Lickert lab and scientific support staff who provided manuscript feedback, especially Amit Frishberg and Nicola Rae Hennersdorf. This work was supported by funds from the Helmholtz Association, Helmholtz Munich, the German Center for Diabetes Research (DZD), the Deutsche Forschungsgemeinschaft (DFG, German Research Foundation, project number 458958943), and the Bavarian Ministry of Science and the Arts in the framework of the Bavarian Research Association “ForInter” (Interaction of human brain cells). F.J.T. acknowledges support by the Helmholtz Association’s Initiative and Networking Fund through Helmholtz AI [ZT-I-PF-5-01]. K.H. acknowledge financial support from the Joachim Herz Stiftung via Add-on Fellowships for Interdisciplinary Life Science and support from Helmholtz Association under the joint research school “Munich School for Data Science”. The human and mouse icons were adapted from publicly available images at vecteezy.com.

## Author contributions

Project conceptualization was performed by KH, FJT, HL, MBu, ABP, and MBa, data curation and computational analyses were performed by KH, laboratory analyses were performed by ABM, CS, MS, AB, and AM, visualizations were prepared by KH, the original draft was written by KH with the help of ABM and MBa and all authors reviewed the manuscript, supervision was provided by LZ, MBu, MBa, HL, and FJT.

## Abbreviations

DGE: -differential gene expression
DEG: -differentially expressed gene
ER: -endoplasmic reticulum
FACS: -fluorescence-activated cell sorting
FDR: -false discovery rate
GP: -gene program
GSIS: -glucose-stimulated insulin secretion
HVG: -highly variable gene
lFC: -log fold change
MIA: -mouse islet atlas
NOD: -non-obese diabetic
P: -postnatal day
PAGA: -partition-based graph abstraction
PC: -principal component
ROS: -reactive oxygen species
RT: -room temperature
scRNA-seq: -single-cell RNA sequencing
(m)STZ: -(multiple low dose) streptozotocin
T1D/T2D: -type 1 or 2 diabetes
UPR: -unfolded protein response
VSG: -vertical sleeve gastrectomy

## Supplementary materials

### Supplementary notes

#### Supplementary Note S1: Integration optimization

On the non-integrated embedding we observed strong separation between endocrine cell types across datasets (**Supplementary Figure S1a**). For example, γ-cells were located in two distinct regions across datasets. They did not even co-localize among in-house datasets; i.e. P16 and 4m datasets located separately from aged, mSTZ, and db/db datasets. As these differences could not be fully explained by biological differences after visual analysis of available metadata, we assumed that they are also driven by technical factors, which would bias our interpretations if we used the embedding without batch effect removal. Thus, we decided to perform integration, as recommended when analyzing multiple datasets at once^55,204,205^.

For integration of the atlas we tested cVAE model implemented in the scArches package^56^ and scVI method^54^ from scvi-tools, as it was recently reported that neural network based integration methods are most suitable for large and complex dataset collections, such as ours^55^. We observed stronger batch correction by scVI and comparable biological conservation on the level of cell types between cVAE and scVI integrations (**Supplementary Figure S1c**). As we were particularly interested in β-cell analysis we also evaluated integration on the level of β-cell states defined with known markers (**Supplementary Figure S1c**). Since scVI-based integrations had much lower biological variation preservation within β-cells we decided to use cVAE to enable exploration of biological heterogeneity within β-cells. We observed strong sample-specific ambient RNA contamination (**Supplementary Figure S1b**), possibly due to different cell type proportions across samples, introducing additional batch effects. Thus, we aimed to improve integration performance by preprocessing with various ambient-contamination removal methods. Interestingly, we found that preprocessing with ambient removal tools (DecontX^192^, SoupX^191^) did not improve integration beyond exclusion of genes with strongest ambient expression contribution, thus we decided to use ambient gene exclusion alone for the final integration (**Supplementary Figure S1c**).

Furthermore, we also attempted to specifically improve β-cell integration by performing integration of β-cells only. We assumed that by integrating only β-cells we could remove more of the ambient effects. That is, the HVG selection and thus the integration would no longer be affected by the variability of markers from other cell types. However, in the β-cell specific integration we were unable to achieve matched biological preservation at the same batch correction level as in the whole-islet integration (**Supplementary Figure S1d**), thus we did not use it further. Better integration of all cell types at once could be explained by different factors, such as a more diverse set of genes used for integration that improves biological variation preservation or by learning batch effects across cell types, thus being better able to separate technical effects from biological variation that may be more cell type specific. For example, in β-cells alone, some biological states are sample specific, resulting in direct confounding of batch and biological variation, while when using multiple cell types, the batch effects can be more easily disentangled from biological variability at the cell type level.

#### Supplementary Note S2: Expression of human diabetes-associated gene sets across mouse diabetes models

To compare mouse diabetes models with human diabetes we plotted for mouse data the expression of gene sets associated with human diabetes, including newly identified (see methods) and previously reported gene sets (**Figure 4e**, significance is reported in **Supplementary Table S4**). Gene set “transport vesicle membrane” was enriched in both our human T1D and T2D upregulated genes and was accordingly upregulated in all three mouse models. Genes upregulated in our human T1D DGE analysis were enriched for immune gene sets, as reported previously^30^, including “MHC class I protein complex” and “antimicrobial humoral response”. Both of these gene sets had a larger fold change between healthy and diabetic β-cells in the NOD than in the mSTZ or db/db. Among the gene sets enriched in our newly identified human T2D upregulated genes were gene sets associated with regulation of response to extracellular stimuli and with hormone metabolic processes, both of which were upregulated in the db/db and mSTZ models, but not in the NOD model. We further supplement our analysis with gene sets previously reported to be upregulated in diabetic β-cells^30,31,103^, including gene sets indicative of β-cell compensation in T2D^30,103^. These gene sets (oxidative phosphorylation, ribosome biogenesis, regulation of cellular response to hypoxia, and proteasomal protein catabolic process) were upregulated in db/db and mSTZ, but not NOD diabetes model β-cells. In **Supplementary Note S3** we discuss the limitation of our gene set comparison when the gene set contains too diverse genes that are associated with opposing molecular processes.

#### Supplementary Note S3: Limitation of cell group comparison with gene set scores

To compare molecular characteristics of diabetes in human and mouse models we collected gene sets upregulated in human diabetes followed by comparison expression of these gene sets in mouse healthy controls and diabetic samples. While this method produced expected results in mouse diabetes models for most gene sets (explained in **Supplementary Note S2**) we observed some unexpected discrepancies between the two species and also current literature. The gene set “endocrine pancreas development” was enriched among human T2D-upregulated genes (**Supplementary Table S4**), which raises the expectation that its genes in mouse would also have higher overall expression in the mouse T2D models compared to healthy controls (**Figure 4e**). However, the gene set was downregulated in the db/db and mSTZ models (T2D) and not significantly dysregulated in the NOD model (T1D). This is not in agreement with current literature as T2D related dedifferentiation was before observed in both mouse and human^31,32,40^. The unexpected direction of the gene set change in the db/db and mSTZ models may be explained by the gene set “endocrine pancreas development” containing both maturation (e.g., *Pax6*, *Pdx1^32^*) and immaturity (e.g., *Neurod1*, *Neurog3^85^*) genes, which likely affects the score in the mouse cells. This shows a limitation of our gene set based analysis when the selected gene set contains too diverse genes.

#### Supplementary Note S4: Description of coarse β-cell states and their markers

We resolved seven coarse states based on metadata information as well as two clusters that are likely low quality populations, characterized by poor quality metric scores (**Supplementary Figure S7a**), which we regarded as technical artifacts and hence did not use in the downstream β-cell coarse state marker identification. Most of the cell states contained cells from multiple datasets (**Supplementary Figure S7c, Supplementary Table S2**). Due to the high transcriptomic similarity between mSTZ and db/db diabetic β-cells, as described above, they were annotated as a single T2D model state. The immature state contained cells ranging from P16 to adult, but was characterized by high expression of known immaturity markers, such as *Rbp4* and *Mafb^104^* (**Figure 5b**). Two healthy datasets of different ages contained mixed sex samples (“P16” - postnatal day 16, “aged” - 2 years old). Separation of cells by sex was observed only in the aged dataset (**Supplementary Figure S7f**), as expected since the P16 dataset was generated from sexually immature cells before puberty. Furthermore, we observed that β-cells from the chem dataset mapped separately from other datasets (**Figure 4a**, **Figure 5a**, **Supplementary Figure S7c**), likely due to *in vitro* islet cultivation or ambient effect of the spike-in RNA used in the protocol^38^.

We used MIA to identify new markers that are both specific for an individual β-cell state and conserved across all datasets mapping to that state by applying one-vs-rest DGE between cell states and retaining the intersection of markers across datasets (**Figure 5c**, **Supplementary Figure S8a**, **Supplementary Table S5**), with each state containing one to four datasets (**Supplementary Figure S8b**). We observed a low number of cell-state markers for some states, such as adult and immature states (**Supplementary Figure S8b**). This may be due to the stringent requirement of markers being consistent in four datasets mapping into these states, however, this ensures high robustness. We also observed on a per-dataset level that certain cell states, including the adult state, had fewer markers, indicating that these states are less molecularly unique (**Supplementary Figure S8b**). Reliable detection of these states would likely benefit from a combination of positive and negative markers. However, we found a large number of db/db+mSTZ state markers that were conserved across both datasets mapping into this state (**Supplementary Figure S8b**), indicating the strength of molecular changes involved in the T2D model dysfunction.

#### Supplementary Note S5: Comparison of β-cell heterogeneity within MIA with previous literature

As β-cell heterogeneity has been extensively described before^14^, we decided to compare our results to key publications in the field. Gradual maturation of β-cells after birth was previously characterized based on upregulation of *Ucn3* and downregulation of *Mafb* by Zeng et al. (2017) and based on downregulation of *Cd81* by Salinno (2021). Accordingly, we observed higher expression of *Mafb* and *Cd81* and lower expression of *Ucn3* in our immature state (imm.) from younger mice compared to healthy adult state (adult, **Figure 5b, Supplementary Figure S7c**). Furthermore, our GP23, which contained multiple immaturity markers (*Rbp4*, *Mafb*), was gradually downregulated from immature to adult states (imm.1 and imm.2, adult-imm.1, and adul1 and adul2). However, multiple genes that were reported to be downregulated during maturation by Zeng et al. (2017) (*Fos*, *Fosb*, *Egr1*, *Junb*, *Jun*, *Txnip*) were present in our aging-associated healthy gene group 4 (**Supplementary Table S8**), showing some discrepancy between exact maturation-associated genes. It was also reported that proliferative β-cells decline after birth^206^ and indeed, we observed the highest proportion of proliferative endocrine cells in our youngest postnatal dataset (P16, **Supplementary Figure S3**). Moreover, Aguayo-Mazzucato et al. (2017) reported increase of senescence marker *Cdkn2a* during aging, as was also observed in our aged states (agedF and agedM, **Figure 5b**). Maturity and aging heterogeneity were previously also observed within individual samples^206,207^. This corresponds to our finding that all healthy adult samples contain subpopulations of cells with high expression of immaturity or aging-related genes (healthy variable gene groups 2 and 4, **Supplementary Note S7**). However, our integrated embedding did not show separation between cell populations sorted for maturity marker Flattop (**Supplementary Figure S6**, datasets: P16, 4m, aged), which was discovered by Bader et al. (2016)^20^ and similarly we did not observe a strong difference in *Cfap126* (Flattop gene) mRNA expression between sorted populations (**Supplementary Figure S7g**). This is expected as *Cfap126* is expressed only transiently and at low levels, while both of the reporters mark the cells even after *Cfap126* is no longer expressed^174,175^.

Xin et al. (2018) reported cycling of cells between states of active insulin production, resulting in metabolic stress, followed by recovery phase. We observed similar metabolic heterogeneity in healthy adult samples, as described in the section on conserved healthy heterogeneity gene groups.

Oppenländer et al. (2021) and Sachs et al. (2020), whose data is contained within MIA, described db/db and mSTZ model populations, respectively, distinct from healthy controls. Their diabetic model populations match our db/db+mSTZ population (**Supplementary Figure S7c,d**) and we detected molecular changes corresponding to previous publications (**Supplementary Note S6**). Similarly, markers *Ucn3* and *Slc2a2* were reported to be downregulated in the STZ model by Feng et al. (2020) and were also downregulated in our diabetic db/db+mSTZ population (**Figure 5b**). However, *ST8SIA1*, which was reported to be upregulated in human T2D by Dorrell et al. (2016), was significantly downregulated in our db/db+mSTZ DGE analysis (**Supplementary Table S10**). Conversely, their marker *CD9*, for which they did not discuss association with diabetes^208^, was upregulated in our db/db+mSTZ DGE analysis. Our integration also preserved finer cellular heterogeneity. The trajectory between mSTZ model cells with and without anti-diabetic treatment and controls, which was reported in the original study^40^, was also observed in MIA (**Supplementary Note S8**). Similarly, Oppenländer et al. (2021) reported differences between db/db model cells with and without VSG and we observed separation of VSG treated cells into a distinct cluster (db/db-VSG state, **Supplementary Figure S7d**). Thompson et al. (2019), whose data is also contained within MIA, described a distinct cell population during NOD model diabetes progression, matched to our NOD-D population (**Figure 5a, Supplementary Figure S7c, Supplementary Note S6**). As expected, based on their reports, our DGE analysis of the NOD model revealed upregulation of senescence markers *Cdkn1a* and *Cxcl10* (**Supplementary Table S10**).

While we largely observe correspondence with previous studies in terms of markers and subpopulations, we also report some results that we could not reproduce. Having a comprehensive atlas that enables quick lookup of genes and cells, such as MIA, thus helps to reveal findings that do not translate to other datasets.

#### Supplementary Note S6: Comparison of healthy and diabetic states with GPs

To showcase our GP-based interpretation approach we applied it to cell states that have been compared before^32,36,40^. Namely, we compared the main healthy state (adult2) to the main T2D model state (db/db+mSTZ) for the db/db and mSTZ datasets and to the T1D model state (NOD-D) for the 8-16wNOD dataset (**Supplementary Figure S10c**). We selected the adult2 and db/db+mSTZ states as the main healthy and T2D model states, respectively, based on them containing cells across datasets for specific conditions. According to our previous observations in the diabetes models comparison, the GP differences were similar for db/db and mSTZ diabetes model cells and more distinct for the NOD model (**Supplementary Figure S10c**). Among the most differentially activate GPs in the diseased state of the db/db and mSTZ models were downregulated GP19, enriched in genes related to insulin or peptide hormone secretion, GP20, enriched in genes associated with stimuli (growth factors, hormones, cytokines) response, and GP17, enriched for p38 MAPK cascade. On the other hand, the diseased state of the db/db and mSTZ models exhibited upregulation of GP3, which is predominantly active in the T2D-model cells (**Supplementary Figure S10b**) and contains known T2D-associated genes (*Cck*, *Gc*, *Slc5a10*, and *Gast*)^32^, and GP4, enriched in protein processing and golgi-ER endomembrane system. In the NOD diabetic state, we observed the strongest change in upregulation of GP27 (enriched for antigen presentation and interferon response) and GP25 (enriched for response to viruses). In all comparisons of diabetes model states there was downregulation of GP15, enriched for circadian clock, insulin secretion, and stimuli response genes. Altogether, this shows that GPs provide relevant insights into differences between states, which can be broadly summarized by immune-mediated β-cell dysfunction in the NOD T1D model and metabolic/stress-related dysfunction in the db/db and mSTZ T2D models^102^.

#### Supplementary Note S7: Description of gene groups consistently variable in healthy samples

Gene groups consistently variable across health samples (**Supplementary Table S8**) were interpreted based on gene set enrichment and group member genes (**Supplementary Table S8**), correlation with known β-cell function genes (**Figure 5g**), and expression across MIA β-cells (**Supplementary Figure S11b,c**), as summarized in **Supplementary Table S8**. Groups 3 and 5 had similar expression patterns in healthy cells (**Supplementary Figure S11a**) and were associated with β-cell maturity and insulin production, with group 3 being more strongly associated with insulin-production related stress and more strongly active in diabetes model states (**Figure 5g, Supplementary Figure S11b**, **Supplementary Table S8**). Group 1, which was highly active in cells with low activity of groups 3 and 5 (**Supplementary Figure S11a**), was negatively correlated with expression of many insulin-production related genes and contained genes previously implicated in β-cell recovery from metabolic stress, such as ATP production related genes^107^, and protective genes (**Figure 5g, Supplementary Table S8**). Group 2 contained immaturity genes (**Figure 5g**), and was more strongly expressed in immature states of MIA and in healthy adult samples with higher proportion of immature cells (**Supplementary Figure S11b,c, Supplementary Figure S7c**). Its high activity in healthy adult cells was mutually exclusive with high activities of groups 1, 4, and 5, and to a lesser extent of group 3 (**Supplementary Figure S11a**), indicating a presence of a distinct immature population in healthy adults. Group 4 contained stress and senescence associated genes (**Figure 5g, Supplementary Table S8**) and had increased activity in the aged and more mature cells from MIA (**Supplementary Figure S11b,c**).

We assessed the localization of cells with the highest expression of individual gene groups in healthy adult samples of the atlas and Feng dataset mapped onto the atlas (**Supplementary Figure S11d**). We observed that cells with high expression of individual gene groups localized to distinct regions of the UMAP embedding, witch cells high in groups 3 and 5 showing somewhat greater overlap, as expected based on expression similarity of these two gene groups (**Supplementary Figure S11a**). In all samples cells from the aged-like gene group (4) co-localized near the aged cells (agedM β-cell coarse state), confirming the relevance of this gene group in aging. In general, group 1 (low insulin expression) high-scoring cells localized somewhat further from diabetes model cells than cells high in groups 2, 3, and 5. However, the exact regions of the group-high cells differed between 4m dataset and mSTZ and db/db datasets, as expected due to different localization of these datasets in the integrated embedding. Especially in the mSTZ, db/db, and Feng datasets we observed a large number of group 3 and also group 5 high cells that localized near the diabetic intermediate cells (D-inter. fine state), confirming the association between stressed or diabetic-like phenotype and increased insulin production. Furthermore, all datasets contained gene group 2 high cells that co-localized with imm.3 and mSTZ fine states. In Feng et al. (2020) it was proposed that a distinct β-cell population within their dataset represents “virgin” β-cells that are located in the islet periphery, but whose function is not fully understood^209^. This “virgin” population within the Feng dataset corresponds to our gene group 2 high cells within imm.3 and mSTZ fine states (**Supplementary Figure S11d,e**). Similarly, gene group 2 activity is anticorrelated to genes downregulated in “virgin” β-cells (*Ucn3*, *Ero1b, G6pc2, Mafa*) and correlated to genes upregulated in “virgin” β-cells (*Mafb* and other immaturity markers, **Figure 5g**)^210,211^. Interestingly, this indicates similarity between “virgin” and a subset of mSTZ-model β-cells. Additionally, the db/db dataset also contained some gene group 2 high cells that mapped near the immature coarse state (imm.) and the gene group 2 high cells from the 4m dataset likewise mapped to both mSTZ-model and immature regions, indicating the presence of different immature populations in some of the healthy adult samples.

#### Supplementary Note S8: Trajectories used in diabetic DGE analysis

Dysfunctional NOD samples were relatively young (14 and 16 weeks) and thus the full dysfunctional phenotype might not yet have been developed^29^. Nevertheless, a recent study of T1D and autoantigen-positive prediabetic human donors revealed that pre-diabetic samples already show similar transcriptomic changes to those with progressed T1D^53^, indicating that despite the youth of our NOD samples they may nevertheless provide valuable insights into diabetes dysfunction.

A gradient of cells from healthy to dysfunctional was observed in both NOD and db/db+mSTZ mice, which we modeled as a linear trajectory based on localization of samples and their metadata using MELD^202^ (**Supplementary Figure S13a**). We confirmed the validity of both fitted trajectories. In NOD cells the trajectory corresponded to *B2m* expression, a known marker of T1D^212^. In db/db+mSTZ trajectory the cells from diabetes model animals receiving anti-diabetic treatment (VSG and insulin administration) had lower dysfunction scores^32,40^ (**Supplementary Figure S13a**).

#### Supplementary Note S9: Groups of DEGs in T2D model support separation of mSTZ model cells into two distinct populations

We described that db/db and mSTZ models co-localize on the atlas embedding and show their molecular similarity, however, we also identified heterogeneity within the mSTZ model. Similarly, in our T2D model DGE analysis, some DEG groups were differently strongly dysregulated across T2D model states, while still showing differences against healthy adult cells (**Figure 6a**). The mSTZ cell state had higher activity of the gene group T2-up1 that was also highly expressed in immature cells. In contrast, db/db+mSTZ cell state had higher activity of T2-up2 that was lowly expressed across healthy cells and was enriched for transmembrane ion transport and cell-cell adhesion, which were previously associated with insulin secretion and diabetic compensatory response^30,46^. Cell state db/db+mSTZ also had stronger downregulation of T2-down6 that contained genes involved in key β-cell functions (**Supplementary Table S10**) and whose downregulation was previously linked to β-cell stress^114,213–215^. This is in accordance with the differences between the two states that we described above based on GPs, with the db/db+mSTZ state showing stronger signs of β-cell metabolic stress than the mSTZ state.

#### Supplementary Note S10: Validation of cells states in islets used for antibody analysis

We validated that the islets used for antibody-based validation of diabetes markers contain the expected healthy and dysfunctional cell states in health and diabetes models, respectively. As expected, insulin positive cells were reduced in both diabetic models (NOD, db/db) compared to control. Ucn3 was also strongly downregulated in the db/db model and to a lesser extent in the NOD model compared to healthy control on both the protein and RNA level. Aldh1a3 protein positive cells were observed only in the diabetes models (NOD and db/db) and not in the healthy control, however, we did not observe upregulation at the RNA level in NOD islets.

#### Supplementary Note S11: Gene program differences between the healthy and the intermediate state resemble changes in diabetes

To decipher what distinguishes the intermediate state (D-inter.) from the main healthy state (adult2) we again used GP score differences for each individual diabetes model dataset (**Supplementary Figure S10e**). The GP differences exhibited a pattern similar to the differences between healthy and diabetes model states (**Supplementary Figure S10c**), but with a lower magnitude. Relatively strong differences were also observed in some GPs with similar changes across all diabetes models. Examples are D-inter. downregulated GP15 (enriched in insulin secretion, stimuli response, and circadian clock (**Supplementary Table S6**)) and GP23 (characteristic for immature samples and enriched in cholesterol metabolism, cell junctions, and response to metal ions).

### Supplementary figures

**Supplementary Figure S1:**
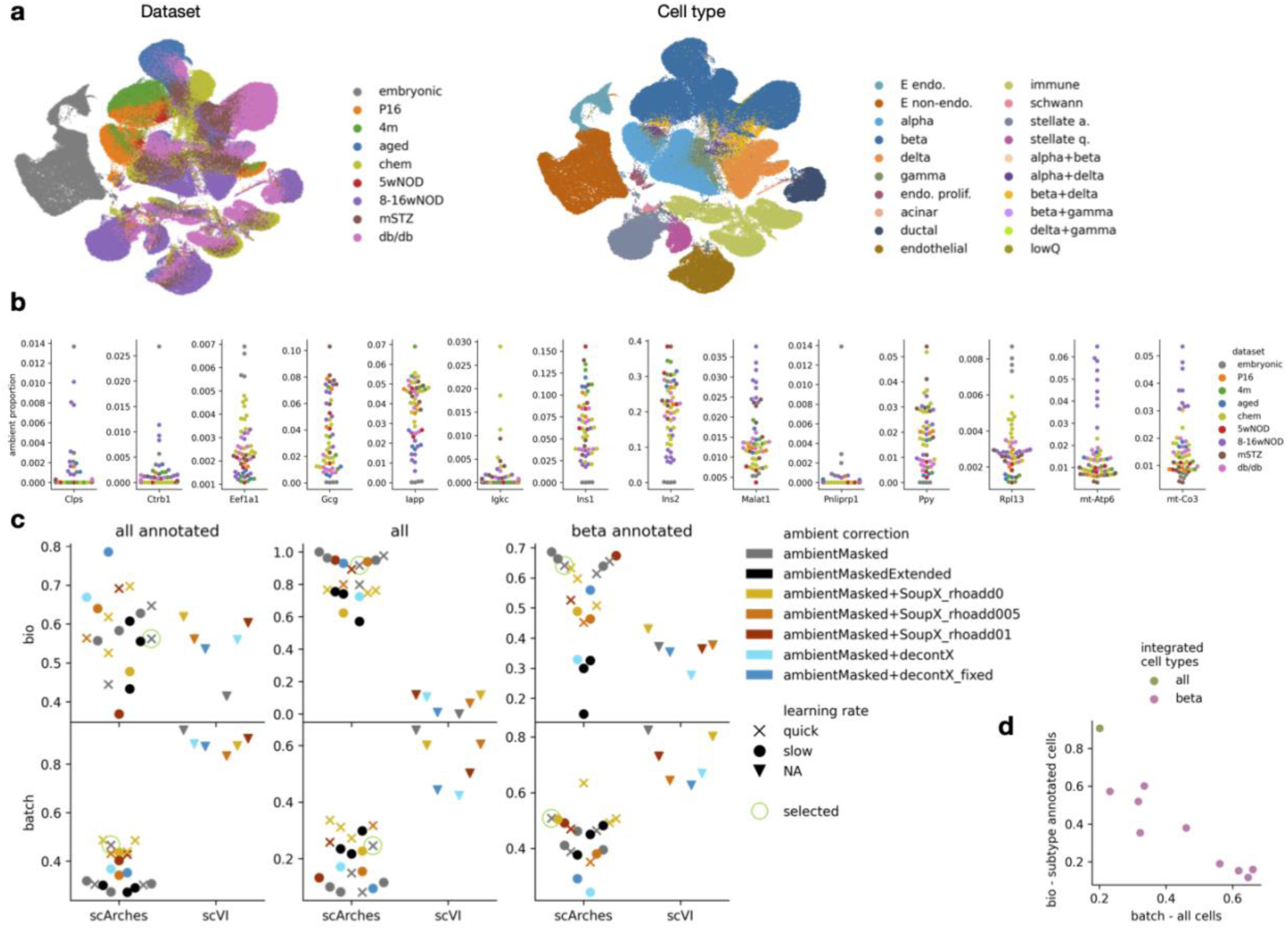
Evaluation of atlas integration. (a) UMAP embedding of unintegrated atlas data. Cell types are colored based on annotation from integrated embedding for comparability to the final integrated atlas (**Figure 2d,e, Supplementary Figure S2a**). (b) Ambient expression strength of most ambient genes across samples. Ambient proportion denotes proportion of counts within empty cells that belong to the gene. Each dot represents a sample colored by dataset. (c) Integration metrics results for different integration protocols (dots) showing biological conservation (bio) and batch removal (batch) scores in scArches-cVAE and scVI integrations. Evaluation was performed on different data subsets: all annotated - cells from all cell types that had cell type annotation, which was used for evaluation (**Supplementary Figure S2b**); all - all cells, including unannotated, for batch removal evaluation we used clusters instead of cell types and for biological preservation evaluation we used only Moran’s I; beta annotated - cells annotated as β (**Supplementary Figure S2b**), using marker-based cell states for evaluation. Some integration protocols were run multiple times, indicated by multiple dots of the same color and shape. Encircled integration is the one used for the atlas. We preprocessed data before integration with different ambient removal methods, including removal of top ambient genes (ambientMasked) or an extended list of ambient genes (ambientMaskedExtended), SoupX using default rho (rhoadd0) or increased rho (rhoadd005 - adding 0.05, rhoadd01 - adding 0.1), and DecontX using estimated delta or a pre-set delta (fixed) to increase the strength of ambient removal. We ran scArches- cVAE with two learning rate settings, as described in the methods. (d) Integration metrics for β-cell specific integrations compared to metrics computed on β-cell embedding from the integration of all cells. We tuned β-cell integration strength, corresponding to multiple points on the graph.

**Supplementary Figure S2:**
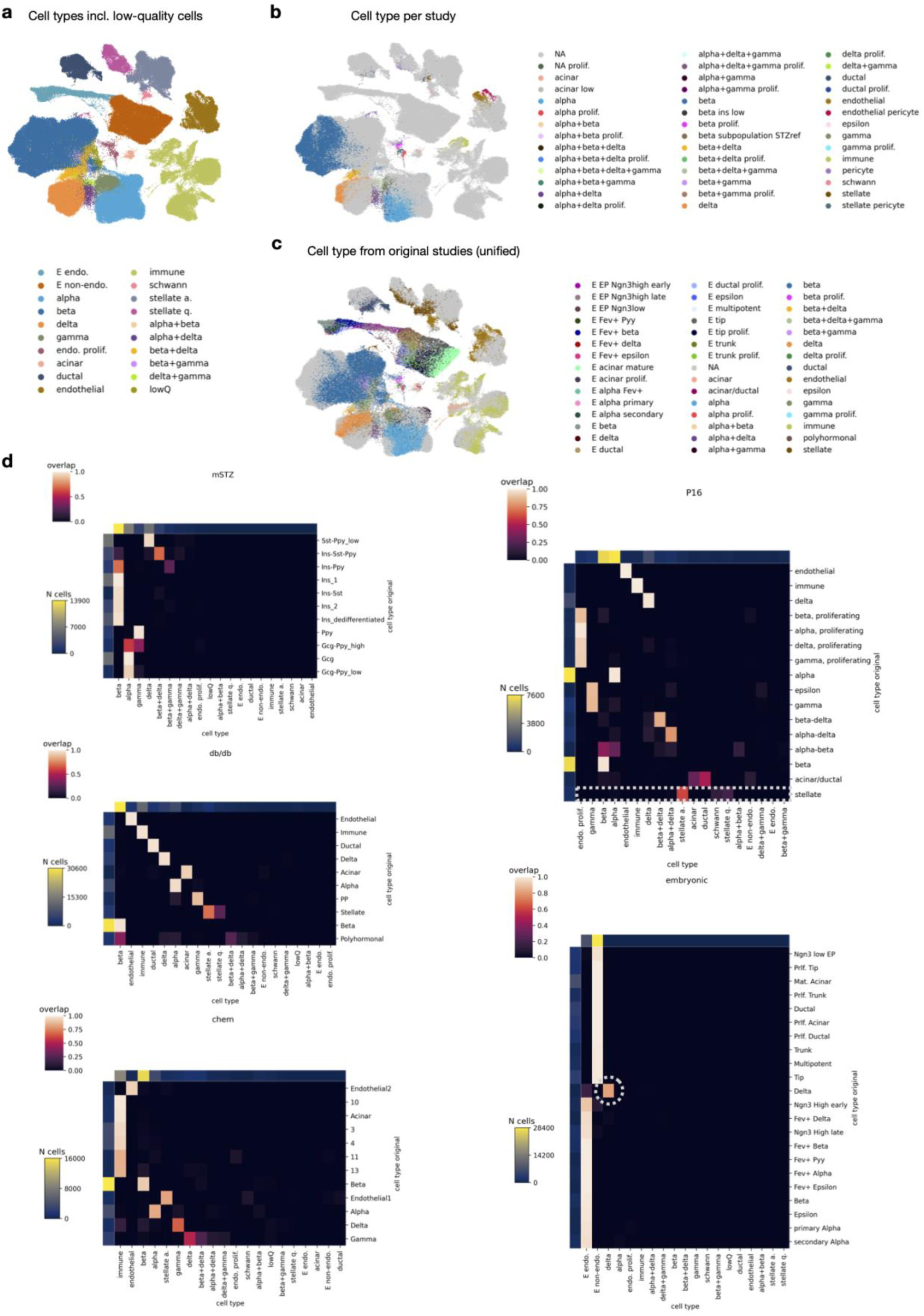
Comparison of cell types assigned in original studies and in integrated atlas re-annotation. (a) Atlas-level cell type re-annotation within atlas shown on UMAP, including low quality and potential doublet cells. (b) Cell types used for integration evaluation. Annotation was performed for selected samples (colored in cells) per-study; unannotated cells are marked with NA. Some cell types were later renamed for the final atlas annotation (e.g., the annotations in panel b contain the name pericyte which was later in panel a corrected to stellate activated). (c) Cell types as reported in the original publications. Cell type names were unified across studies and cells with missing annotation are marked with NA. (d) Comparison of integration-based re-annotated and previously reported cell type labels. Datasets that did not have previously reported annotation are not shown. Overlaps were normalized per previously-reported cell type. In the P16 dataset the dotted rectangle indicates rare schwann cells that were merged with a larger population of stellate cells in the original annotation. In the embryonic dataset the dotted circle indicates the mapping of embryonic δ-cells to the postnatal δ-cells cluster. Abbreviations: “+” - potential doublet, lowQ - low quality, EP - endocrine progenitor/precursor, prlf. - proliferative, mat. - mature.

**Supplementary Figure S3:**
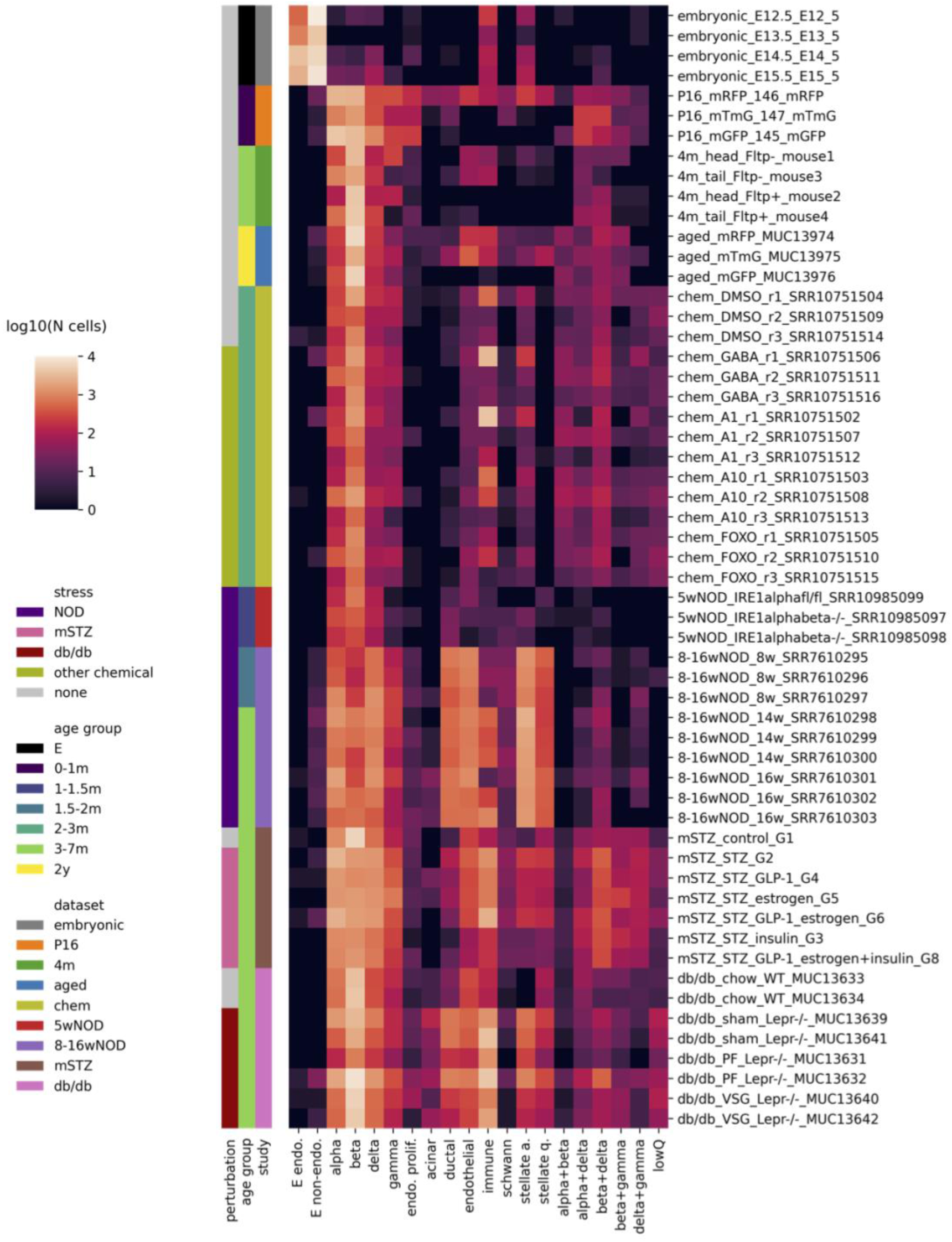
Number of cells per cell type in each sample. Sample names are given as study_sampleDescription_sampleIdentifier. Some of the datasets contained samples enriched for endocrine cells (**Supplementary Table S1**), which prevents direct cell type proportion comparison between samples with different cell sorting.

**Supplementary Figure S4:**
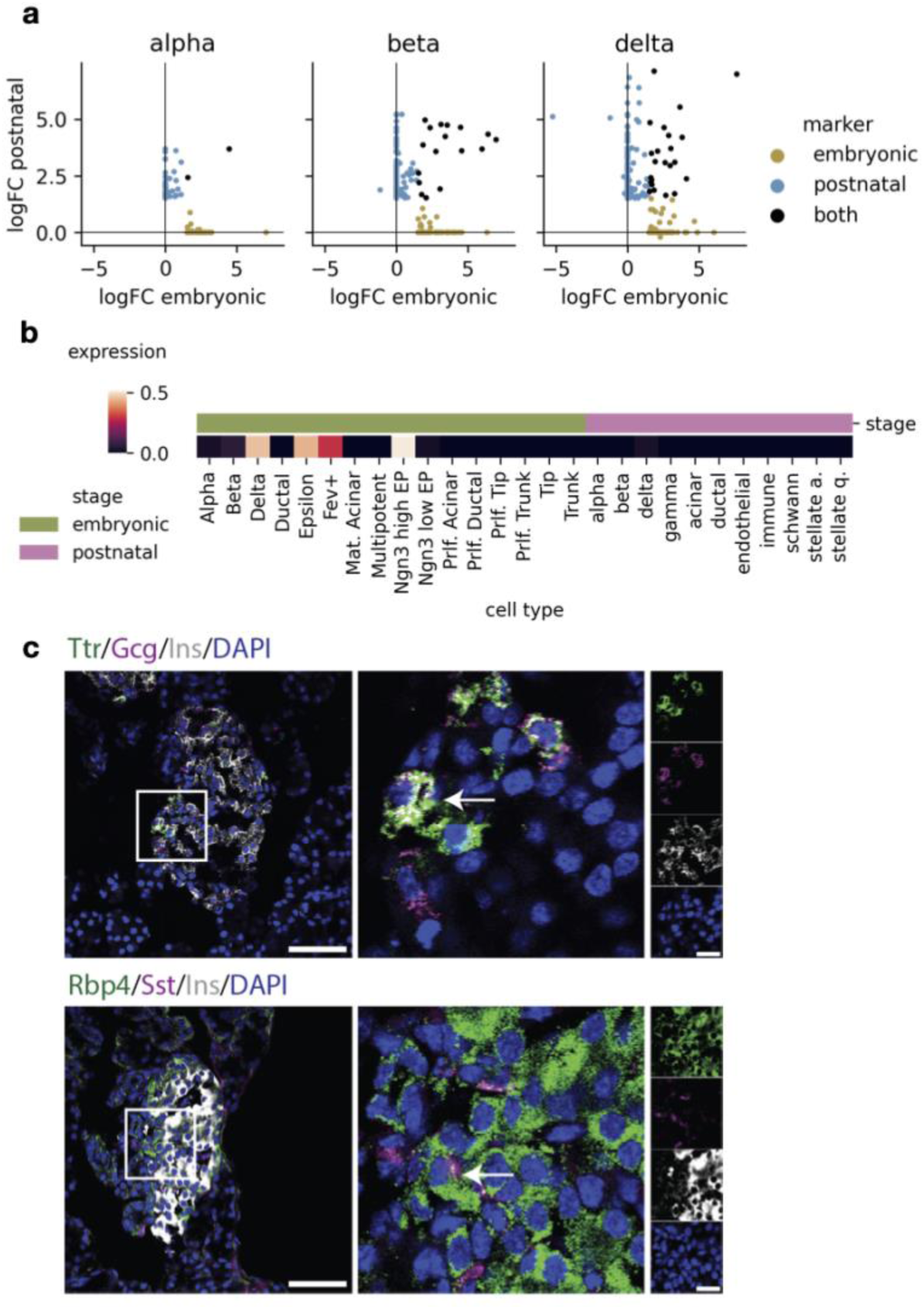
Endocrine markers differ in embryonic and postnatal datasets. (a) Comparison of endocrine markers in embryonic and postnatal data, showing whether genes were selected as potential markers in each stage (color). Genes missing from a stage-specific DGE analysis were assigned a logFC of 0. (b) Expression of *Cer1* across embryonic cell types (original study annotation) and postnatal cell types (atlas-level re-annotation). (c) Validation of selected endocrine markers with immunohistochemistry. Arrows indicate Ttr and Gcg double-positive α-cells (top) and Rbp4 and Sst double-positive δ-cells (bottom).

**Supplementary Figure S5:**
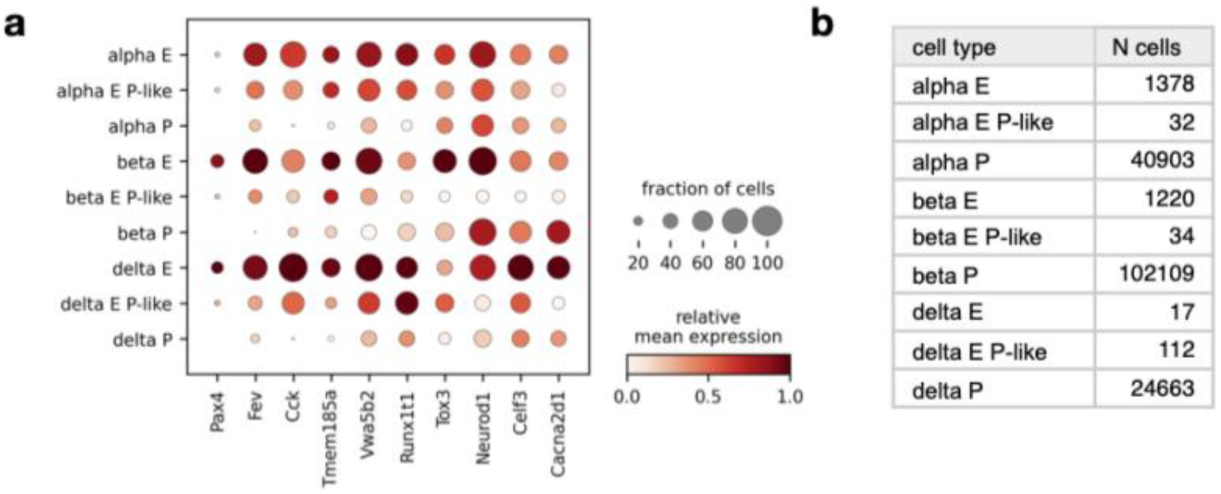
Endocrine cell groups in embryonic and postnatal datasets show gradual decrease in endocrine precursor markers. (a) Expression of embryonic Fev+ EP markers from Bastidas-Ponce et al. (2019) across endocrine cell groups. (b) Number of cells in each endocrine cell group. Cell groups are as in **Figure 3**.

**Supplementary Figure S6:**
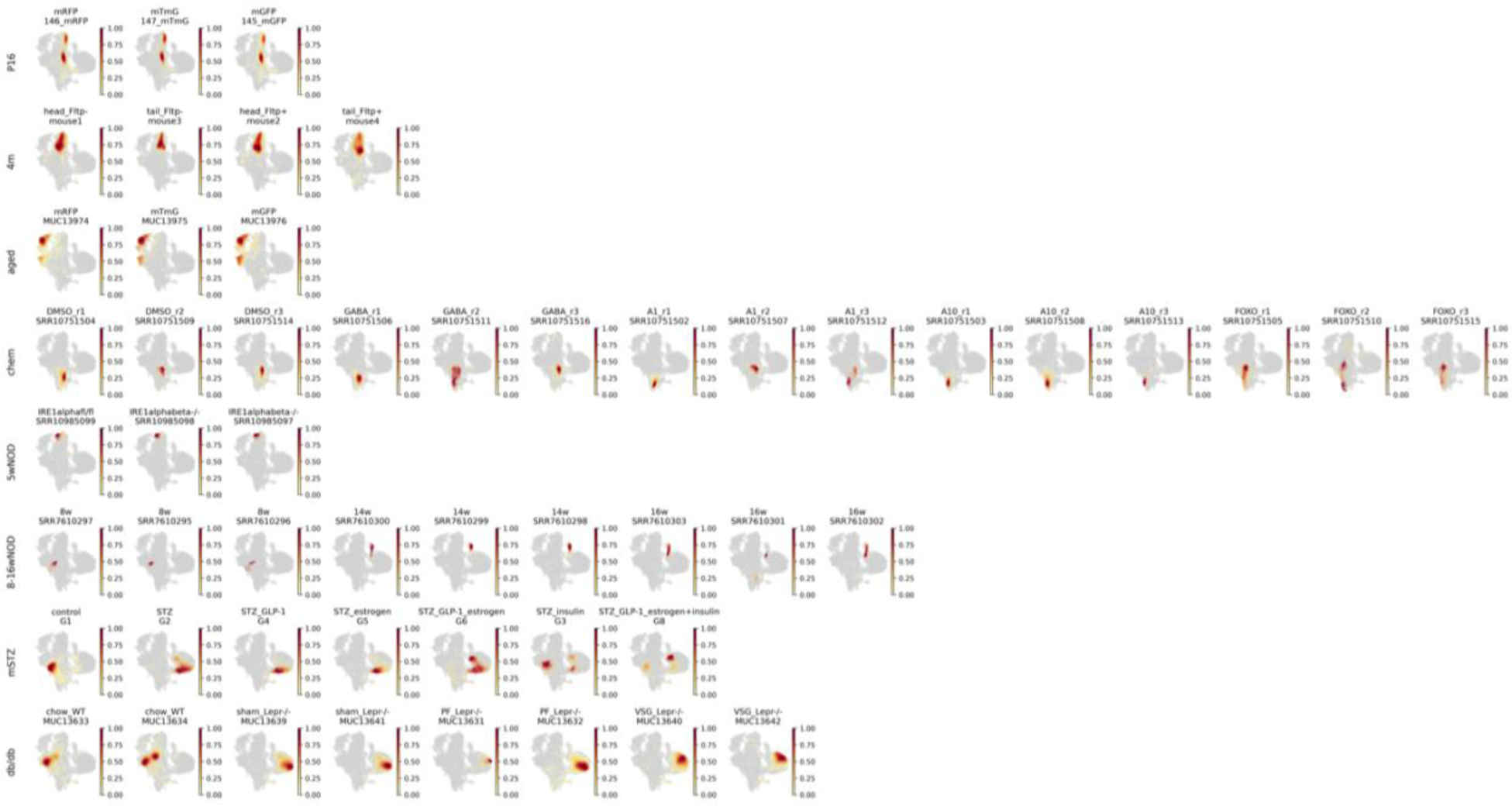
Integrated embedding of β-cells from individual samples corresponds to biological conditions. Distribution density of cells from each sample on a UMAP of the β-cell atlas subset. Sample names are reported with sample description and identifier. Embryo dataset is not shown due to a small number of cells within the β-cell cluster.

**Supplementary Figure S7:**
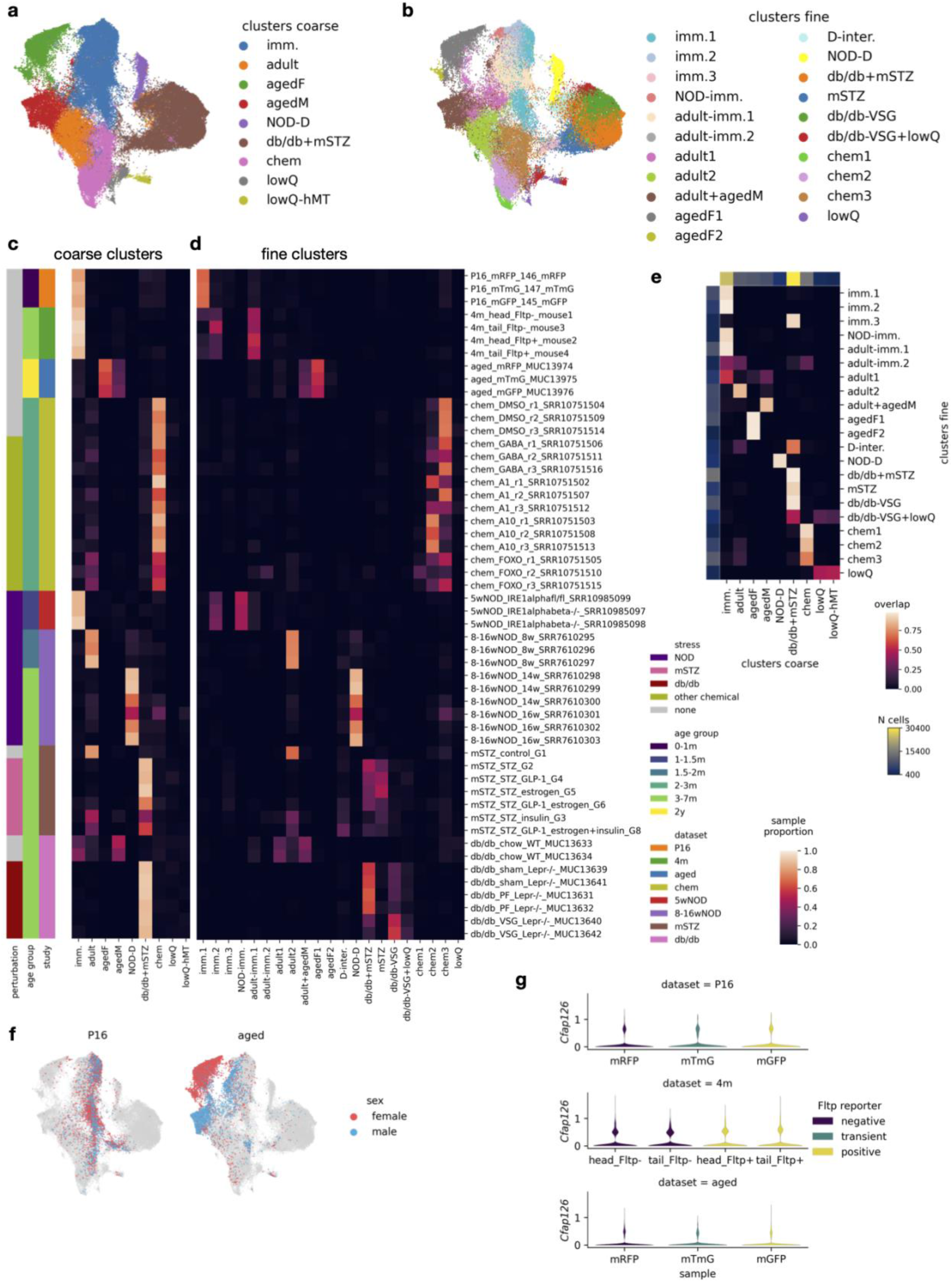
Different resolution β-cell states correspond to biological conditions. (a) Coarse β-cell states, including low quality clusters. (b) Fine β-cell states, including low quality clusters. (c) and (d) Coarse and fine, respectively, β-cell state proportions in each sample, also displaying corresponding sample metadata. Sample names are given as study_sampleDescription_sampleIdentifier. (e) Comparison of coarse and fine β-cell state annotation given as a normalized distribution of each fine state across coarse states. (f) A UMAP embedding of male and female β-cells across ages. Colored in are cells from datasets that have mixed sexes within samples, other β-cells are displayed as a background. (g) Expression of *Cfap126* (Flattop gene) across cell populations that were sorted based on the Flattop reporter system. Abbreviations: hMT - high mitochondrial transcript read fraction.

**Supplementary Figure S8:**
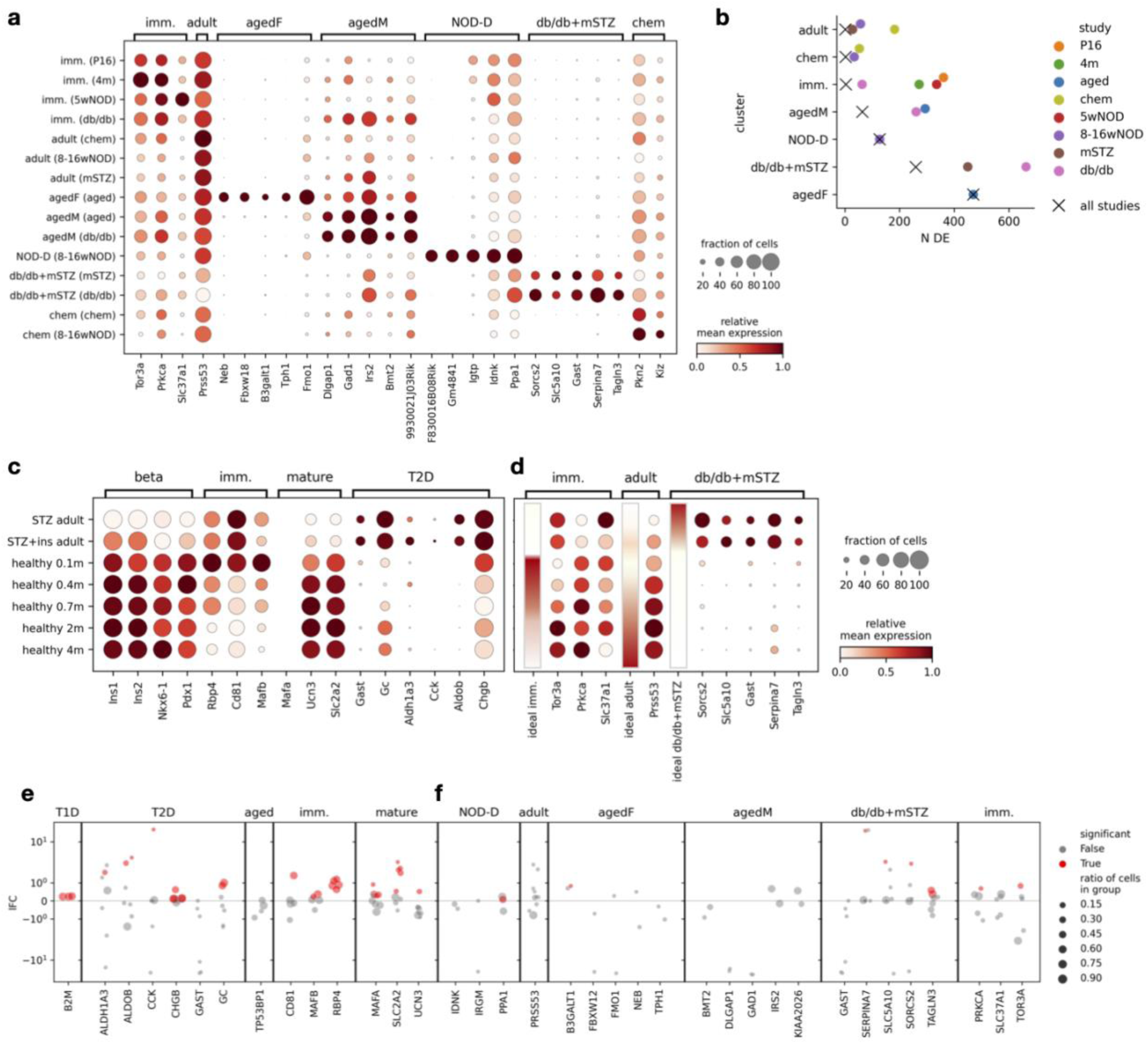
Newly defined β-cell markers are robust across mouse datasets, but do not directly translate to human. (a) Expression of proposed β-cell state markers across coarse β-cell states per dataset (in brackets). (b) Number of cluster specific markers extracted per dataset and state or as intersection of all datasets within a state. (c) and (d) Expression of known and newly proposed, respectively, β-cell state markers on the external Feng mouse dataset. Ideal marker bar represents how we would expect markers of specific clusters to be expressed across the cell groups. (e) and (f) Translation of known and newly proposed, respectively, β-cell state markers to human datasets. In each dataset (dot) we compared marker expression within the relevant sample group to all other samples, showing comparison lFC and statistical significance as well as the ratio of cells expressing the gene in the target group. Abbreviations: ins - insulin.

**Supplementary Figure S9:**
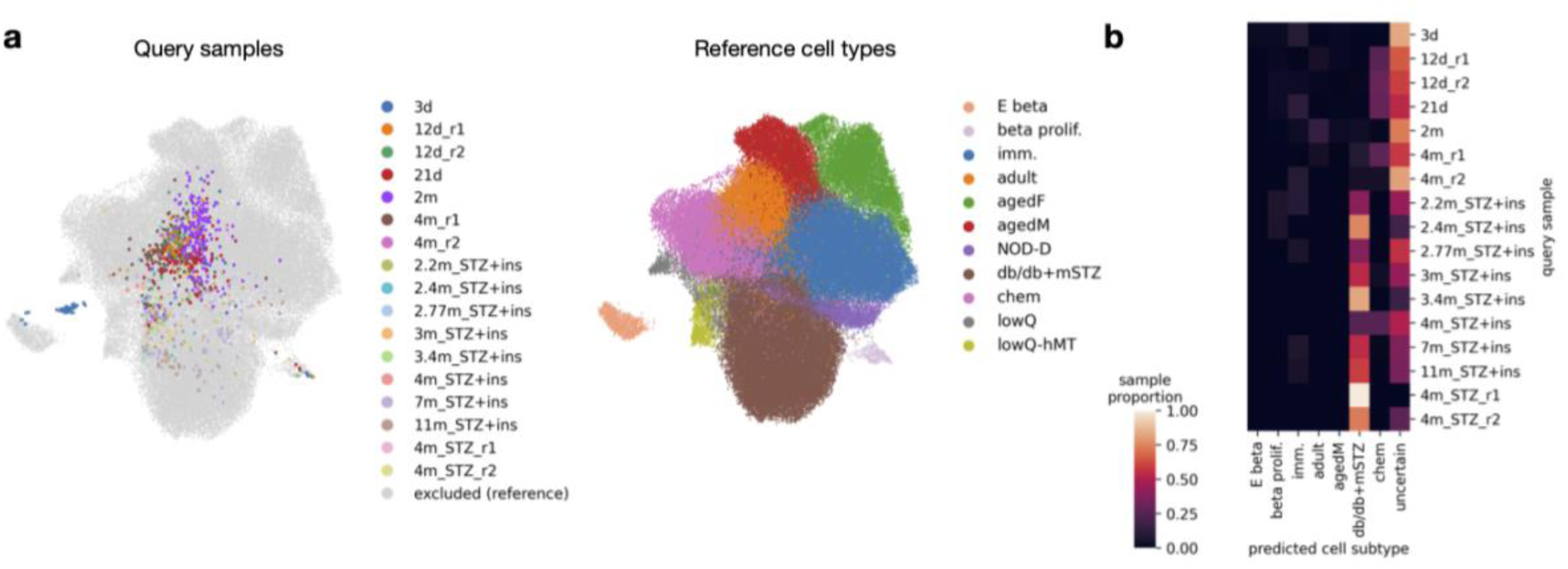
Reference mapping and cell state transfer for external mouse dataset β-cells reveals expected reference and query sample relationships. (a) Joint embedding of the atlas (reference) and the external mouse dataset (query). Left: All query samples, named as age, treatment, and replicate when multiple samples with the same age and treatment were present. Right: Reference cell groups showing coarse β-cell states, as shown in **Figure 5a**, and proliferative (part of the “endocrine proliferative” atlas cluster) and embryonic β-cells (as annotated in the original study of the embryonic dataset). (b) Label transfer from atlas to query, using cell groups as described in a. Cells with low label-transfer probability were assigned to the uncertain group. Shown are ratios of each query sample predicted as a certain cell group.

**Supplementary Figure S10:**
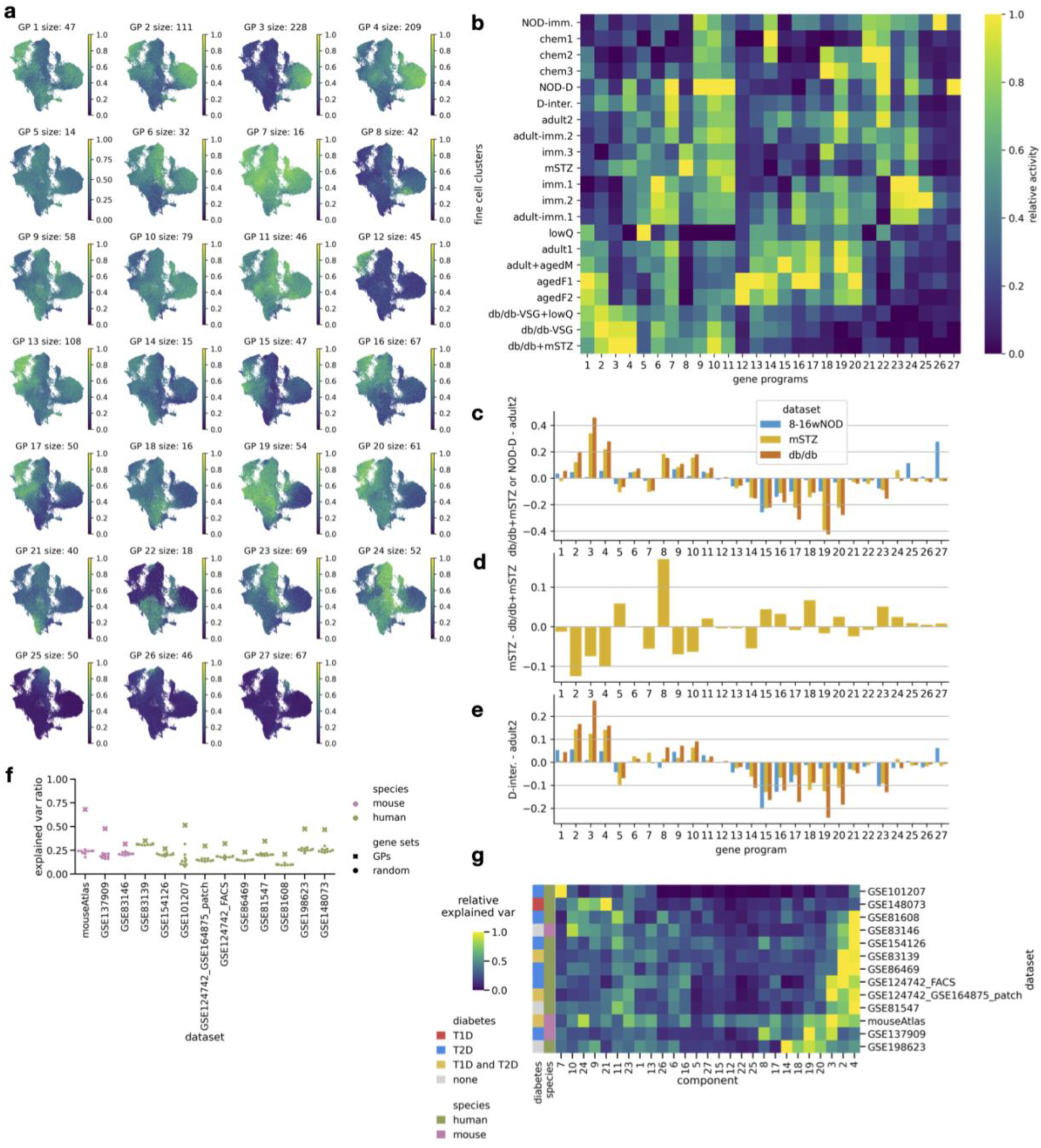
Activity of GPs across β-cells helps in β-cells state interpretation. (a) Activity of GPs on UMAP of the β-cell atlas subset. (b) Activity of GPs across fine β-cell states normalized per GP across states. (c), (d), (e) Differences in GP activity between pairs of fine β-cell states (specified on y axis) for individual datasets. (f) Ratio of variance explained by GPs across datasets compared to random groups of genes. (g) Relative variance explained by each GP, scaled as a ratio of maximal absolute value per dataset.

**Supplementary Figure S11:**
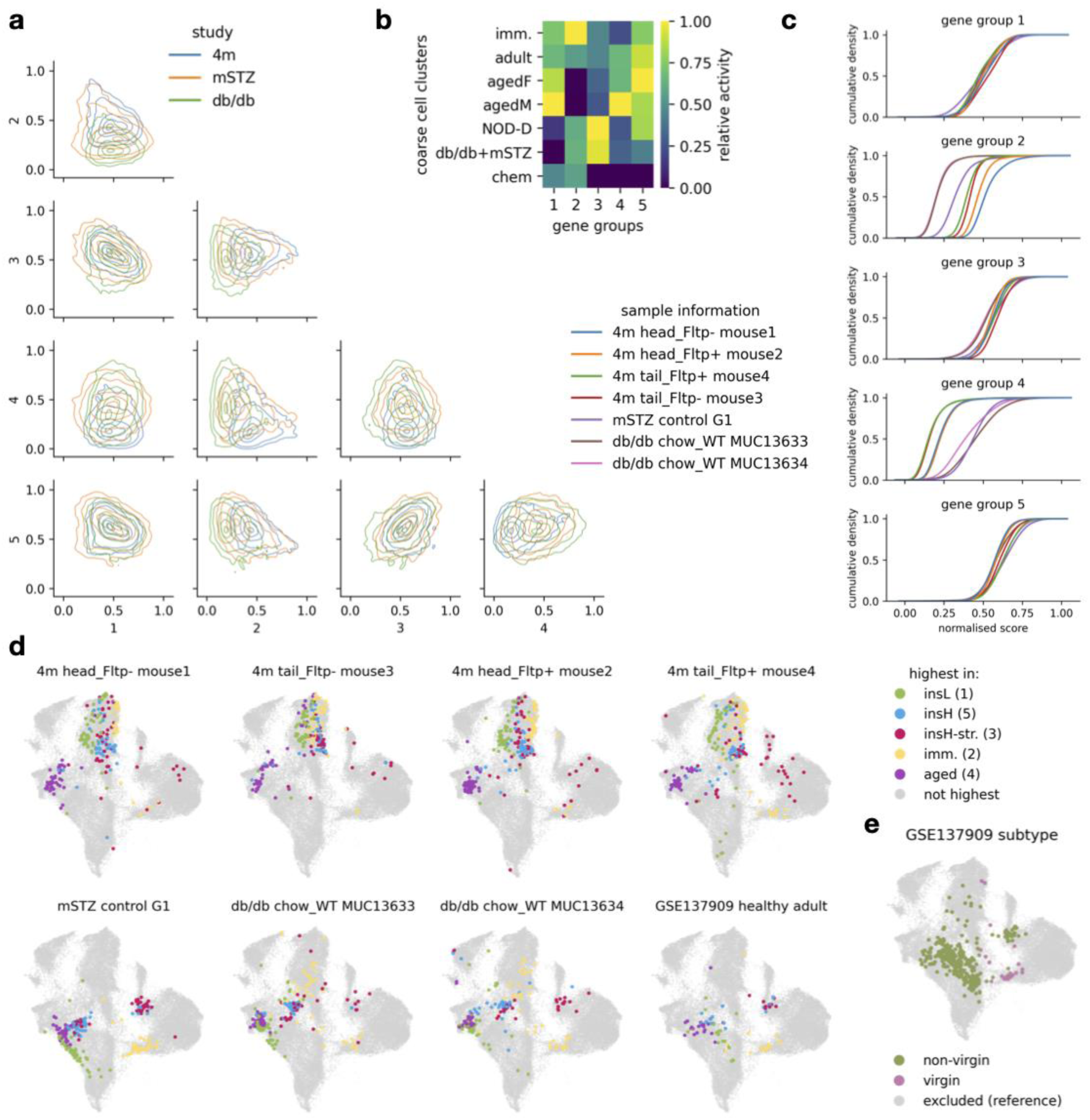
Healthy β-cells contain five distinct variable gene groups. (a) Pairwise comparison of the normalized activity of gene groups on β-cells from healthy samples shown as kernel-estimated density plots colored by study. Lines represent regions containing 5, 34, 67, 95, and 99% of cells. Axes represent activity of the compared gene groups. (b) Mean gene group activity within coarse β-cell clusters, normalized across clusters. (c) Per-sample distribution of normalized gene group activities in β-cells shown as cumulative density functions. (d) Localization of cells with the highest activity of the five gene groups on the atlas β-cell UMAP for individual health adult atlas samples (named as: dataset sample_metadata sample_name) and the healthy adult samples from the external dataset (Feng, GSE137909) mapped on top of the atlas. (e) Localization of virgin and non-virgin β-cells from Feng (GSE137909) dataset on the integrated atlas UMAP as annotated in the original publication.

**Supplementary Figure S12:**
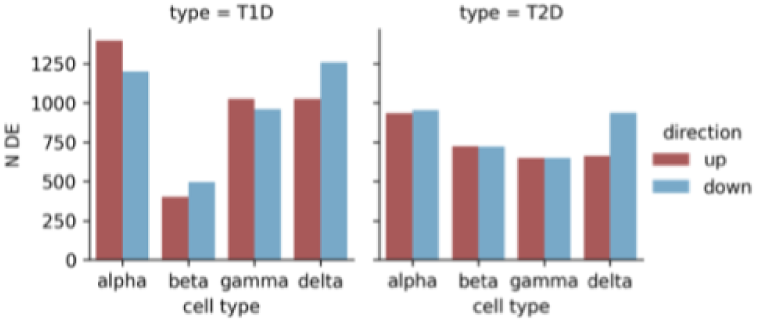
All endocrine cell types exhibit expression changes in T1D (NOD) and T2D (db/db+mSTZ) models. Number of DEGs across cell types and diabetes models.

**Supplementary Figure S13:**
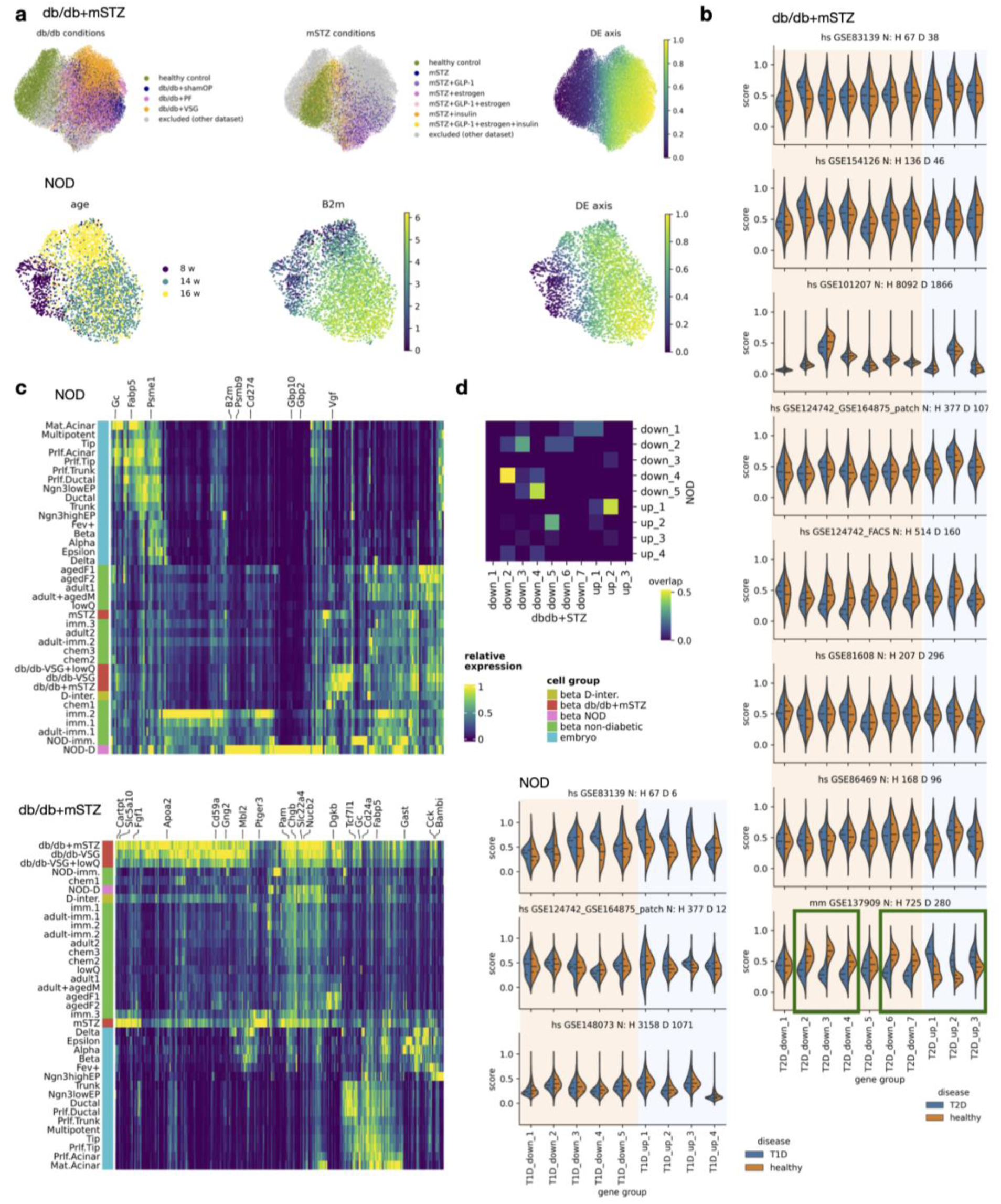
Diabetes-related molecular changes of β-cells show similarities and differences across dysfunctional states and translate to an external mouse dataset. (a) Design of DGE analysis showing original conditions in each used dataset (dataset 8-16wNOD for NOD DGE, datasets db/db and mSTZ for db/db+mSTZ DGE) and axis used for fitting the DGE model. For NOD we also show expression of a known T1D marker *B2m*. (b) Translation of diabetes model DEG groups (T1D -NOD, T2D -db/db+mSTZ) to external human and mouse datasets, indicated as normalized activity of gene groups in T1D or T2D (in mice STZ-treated) and healthy samples. Plot titles contain information on species (hs - human, mm - mouse), dataset, and number of cells in healthy (H) and diabetic (D) groups. Encircled are gene groups that translate to the external mouse dataset. (c) Expression of genes upregulated in diabetic NOD or db/db+mSTZ cells shown across fine β-cell states and embryonic cell types as annotated in the original study. Cell color annotations are based on healthy and developmental conditions. (d) Overlap between NOD and db/db+mSTZ DEG groups as ratio of the smaller group.

**Supplementary Figure S14:**
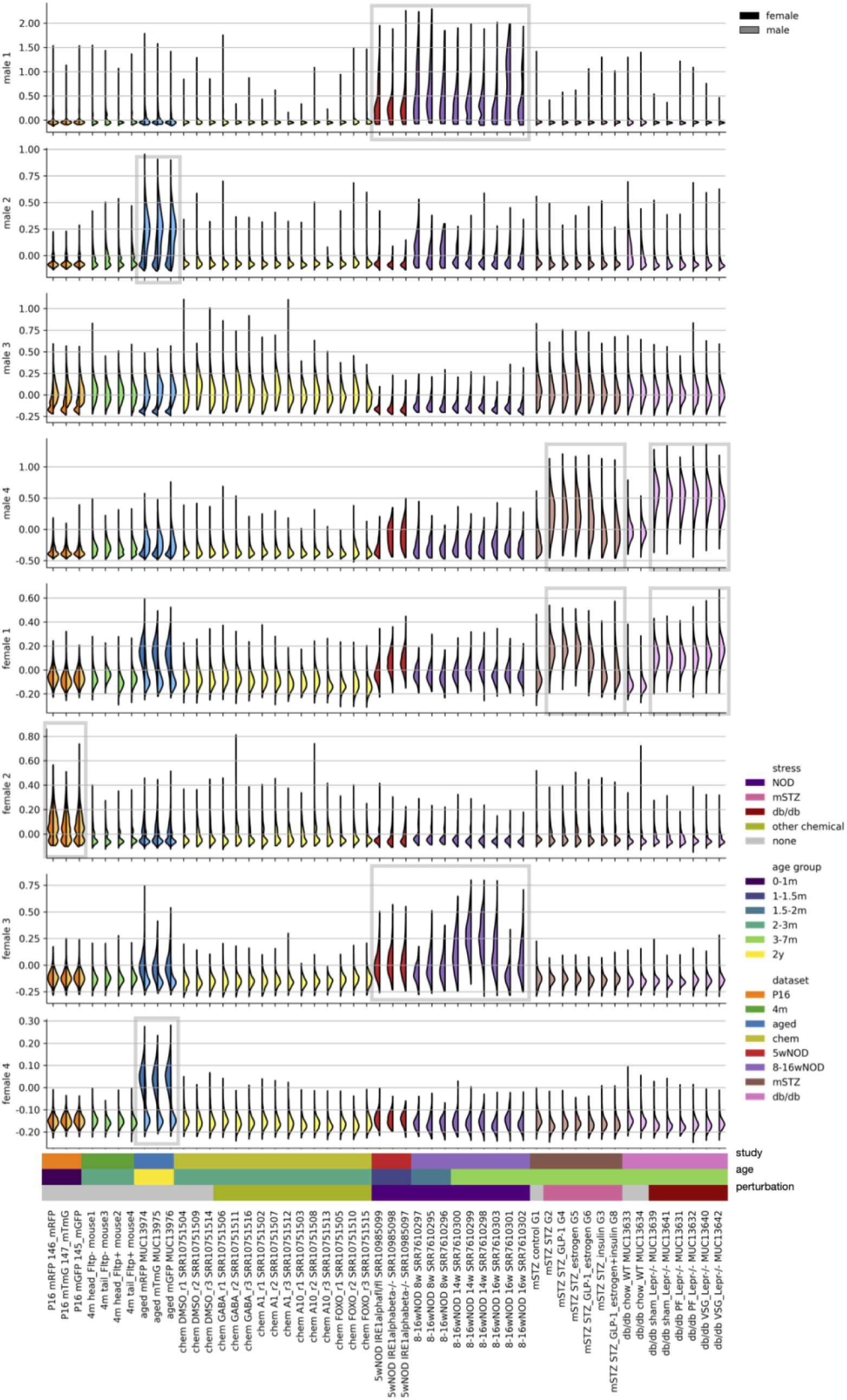
Genes differentially expressed between aged male and female β-cells cluster in distinct groups, some of which also show high activity in other conditions. Plot shows activity of gene groups across samples and sexes (indicated by color darkness). Sample names are given as study_sampleDescription_sampleIdentifier.

**Supplementary Figure S15:**
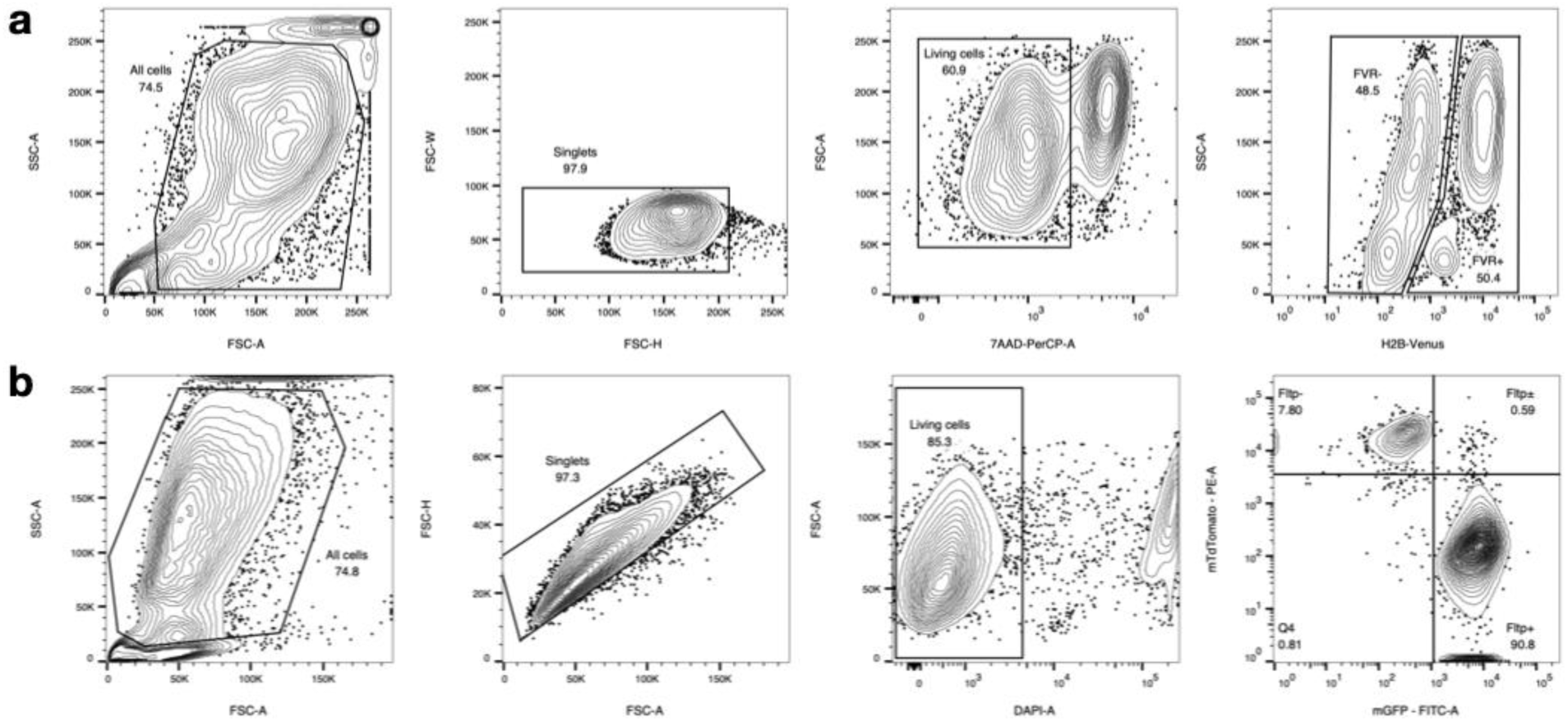
FACS analysis of samples used for newly generated scRNA-seq datasets. Sorting strategy for (a) Fltp^ZV/+^ and (b) Fltp iCre mTmG mouse strains. On plots are reported names and percentages of the used populations.

### Supplementary tables

**Supplementary Table S1:** Detailed description of individual samples from each dataset (given in separate sheets).

**Supplementary Table S2:** Number of cells per cell type (sheet name cell_type) and β-cell state (sheet names beta_subtype_coarse and beta_subtype_fine) within atlas samples. Samples in columns are named as “dataset sample_description sample_name”. Sheet cell_type also contains columns: total -number of all cells within cluster; low_quality - number of low quality cells within cluster, where blank/nan means that annotation of low quality cells was not performed within that cluster; doublet - whether that cluster was assigned to contain doublets/multiplets.

**Supplementary Table S3:** Endocrine cell type markers in postnatal and embryonic data. On each sheet is given the result of a single developmental stage and cell type combination.

**Supplementary Table S4:** Results of human healthy (reference level) to T1D and T2D β-cell comparison for each dataset (sheets T1D_DE and T2D_DE, concatenated across datasets), gene set enrichment of genes consistently upregulated in human T1D or T2D β-cells, and comparison of gene set activity between healthy and dysfunctional mouse diabetes model β-cell groups for selected human diabetes-associated gene sets (sheet name mouse_genesets_DE). Column names for the DGE tables: dataset - dataset in which DGE was performed; n_group - n cells from diabetic samples; n_ref - n healthy cells; ratio_group - ratio of cells from diabetic samples expressing the gene. Column names for the mouse comparison table: me_logFC - logFC on medians.

**Supplementary Table S5:** Results of coarse β-cell state marker identification. Column names: n_cl_de - number of clusters against which the marker was upregulated in the cluster of interest, min_lfc - minimal lFC across comparisons.

**Supplementary Table S6:** Genes present in GPs explaining variability across β-cells (sheet name GPs), their interpretation, gene set enrichment, and variance explained by selected diabetes-associated GPs in β-cells of healthy mouse and human samples (sheet name sample_explained_var). Column name explanation for the variance table: n_cells - n β-cells present in each sample.

**Supplementary Table S7:** β-cell heterogeneity markers (sheet beta_heterogeneity_markers) and function genes (sheet beta_function_genes) manually extracted from literature. Explanation of column names: gene_name - gene symbol of the associated organism, subtype - β-cell subtype with “NA” used when a specific subtype was not associated and “dedifferentiated” subtype also encompassing T2D models, organism - animal species for which marker was reported, function - molecular function associated with the gene, function_specific - subcategory of function, reference - DOI of papers from which gene information was extracted.

**Supplementary Table S8:** Genes present in gene groups consistently variable across healthy adult β-cells (sheet name GPs), their interpretation, and gene set enrichment.

**Supplementary Table S9:** Results of DGE analysis between healthy and different diabetes-model groups across endocrine cell types; given in individual sheets, named as celltype_diabestype_DE, with T1D representing NOD model and T2D representing db/db+mSTZ models. Additionally, gene set enrichment of genes jointly up or downregulated across diabetes models in α-, γ-, and δ-cells (sheet names starting with sharedADG_DE and ending with DGE direction).

**Supplementary Table S10:** Results of DGE analysis and DEG clustering in 8-16wNOD (sheet T1D_DE) or db/db+mSTZ (Sheet T2D_DE) healthy (reference level) and dysfunctional β-cell comparison, gene set enrichment of individual DEG groups, DEG group interpretation, and gene set enrichment of DEGs in the same direction in db/db+mSTZ and 8-16wNOD DGE tests (sheet names starting with sharedT1DT2D_DE and ending with DGE direction).

**Supplementary Table S11:** Results of DGE analysis between male (reference level) and female β-cells across datasets (reported in sheets). The aged dataset also contains DEG groups named as down for male and up for female upregulated groups and their interpretation.

**Supplementary Table S12:** Detailed description of individual samples from each of the external validation datasets.

**Supplementary Table S13:** List of antibodies used in laboratory validation of diabetes markers.

Explanation of column names used in multiple tables: hc - gene group, if empty the gene was not assigned to any gene group; rel_beta_expr/rel_ct_expr - relative expression in β-cells/relevant cell type compared to other cell types; N_PMID - number of papers associated with a gene; N_PMID_pancreas_notCancerNonendo - as N_PMID, but filtering papers based on terms that indicate the paper is focused on endocrine pancreas compartment; mean_expr_in_expr_cells - mean expression in β- cells that have above zero expression.

Explanation of enrichment sheets: Sheet names containing the term “enrichment” represent gene set enrichment for individual gene groups. When enrichment analysis did not produce any significant results for a specific gene group the corresponding enrichment sheet is empty. Columns: signature/query_size - N genes in gene group, geneset - N genes in gene set, overlap - N overlapping genes, background - N genes in the background, hits - overlapping genes, recall - ratio overlap/geneset.

Explanation of gene group interpretation sheets: Sheet names containing the term “interpretation” contain interpretation for gene groups within the excel file. Columns: group - group name, enrichment - manually curated summary of gene set enrichment, example_genes - example genes contained within the group, cell_cluster_expression - summary of expression pattern across β-cell states. Empty table cells indicate lack of interpretation for given gene group characteristics.

