## Supplementary material for "Delineating mouse β-cell identity during lifetime and in diabetes with a single cell atlas": High-resolution figures

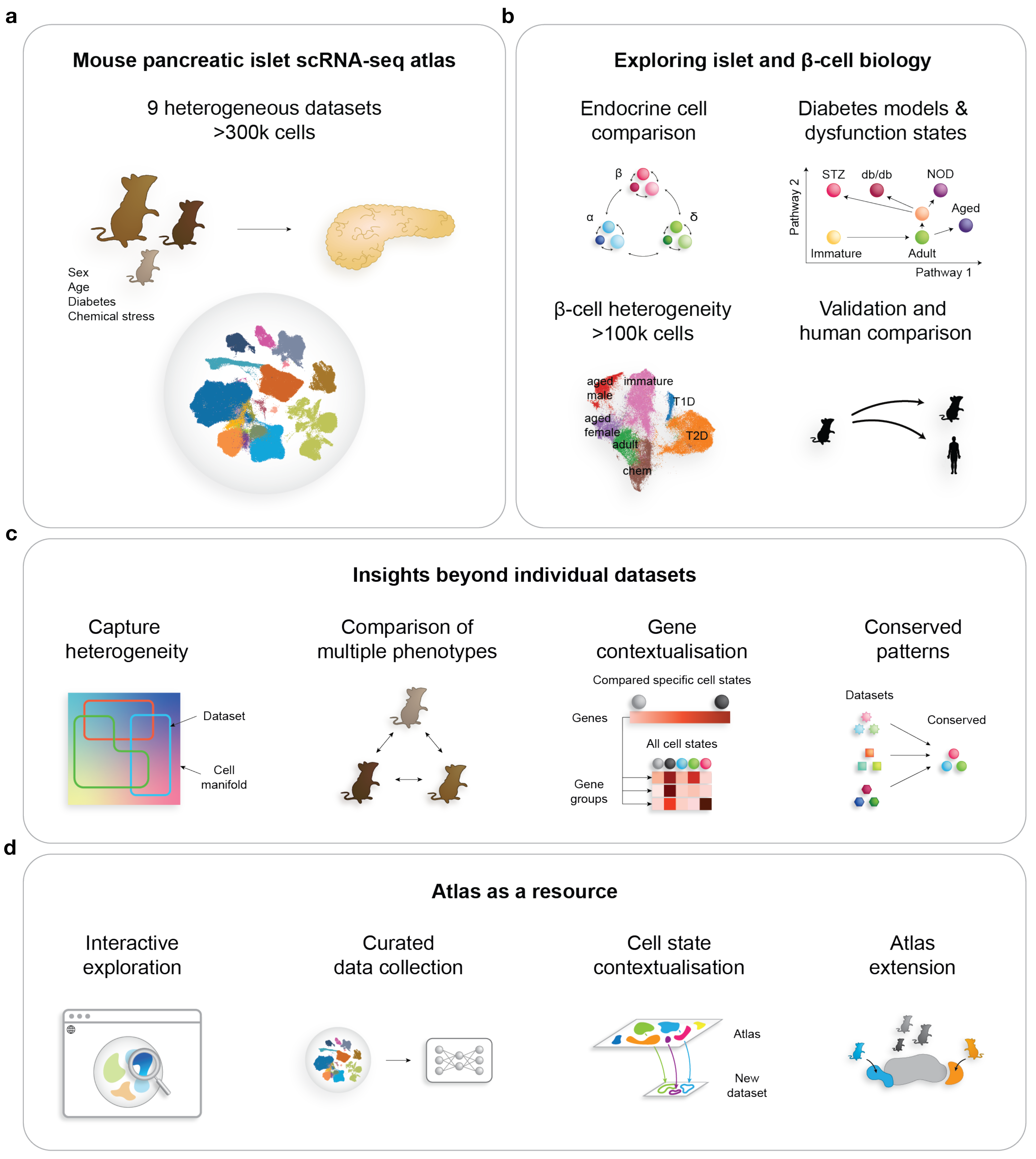
a
Mouse pancreatic islet scRNA-seq atlas
9 heterogeneous datasets
>300k cells
Sex
Age
Diabetes
Chemical stress
b
Exploring islet and β-cell biology
Endocrine cell comparison
β
α
δ
Diabetes models & dysfunction states
Pathway 2
STZ
db/db
NOD
Aged
Immature
Adult
Pathway 1
β-cell heterogeneity
>100k cells
aged male
immature
T1D
aged female
adult
T2D
chem
Validation and human comparison
c
Insights beyond individual datasets
Capture heterogeneity
Dataset
Cell manifold
Comparison of multiple phenotypes
Gene contextualisation
Compared specific cell states
Genes
All cell states
Gene groups
Conserved patterns
Datasets
Conserved
d
Atlas as a resource
Interactive exploration
Curated data collection
Cell state contextualisation
Atlas
New dataset
Atlas extension

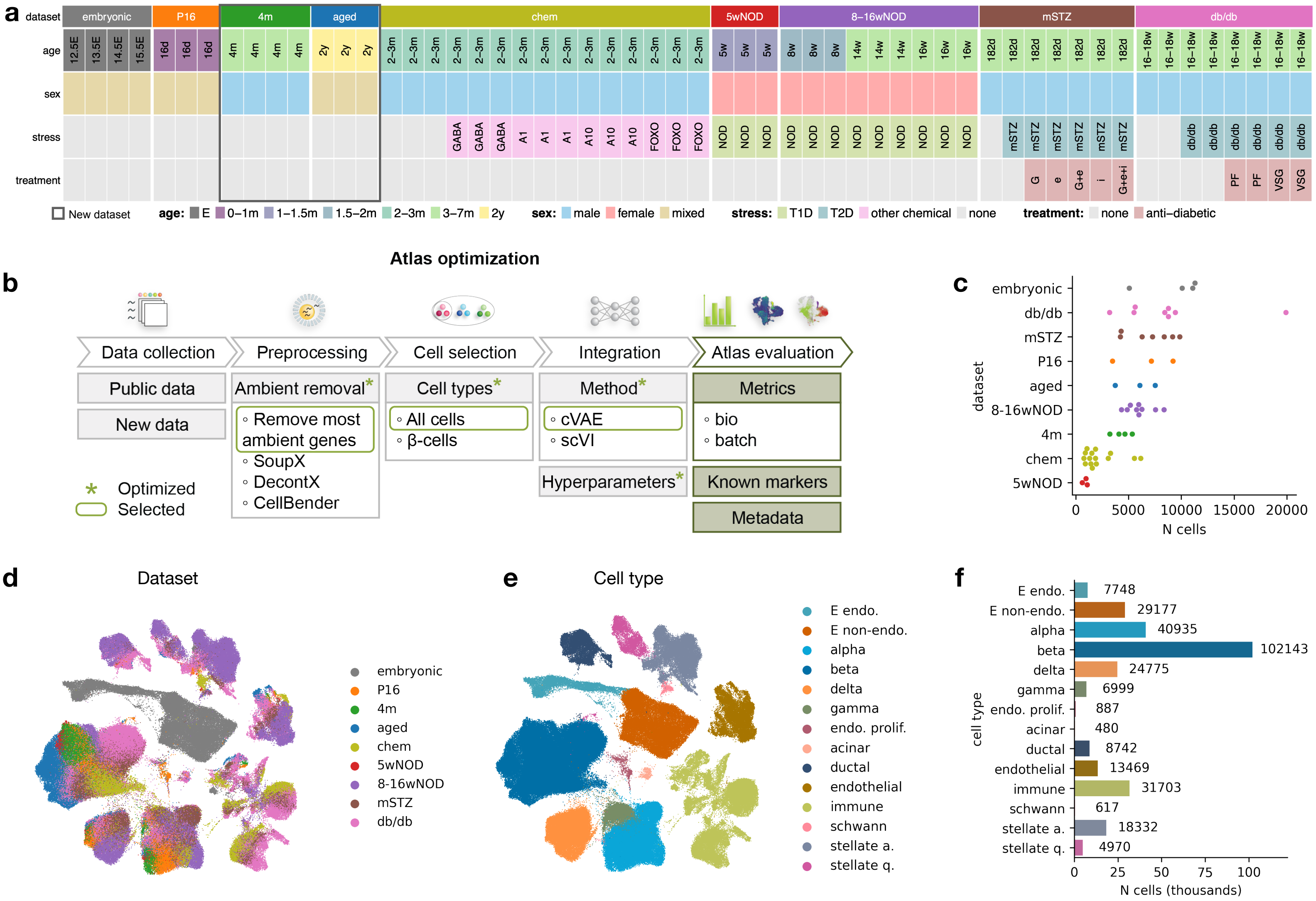
a
dataset
embryonic
P16
4m
aged
chem
5wNOD
8–16wNOD
mSTZ
db/db
age
12.5E
13.5E
14.5E
15.5E
16d
4m
2y
2–3m
5w
8w
14w
16w
182d
16–18w
sex
stress
GABA
A1
A10
FOXO
NOD
mSTZ
db/db
treatment
G
e
G+e
i
G+e+i
PF
VSG
New dataset
age: E 0–1m 1–1.5m 1.5–2m 2–3m 3–7m 2y
sex: male female mixed
stress: T1D T2D other chemical none
treatment: none anti–diabetic
Atlas optimization
b
Data collection
Preprocessing
Cell selection
Integration
Atlas evaluation
Public data
New data
Ambient removal*
◦ Remove most ambient genes
◦ SoupX
◦ DecontX
◦ CellBender
Cell types*
◦ All cells
◦ β-cells
Method*
◦ cVAE
◦ scVI
Hyperparameters*
Metrics
◦ bio
◦ batch
Known markers
Metadata
* Optimized
Selected
c
embryonic
db/db
mSTZ
P16
aged
8-16wNOD
4m
chem
5wNOD
dataset
0
5000
10000
15000
20000
N cells
d
Dataset
embryonic
P16
4m
aged
chem
5wNOD
8-16wNOD
mSTZ
db/db
e
Cell type
E endo.
E non-endo.
alpha
beta
delta
gamma
endo. prolif.
acinar
ductal
endothelial
immune
schwann
stellate a.
stellate q.
f
E endo. 7748
E non-endo. 29177
alpha 40935
beta 102143
delta 24775
gamma 6999
endo. prolif. 887
acinar 480
ductal 8742
endothelial 13469
immune 31703
schwann 617
stellate a. 18332
stellate q. 4970
cell type
0
25
50
75
100
N cells (thousands)

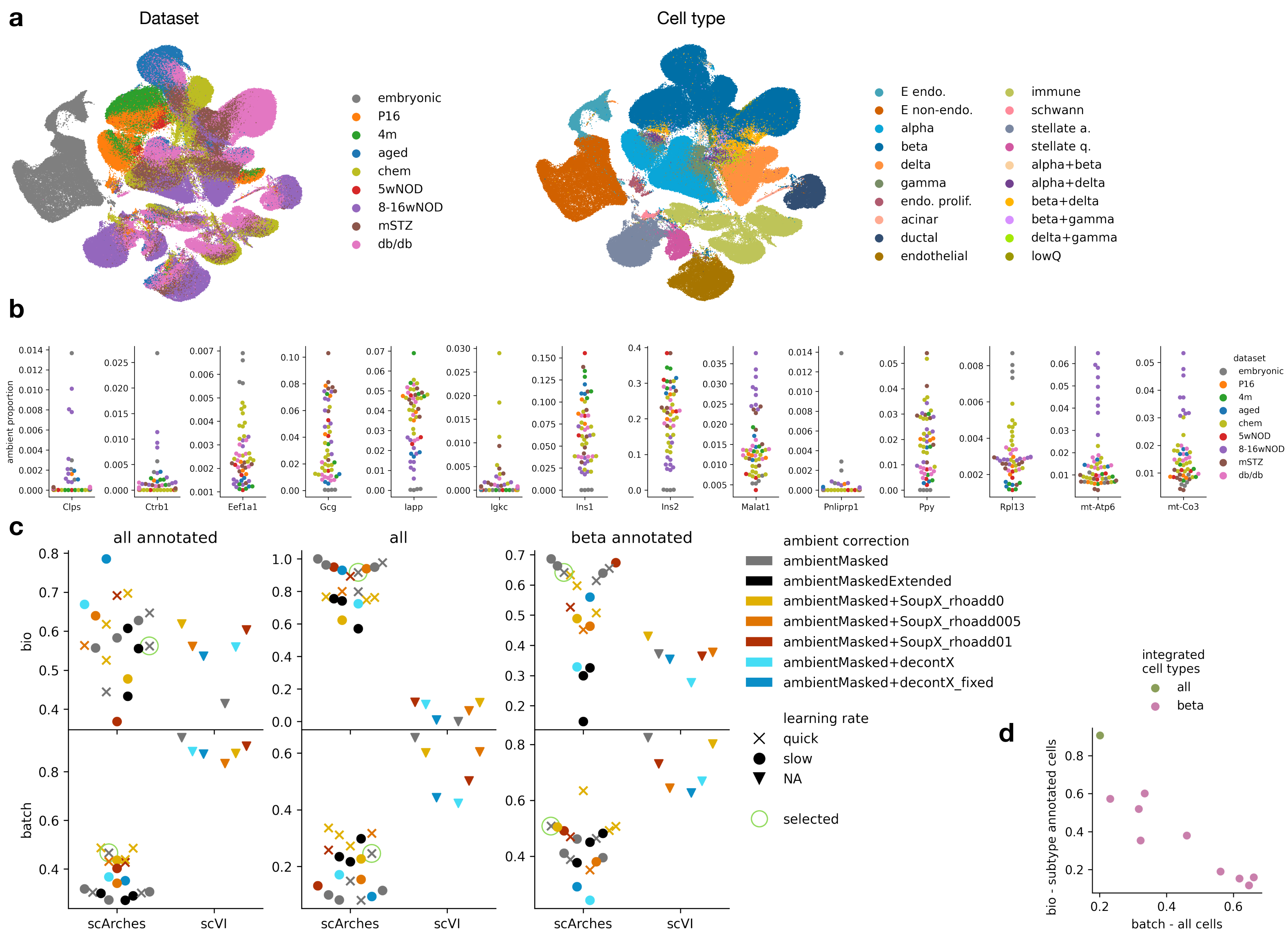
a
Dataset
embryonic
P16
4m
aged
chem
5wNOD
8-16wNOD
mSTZ
db/db
Cell type
E endo.
E non-endo.
alpha
beta
delta
gamma
endo. prolif.
acinar
ductal
endothelial
immune
schwann
stellate a.
stellate q.
alpha+beta
alpha+delta
beta+delta
beta+gamma
delta+gamma
lowQ
b
ambient proportion
Clps
Ctrb1
Eef1a1
Gcg
Iapp
Igkc
Ins1
Ins2
Malat1
Pnliprp1
Ppy
Rpl13
mt-Atp6
mt-Co3
dataset
embryonic
P16
4m
aged
chem
5wNOD
8-16wNOD
mSTZ
db/db
c
all annotated
all
beta annotated
bio
batch
scArches
scVI
ambient correction
ambientMasked
ambientMaskedExtended
ambientMasked+SoupX_rhoadd0
ambientMasked+SoupX_rhoadd005
ambientMasked+SoupX_rhoadd01
ambientMasked+decontX
ambientMasked+decontX_fixed
learning rate
quick
slow
NA
selected
integrated
cell types
all
beta
d
bio - subtype annotated cells
batch - all cells

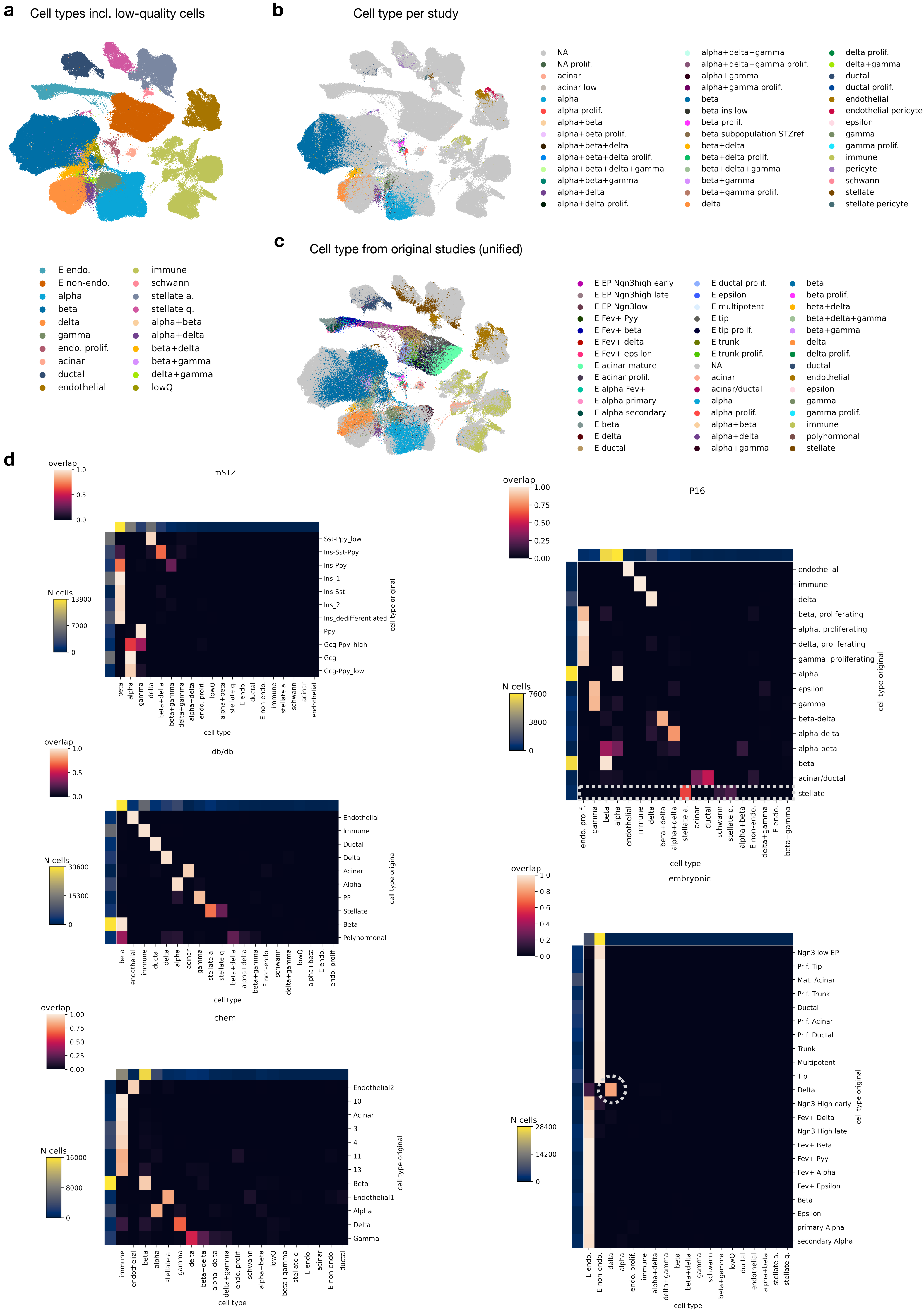

a
Cell types incl. low-quality cells
E endo.
E non-endo.
alpha
beta
delta
gamma
endo. prolif.
acinar
ductal
endothelial
immune
schwann
stellate a.
stellate q.
alpha+beta
alpha+delta
beta+delta
beta+gamma
delta+gamma
lowQ
b
Cell type per study
NA
NA prolif.
acinar
acinar low
alpha
alpha prolif.
alpha+beta
alpha+beta prolif.
alpha+beta+delta
alpha+beta+delta prolif.
alpha+beta+delta+gamma
alpha+beta+gamma
alpha+delta
alpha+delta prolif.
alpha+delta+gamma
alpha+delta+gamma prolif.
alpha+gamma
alpha+gamma prolif.
beta
beta ins low
beta prolif.
beta subpopulation STZref
beta+delta
beta+delta prolif.
beta+delta+gamma
beta+gamma
beta+gamma prolif.
delta
delta prolif.
delta+gamma
ductal
ductal prolif.
endothelial
endothelial pericyte
epsilon
gamma
gamma prolif.
immune
pericyte
schwann
stellate
stellate pericyte
c
Cell type from original studies (unified)
E EP Ngn3high early
E EP Ngn3high late
E EP Ngn3low
E Fev+ Pyy
E Fev+ beta
E Fev+ delta
E Fev+ epsilon
E acinar mature
E acinar prolif.
E alpha Fev+
E alpha primary
E alpha secondary
E beta
E delta
E ductal
E ductal prolif.
E epsilon
E multipotent
E tip
E tip prolif.
E trunk
E trunk prolif.
NA
acinar
acinar/ductal
alpha
alpha prolif.
alpha+beta
alpha+delta
alpha+gamma
beta
beta prolif.
beta+delta
beta+delta+gamma
beta+gamma
delta
delta prolif.
ductal
endothelial
epsilon
gamma
gamma prolif.
immune
polyhormonal
stellate
d
mSTZ
overlap
1.0
0.5
0.0
N cells
13900
7000
0
Sst-Ppy_low
Ins-Sst-Ppy
Ins-Ppy
Ins_1
Ins-Sst
Ins_2
Ins_dedifferentiated
Ppy
Gcg-Ppy_high
Gcg
Gcg-Ppy_low
cell type original
beta
alpha
gamma
delta
beta+delta
beta+gamma
delta+gamma
alpha+delta
endo. prolif.
lowQ
alpha+beta
stellate q.
E endo.
ductal
E non-endo.
immune
stellate a.
schwann
acinar
endothelial
cell type
db/db
overlap
1.0
0.5
0.0
N cells
30600
15300
0
Endothelial
Immune
Ductal
Delta
Acinar
Alpha
PP
Stellate
Beta
Polyhormonal
cell type original
beta
endothelial
immune
ductal
delta
alpha
acinar
gamma
stellate a.
stellate q.
beta+delta
alpha+delta
beta+gamma
E non-endo.
schwann
delta+gamma
lowQ
alpha+beta
E endo.
endo. prolif.
cell type
chem
overlap
1.00
0.75
0.50
0.25
0.00
N cells
16000
8000
0
Endothelial2
10
Acinar
3
4
11
13
Beta
Endothelial1
Alpha
Delta
Gamma
cell type original
immune
endothelial
beta
alpha
stellate a.
gamma
delta
beta+delta
alpha+delta
delta+gamma
endo. prolif.
schwann
alpha+beta
lowQ
beta+gamma
stellate q.
E endo.
acinar
E non-endo.
ductal
cell type
P16
overlap
1.00
0.75
0.50
0.25
0.00
N cells
7600
3800
0
endothelial
immune
delta
beta, proliferating
alpha, proliferating
delta, proliferating
gamma, proliferating
alpha
epsilon
gamma
beta-delta
alpha-delta
alpha-beta
beta
acinar/ductal
stellate
cell type original
endo. prolif.
gamma
beta
alpha
endothelial
immune
delta
beta+delta
alpha+delta
stellate a.
acinar
ductal
schwann
stellate q.
alpha+beta
E non-endo.
delta+gamma
E endo.
beta+gamma
cell type
embryonic
overlap
1.0
0.8
0.6
0.4
0.2
0.0
N cells
28400
14200
0
Ngn3 low EP
Prlf. Tip
Mat. Acinar
Prlf. Trunk
Ductal
Prlf. Acinar
Prlf. Ductal
Trunk
Multipotent
Tip
Delta
Ngn3 High early
Fev+ Delta
Ngn3 High late
Fev+ Beta
Fev+ Pyy
Fev+ Alpha
Fev+ Epsilon
Beta
Epsilon
primary Alpha
secondary Alpha
cell type original
E endo.
E non-endo.
delta
alpha
endo. prolif.
immune
alpha+delta
delta+gamma
beta
beta+delta
gamma
schwann
beta+gamma
lowQ
ductal
endothelial
alpha+beta
stellate a.
stellate q.
cell type

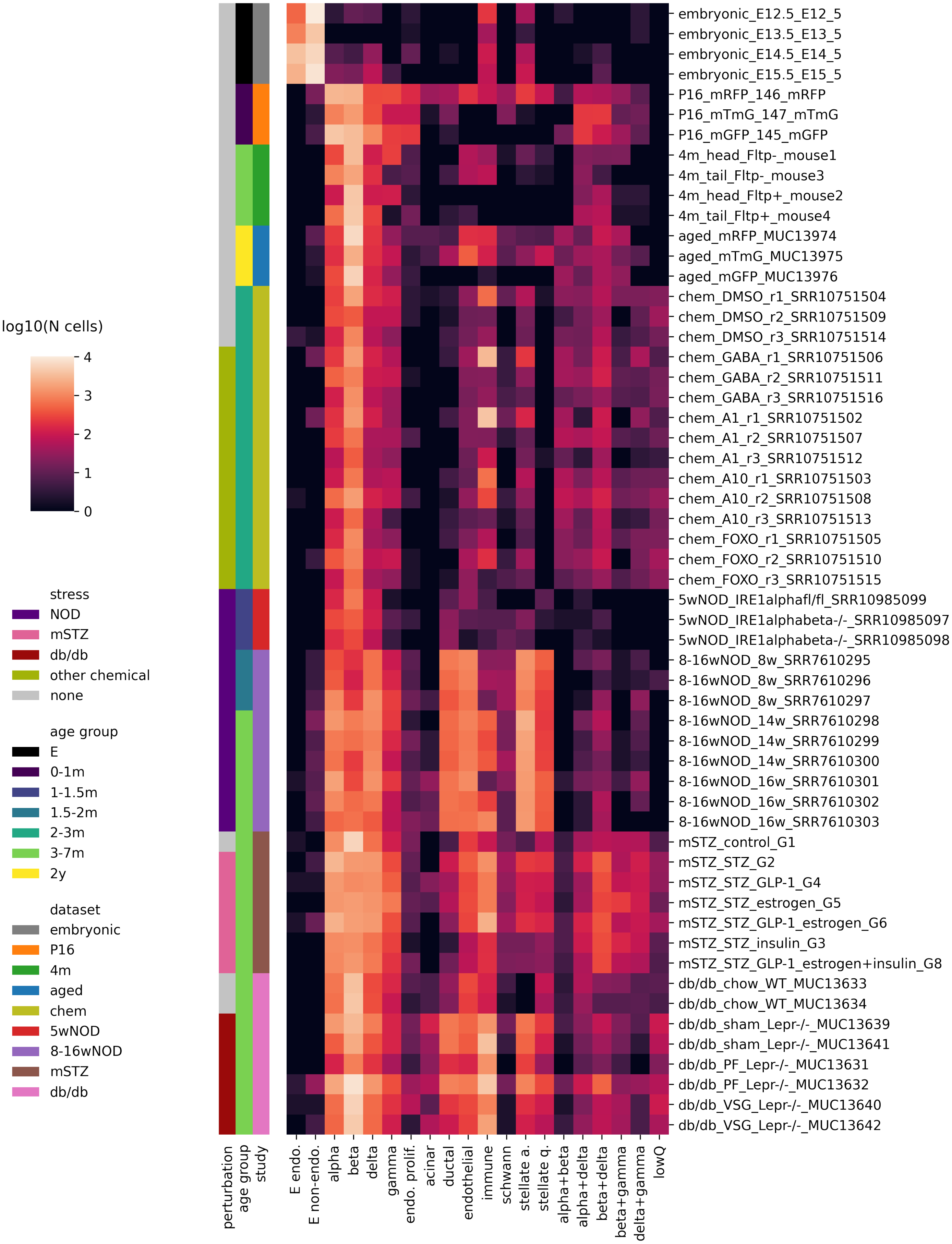
log10(N cells)
4
3
2
1
0
stress
NOD
mSTZ
db/db
other chemical
none
age group
E
0-1m
1-1.5m
1.5-2m
2-3m
3-7m
2y
dataset
embryonic
P16
4m
aged
chem
5wNOD
8-16wNOD
mSTZ
db/db
embryonic_E12.5_E12_5
embryonic_E13.5_E13_5
embryonic_E14.5_E14_5
embryonic_E15.5_E15_5
P16_mRFP_146_mRFP
P16_mTmG_147_mTmG
P16_mGFP_145_mGFP
4m_head_Fltp-_mouse1
4m_tail_Fltp-_mouse3
4m_head_Fltp+_mouse2
4m_tail_Fltp+_mouse4
aged_mRFP_MUC13974
aged_mTmG_MUC13975
aged_mGFP_MUC13976
chem_DMSO_r1_SRR10751504
chem_DMSO_r2_SRR10751509
chem_DMSO_r3_SRR10751514
chem_GABA_r1_SRR10751506
chem_GABA_r2_SRR10751511
chem_GABA_r3_SRR10751516
chem_A1_r1_SRR10751502
chem_A1_r2_SRR10751507
chem_A1_r3_SRR10751512
chem_A10_r1_SRR10751503
chem_A10_r2_SRR10751508
chem_A10_r3_SRR10751513
chem_FOXO_r1_SRR10751505
chem_FOXO_r2_SRR10751510
chem_FOXO_r3_SRR10751515
5wNOD_IRE1alphafl/fl_SRR10985099
5wNOD_IRE1alphabeta-/-_SRR10985097
5wNOD_IRE1alphabeta-/-_SRR10985098
8-16wNOD_8w_SRR7610295
8-16wNOD_8w_SRR7610296
8-16wNOD_8w_SRR7610297
8-16wNOD_14w_SRR7610298
8-16wNOD_14w_SRR7610299
8-16wNOD_14w_SRR7610300
8-16wNOD_16w_SRR7610301
8-16wNOD_16w_SRR7610302
8-16wNOD_16w_SRR7610303
mSTZ_control_G1
mSTZ_STZ_G2
mSTZ_STZ_GLP-1_G4
mSTZ_STZ_estrogen_G5
mSTZ_STZ_GLP-1_estrogen_G6
mSTZ_STZ_insulin_G3
mSTZ_STZ_GLP-1_estrogen+insulin_G8
db/db_chow_WT_MUC13633
db/db_chow_WT_MUC13634
db/db_sham_Lepr-/-_MUC13639
db/db_sham_Lepr-/-_MUC13641
db/db_PF_Lepr-/-_MUC13631
db/db_PF_Lepr-/-_MUC13632
db/db_VSG_Lepr-/-_MUC13640
db/db_VSG_Lepr-/-_MUC13642
perturbation
age group
study
E endo.
E non-endo.
alpha
beta
delta
gamma
endo. prolif.
acinar
ductal
endothelial
immune
schwann
stellate a.
stellate q.
alpha+beta
alpha+delta
beta+delta
beta+gamma
delta+gamma
lowQ

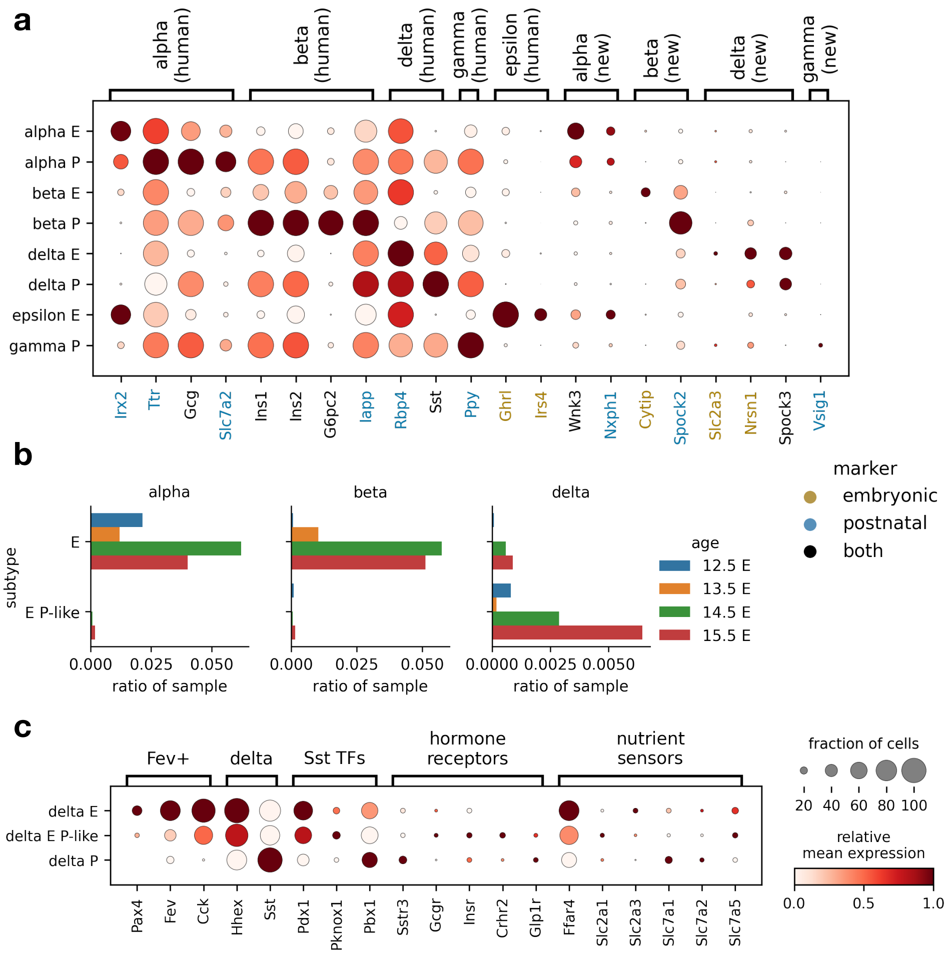

a
alpha (human)
beta (human)
delta (human)
gamma (human)
epsilon (human)
alpha (new)
beta (new)
delta (new)
gamma (new)
alpha E
alpha P
beta E
beta P
delta E
delta P
epsilon E
gamma P
Irx2
Ttr
Gcg
Slc7a2
Ins1
Ins2
G6pc2
Iapp
Rbp4
Sst
Ppy
Ghrl
Irs4
Wnk3
Nxph1
Cytip
Spock2
Slc2a3
Nrsn1
Spock3
Vsig1
b
alpha
beta
delta
subtype
E
E P-like
0.000
0.025
0.050
0.0000
0.0025
0.0050
ratio of sample
age
12.5 E
13.5 E
14.5 E
15.5 E
marker
embryonic
postnatal
both
c
Fev+
delta
Sst TFs
hormone receptors
nutrient sensors
delta E
delta E P-like
delta P
Pax4
Fev
Cck
Hhex
Sst
Pdx1
Pknox1
Pbx1
Sstr3
Gcgr
Insr
Crhr2
Glp1r
Ffar4
Slc2a1
Slc2a3
Slc7a1
Slc7a2
Slc7a5
fraction of cells
20
40
60
80
100
relative mean expression
0.0
0.5
1.0

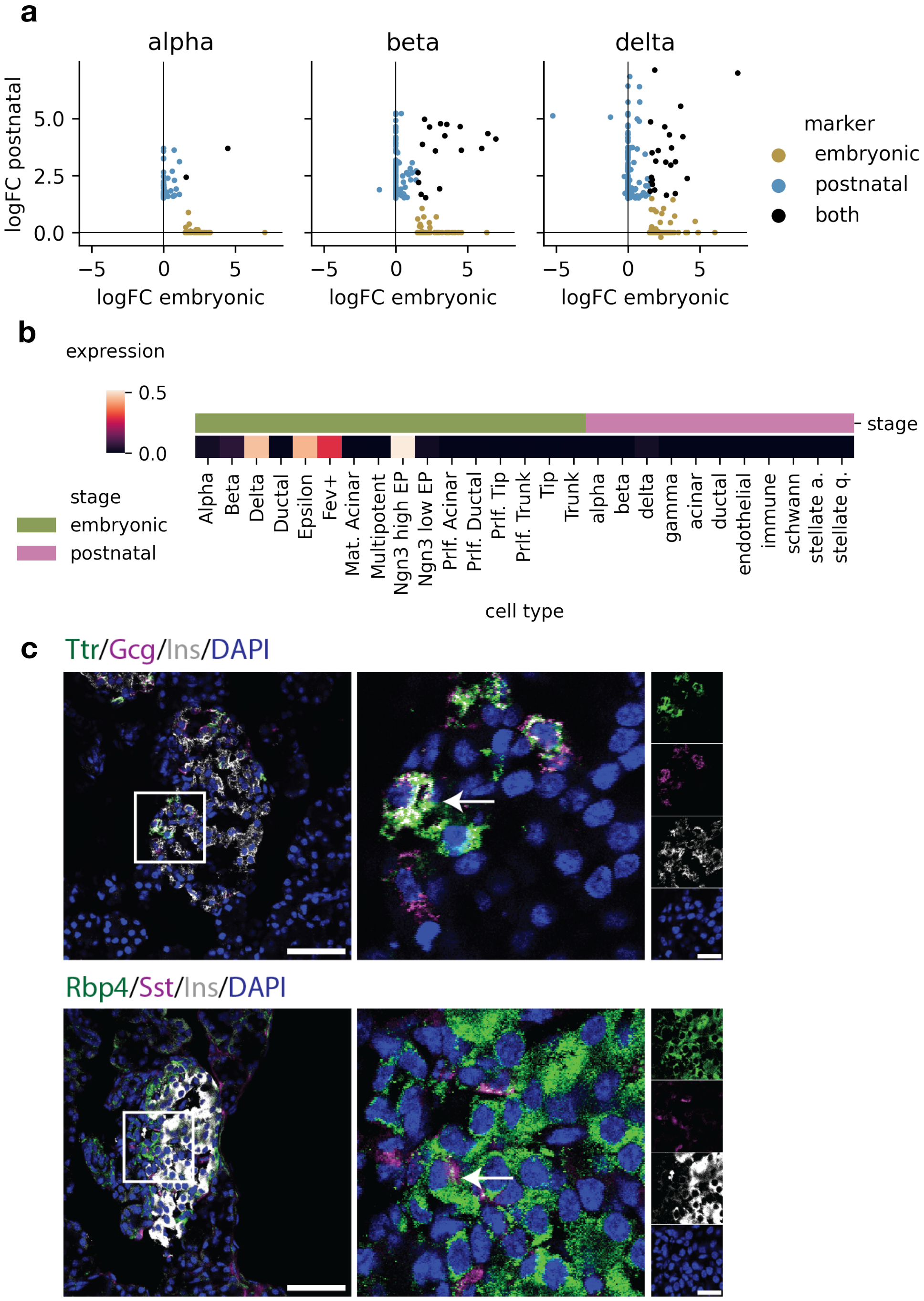
a
alpha
beta
delta
logFC postnatal
logFC embryonic
marker
embryonic
postnatal
both
b
expression
stage
embryonic
postnatal
Alpha
Beta
Delta
Ductal
Epsilon
Fev+
Mat. Acinar
Multipotent
Ngn3 high EP
Ngn3 low EP
Prlf. Acinar
Prlf. Ductal
Prlf. Tip
Prlf. Trunk
Tip
Trunk
alpha
beta
delta
gamma
acinar
ductal
endothelial
immune
schwann
stellate a.
stellate q.
cell type
c
Ttr/Gcg/Ins/DAPI
Rbp4/Sst/Ins/DAPI

**a**

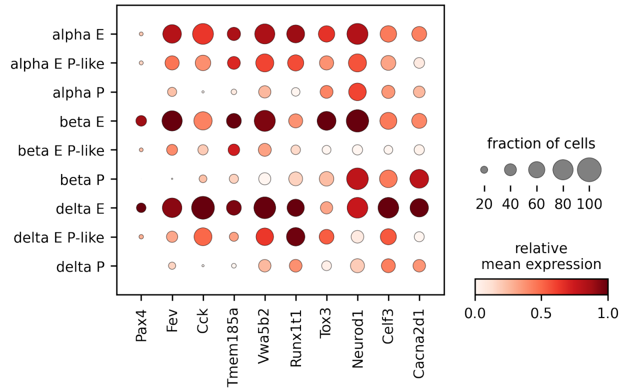


**b**

| cell type | N cells |
|---|---|
| alpha E | 1378 |
| alpha E P-like | 32 |
| alpha P | 40903 |
| beta E | 1220 |
| beta E P-like | 34 |
| beta P | 102109 |
| delta E | 17 |
| delta E P-like | 112 |
| delta P | 24663 |

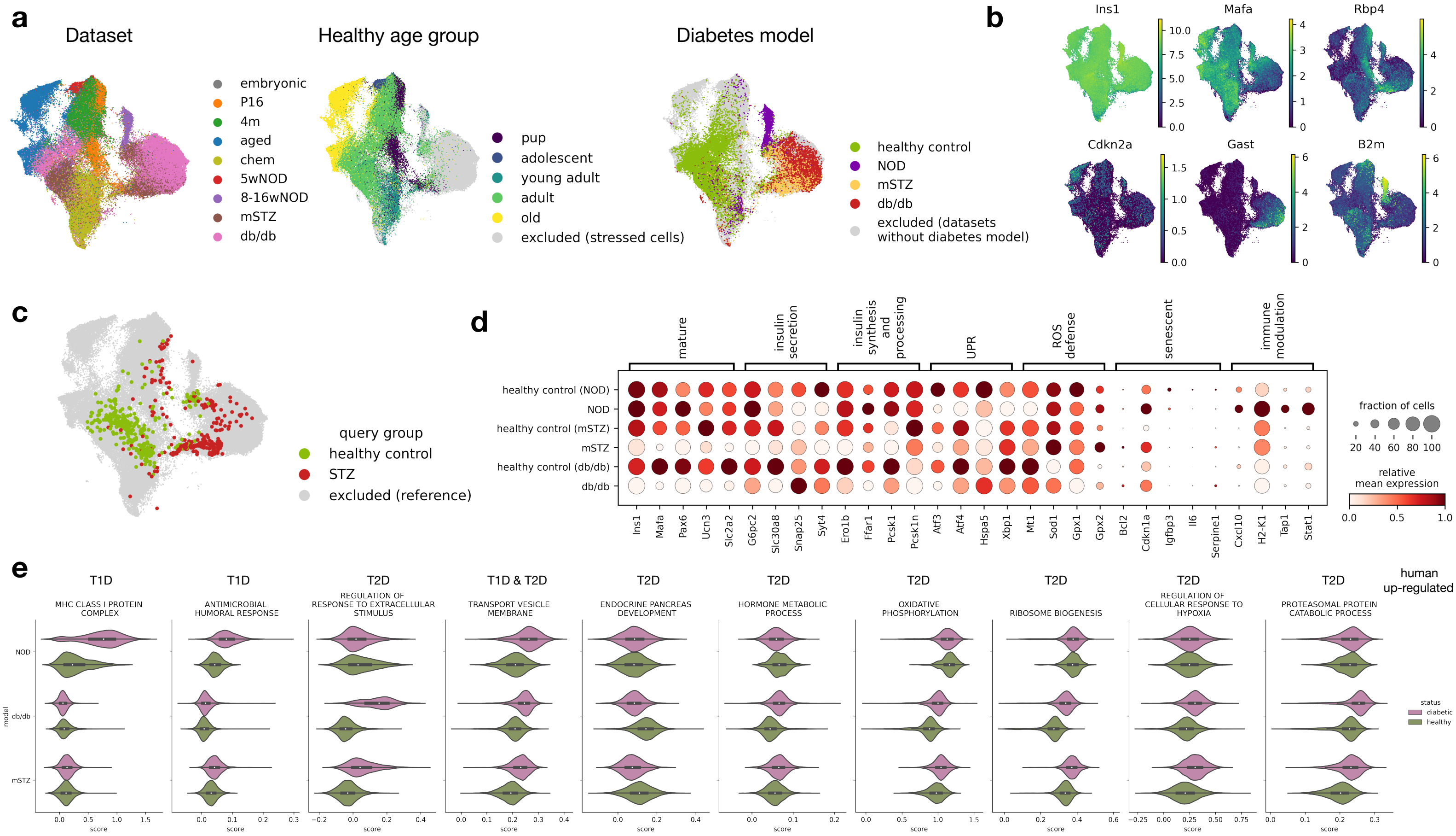
a
Dataset
embryonic
P16
4m
aged
chem
5wNOD
8-16wNOD
mSTZ
db/db
Healthy age group
pup
adolescent
young adult
adult
old
excluded (stressed cells)
Diabetes model
healthy control
NOD
mSTZ
db/db
excluded (datasets without diabetes model)
b
Ins1
Mafa
Rbp4
Cdkn2a
Gast
B2m
c
query group
healthy control
STZ
excluded (reference)
d
mature
insulin secretion
insulin synthesis and processing
UPR
ROS defense
senescent
immune modulation
healthy control (NOD)
NOD
healthy control (mSTZ)
mSTZ
healthy control (db/db)
db/db
Ins1
Mafa
Pax6
Ucn3
Slc2a2
G6pc2
Slc30a8
Snap25
Syt4
Ero1b
Ffar1
Pcsk1
Pcsk1n
Atf3
Atf4
Hspa5
Xbp1
Mt1
Sod1
Gpx1
Gpx2
Bcl2
Cdkn1a
Igfbp3
Il6
Serpine1
Cxcl10
H2-K1
Tap1
Stat1
fraction of cells
20 40 60 80 100
relative mean expression
0.0 0.5 1.0
e
T1D
MHC CLASS I PROTEIN COMPLEX
T1D
ANTIMICROBIAL HUMORAL RESPONSE
T2D
REGULATION OF RESPONSE TO EXTRACELLULAR STIMULUS
T1D & T2D
TRANSPORT VESICLE MEMBRANE
T2D
ENDOCRINE PANCREAS DEVELOPMENT
T2D
HORMONE METABOLIC PROCESS
T2D
OXIDATIVE PHOSPHORYLATION
T2D
RIBOSOME BIOGENESIS
T2D
REGULATION OF CELLULAR RESPONSE TO HYPOXIA
T2D
PROTEASOMAL PROTEIN CATABOLIC PROCESS
human up-regulated
model
NOD
db/db
mSTZ
score
status
diabetic
healthy

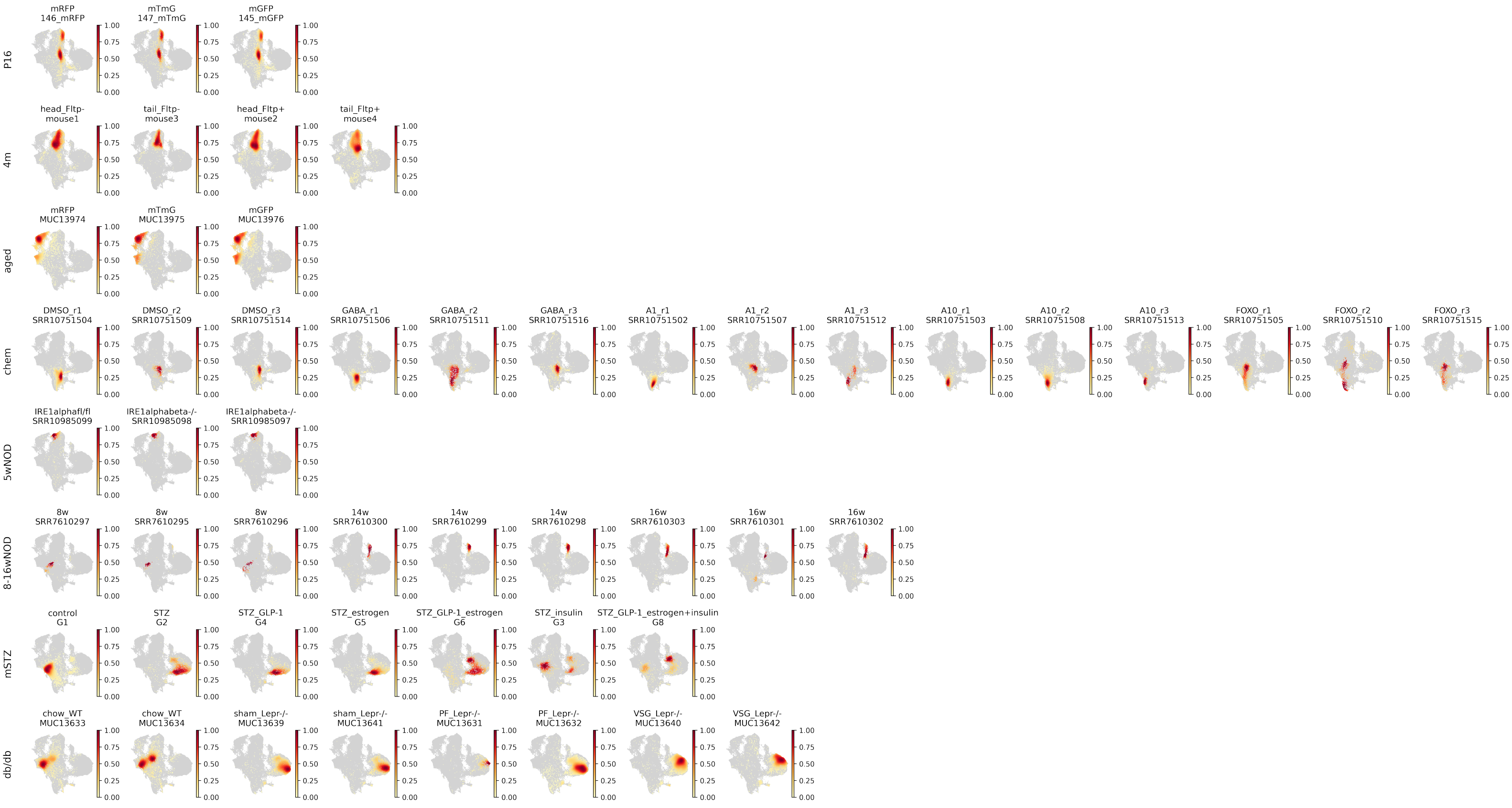
P16
mRFP 146_mRFP
mTmG 147_mTmG
mGFP 145_mGFP
4m
head_Fltp- mouse1
tail_Fltp- mouse3
head_Fltp+ mouse2
tail_Fltp+ mouse4
aged
mRFP MUC13974
mTmG MUC13975
mGFP MUC13976
chem
DMSO_r1 SRR10751504
DMSO_r2 SRR10751509
DMSO_r3 SRR10751514
GABA_r1 SRR10751506
GABA_r2 SRR10751511
GABA_r3 SRR10751516
A1_r1 SRR10751502
A1_r2 SRR10751507
A1_r3 SRR10751512
A10_r1 SRR10751503
A10_r2 SRR10751508
A10_r3 SRR10751513
FOXO_r1 SRR10751505
FOXO_r2 SRR10751510
FOXO_r3 SRR10751515
5wNOD
IRE1alphafl/fl SRR10985099
IRE1alphabeta-/- SRR10985098
IRE1alphabeta-/- SRR10985097
8-16wNOD
8w SRR7610297
8w SRR7610295
8w SRR7610296
14w SRR7610300
14w SRR7610299
14w SRR7610298
16w SRR7610303
16w SRR7610301
16w SRR7610302
mSTZ
control G1
STZ G2
STZ_GLP-1 G4
STZ_estrogen G5
STZ_GLP-1_estrogen G6
STZ_insulin G3
STZ_GLP-1_estrogen+insulin G8
db/db
chow_WT MUC13633
chow_WT MUC13634
sham_Lepr-/- MUC13639
sham_Lepr-/- MUC13641
PF_Lepr-/- MUC13631
PF_Lepr-/- MUC13632
VSG_Lepr-/- MUC13640
VSG_Lepr-/- MUC13642
0.00
0.25
0.50
0.75
1.00

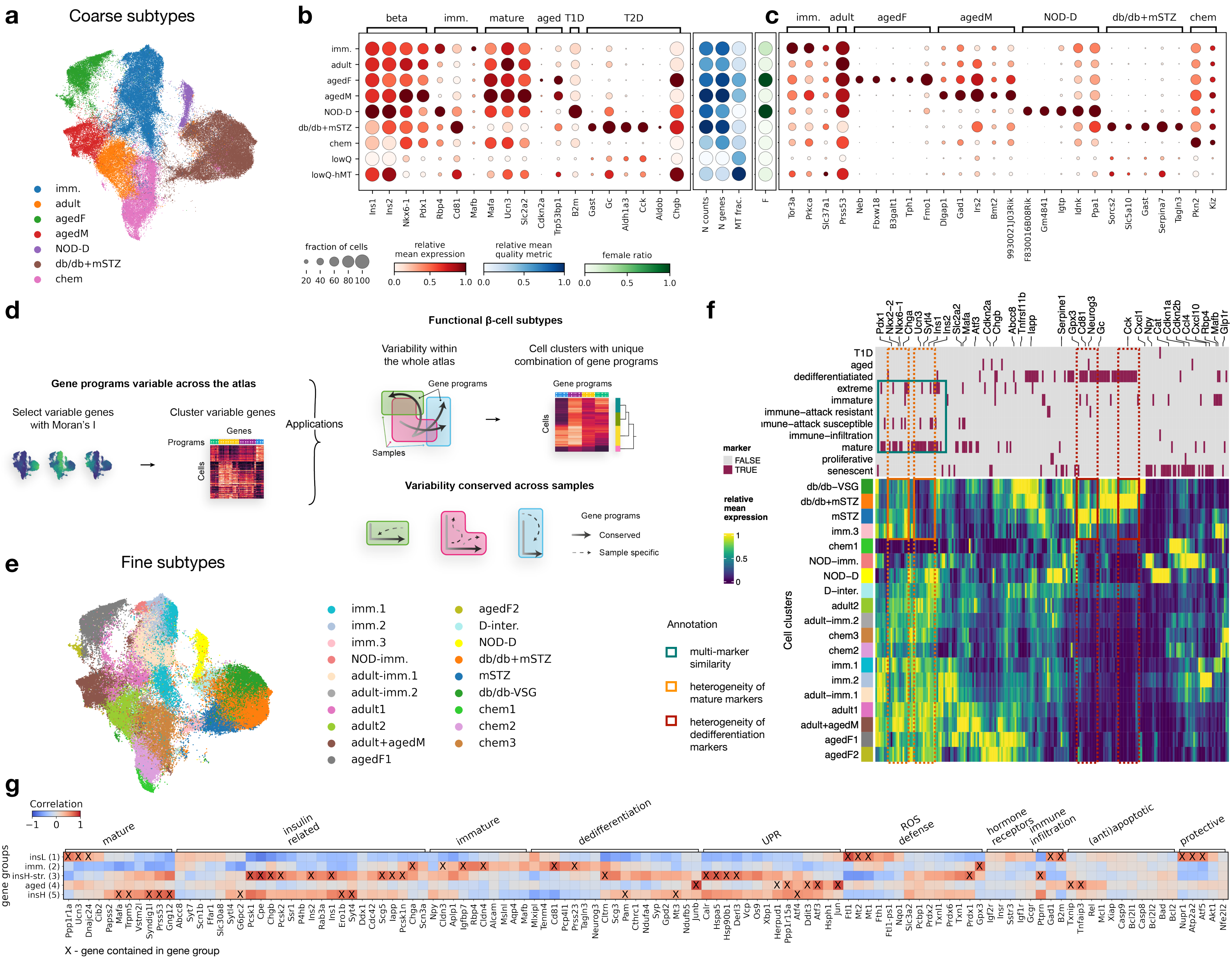

a
Coarse subtypes
imm.
adult
agedF
agedM
NOD-D
db/db+mSTZ
chem
b
beta
imm.
mature
aged
T1D
T2D
imm.
adult
agedF
agedM
NOD-D
db/db+mSTZ
chem
lowQ
lowQ-hMT
Ins1
Ins2
Nkx6-1
Pdx1
Rbp4
Cd81
Mafb
Mafa
Ucn3
Slc2a2
Cdkn2a
Trp53bp1
B2m
Gast
Gc
Aldh1a3
Cck
Aldob
Chgb
N counts
N genes
MT frac.
F
fraction of cells
20 40 60 80 100
relative mean expression
relative mean quality metric
female ratio
0.0 0.5 1.0
c
imm.
adult
agedF
agedM
NOD-D
db/db+mSTZ
chem
Tor3a
Prkca
Slc37a1
Prss53
Neb
Fbxw18
B3galt1
Tph1
Fmo1
Dlgap1
Gad1
Irs2
Bmt2
9930021J03Rik
F830016B08Rik
Gm4841
Iigp
Idnk
Ppa1
Sorcs2
Slc5a10
Gast
Serpina7
Tagln3
Pkn2
Kiz
d
Gene programs variable across the atlas
Select variable genes with Moran's I
Cluster variable genes
Programs
Genes
Cells
Applications
Functional β-cell subtypes
Variability within the whole atlas
Cell clusters with unique combination of gene programs
Gene programs
Samples
Variability conserved across samples
Gene programs
Conserved
Sample specific
e
Fine subtypes
imm.1
imm.2
imm.3
NOD-imm.
adult-imm.1
adult-imm.2
adult1
adult2
adult+agedM
agedF1
agedF2
D-inter.
NOD-D
db/db+mSTZ
mSTZ
db/db-VSG
chem1
chem2
chem3
Annotation
multi-marker similarity
heterogeneity of mature markers
heterogeneity of dedifferentiation markers
f
T1D
aged
dedifferentiatied
extreme
immature
immune−attack resistant
immune−attack susceptible
immune−infiltration
mature
proliferative
senescent
marker
FALSE
TRUE
relative mean expression
1
0.5
0
Cell clusters
db/db−VSG
db/db+mSTZ
mSTZ
imm.3
chem1
NOD−imm.
NOD−D
D−inter.
adult2
adult−imm.2
chem3
chem2
imm.1
imm.2
adult−imm.1
adult1
adult+agedM
agedF1
agedF2
g
Correlation
−1 0 1
gene groups
insL (1)
imm. (2)
insH-str. (3)
aged (4)
insH (5)
mature
insulin related
immature
dedifferentiation
UPR
ROS defense
hormone receptors
immune infiltration
(anti)apoptotic
protective
X - gene contained in gene group

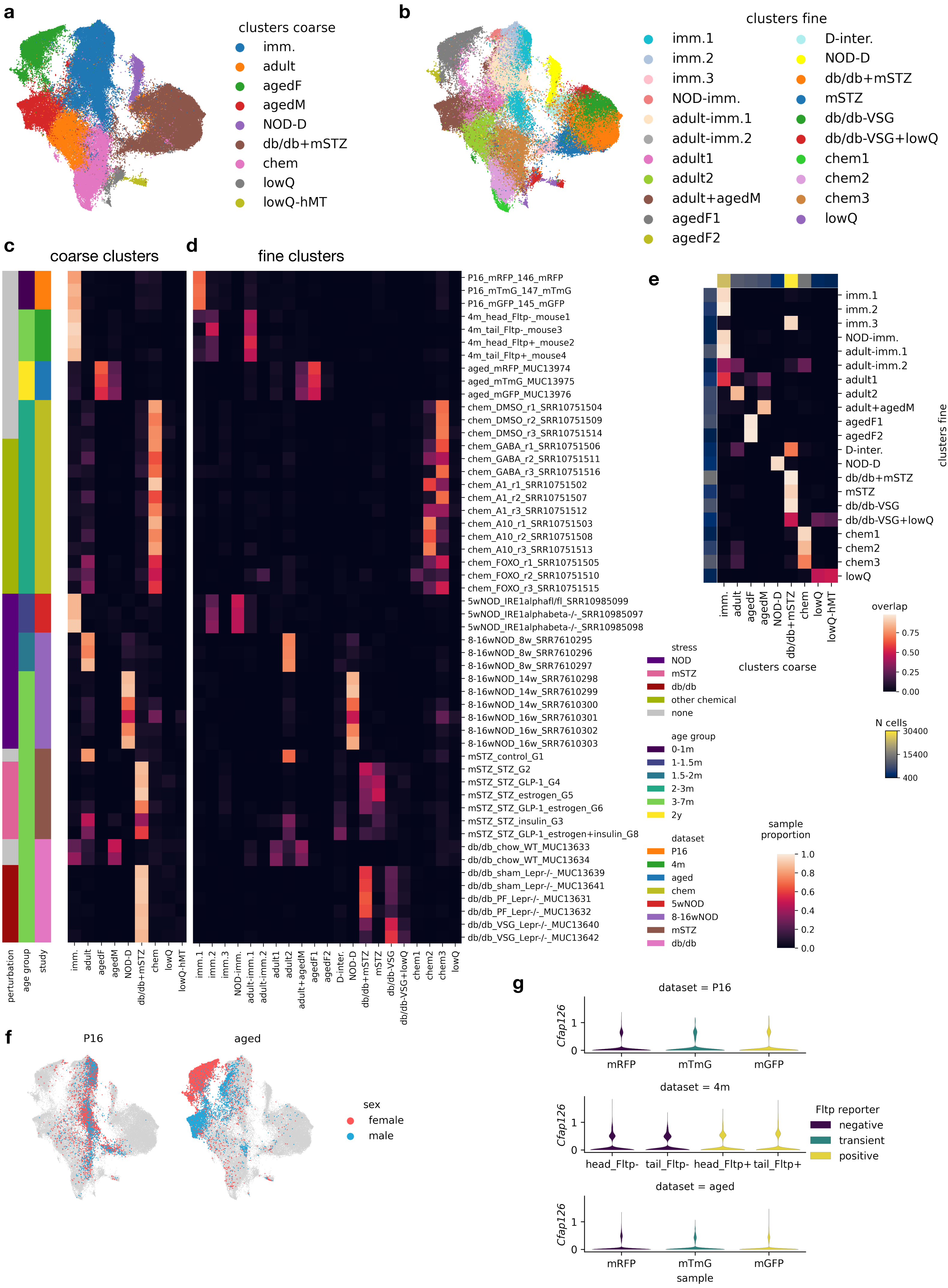

a
clusters coarse
imm.
adult
agedF
agedM
NOD-D
db/db+mSTZ
chem
lowQ
lowQ-hMT
b
clusters fine
imm.1
imm.2
imm.3
NOD-imm.
adult-imm.1
adult-imm.2
adult1
adult2
adult+agedM
agedF1
agedF2
D-inter.
NOD-D
db/db+mSTZ
mSTZ
db/db-VSG
db/db-VSG+lowQ
chem1
chem2
chem3
lowQ
c
coarse clusters
d
fine clusters
P16_mRFP_146_mRFP
P16_mTmG_147_mTmG
P16_mGFP_145_mGFP
4m_head_Fltp-_mouse1
4m_tail_Fltp-_mouse3
4m_head_Fltp+_mouse2
4m_tail_Fltp+_mouse4
aged_mRFP_MUC13974
aged_mTmG_MUC13975
aged_mGFP_MUC13976
chem_DMSO_r1_SRR10751504
chem_DMSO_r2_SRR10751509
chem_DMSO_r3_SRR10751514
chem_GABA_r1_SRR10751506
chem_GABA_r2_SRR10751511
chem_GABA_r3_SRR10751516
chem_A1_r1_SRR10751502
chem_A1_r2_SRR10751507
chem_A1_r3_SRR10751512
chem_A10_r1_SRR10751503
chem_A10_r2_SRR10751508
chem_A10_r3_SRR10751513
chem_FOXO_r1_SRR10751505
chem_FOXO_r2_SRR10751510
chem_FOXO_r3_SRR10751515
5wNOD_IRE1alphafl/fl_SRR10985099
5wNOD_IRE1alphabeta-/-_SRR10985097
5wNOD_IRE1alphabeta-/-_SRR10985098
8-16wNOD_8w_SRR7610295
8-16wNOD_8w_SRR7610296
8-16wNOD_8w_SRR7610297
8-16wNOD_14w_SRR7610298
8-16wNOD_14w_SRR7610299
8-16wNOD_14w_SRR7610300
8-16wNOD_16w_SRR7610301
8-16wNOD_16w_SRR7610302
8-16wNOD_16w_SRR7610303
mSTZ_control_G1
mSTZ_STZ_G2
mSTZ_STZ_GLP-1_G4
mSTZ_STZ_estrogen_G5
mSTZ_STZ_GLP-1_estrogen_G6
mSTZ_STZ_insulin_G3
mSTZ_STZ_GLP-1_estrogen+insulin_G8
db/db_chow_WT_MUC13633
db/db_chow_WT_MUC13634
db/db_sham_Lepr-/-_MUC13639
db/db_sham_Lepr-/-_MUC13641
db/db_PF_Lepr-/-_MUC13631
db/db_PF_Lepr-/-_MUC13632
db/db_VSG_Lepr-/-_MUC13640
db/db_VSG_Lepr-/-_MUC13642
perturbation
age group
study
imm.
adult
agedF
agedM
NOD-D
db/db+mSTZ
chem
lowQ
lowQ-hMT
imm.1
imm.2
imm.3
NOD-imm.
adult-imm.1
adult-imm.2
adult1
adult2
adult+agedM
agedF1
agedF2
D-inter.
NOD-D
db/db+mSTZ
mSTZ
db/db-VSG
db/db-VSG+lowQ
chem1
chem2
chem3
lowQ
stress
NOD
mSTZ
db/db
other chemical
none
age group
0-1m
1-1.5m
1.5-2m
2-3m
3-7m
2y
dataset
P16
4m
aged
chem
5wNOD
8-16wNOD
mSTZ
db/db
sample proportion
1.0
0.8
0.6
0.4
0.2
0.0
e
imm.1
imm.2
imm.3
NOD-imm.
adult-imm.1
adult-imm.2
adult1
adult2
adult+agedM
agedF1
agedF2
D-inter.
NOD-D
db/db+mSTZ
mSTZ
db/db-VSG
db/db-VSG+lowQ
chem1
chem2
chem3
lowQ
clusters fine
imm.
adult
agedF
agedM
NOD-D
db/db+mSTZ
chem
lowQ
lowQ-hMT
clusters coarse
overlap
0.75
0.50
0.25
0.00
N cells
30400
15400
400
f
P16
aged
sex
female
male
g
dataset = P16
Cfap126
1
0
mRFP
mTmG
mGFP
dataset = 4m
Cfap126
1
0
head_Fltp-
tail_Fltp-
head_Fltp+
tail_Fltp+
Fltp reporter
negative
transient
positive
dataset = aged
Cfap126
1
0
mRFP
mTmG
mGFP
sample

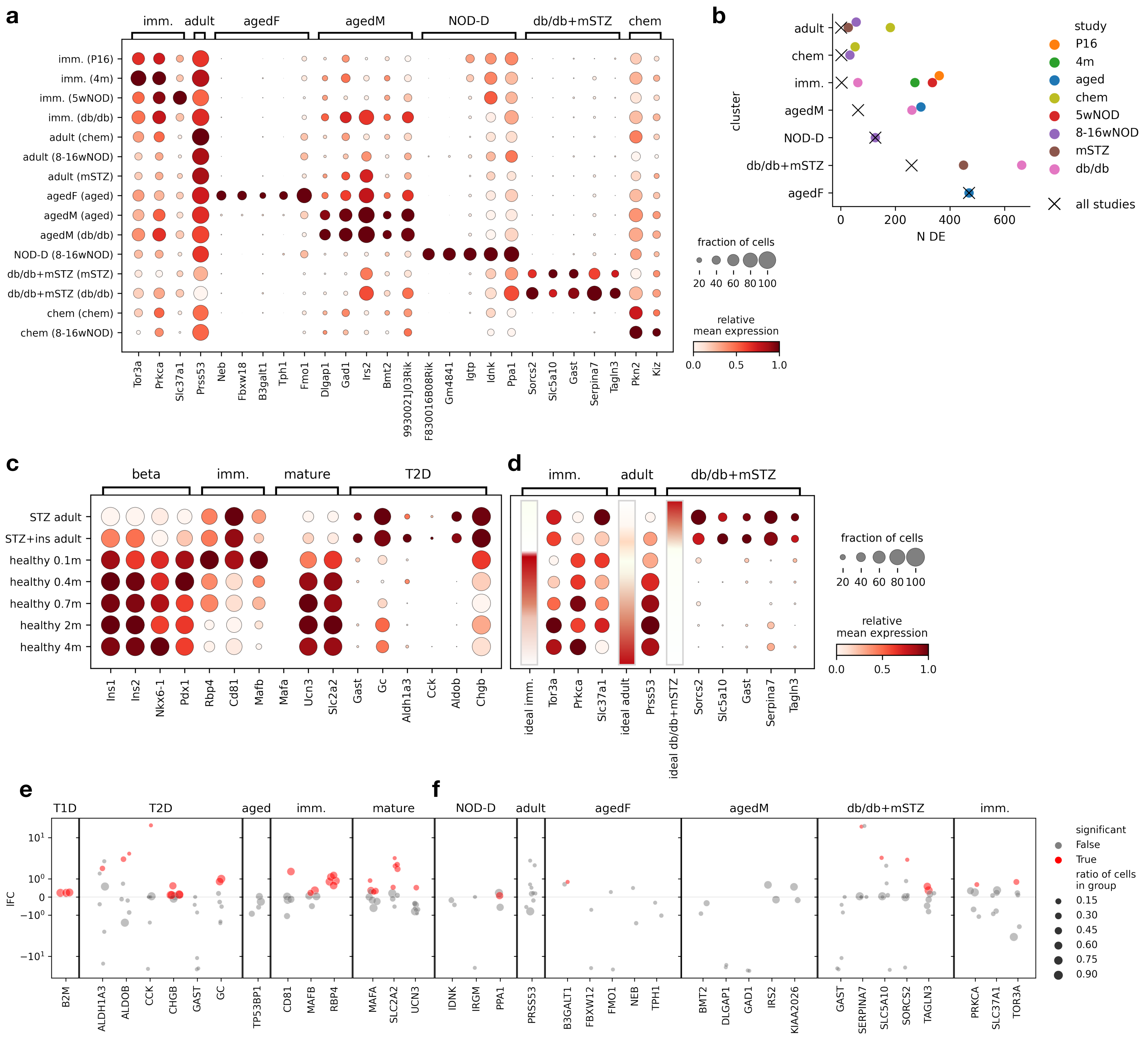
a
imm.
adult
agedF
agedM
NOD-D
db/db+mSTZ
chem
imm. (P16)
imm. (4m)
imm. (5wNOD)
imm. (db/db)
adult (chem)
adult (8-16wNOD)
adult (mSTZ)
agedF (aged)
agedM (aged)
agedM (db/db)
NOD-D (8-16wNOD)
db/db+mSTZ (mSTZ)
db/db+mSTZ (db/db)
chem (chem)
chem (8-16wNOD)
Tor3a
Prkca
Slc37a1
Prss53
Neb
Fbxw18
B3galt1
Tph1
Fmo1
Dlgap1
Gad1
Irs2
Bmt2
9930021J03Rik
F830016B08Rik
Gm4841
Igtp
Idnk
Ppa1
Sorcs2
Slc5a10
Gast
Serpina7
Tagln3
Pkn2
Kiz
fraction of cells
20 40 60 80 100
relative
mean expression
0.0 0.5 1.0
b
adult
chem
imm.
agedM
NOD-D
db/db+mSTZ
agedF
cluster
0 200 400 600
N DE
study
P16
4m
aged
chem
5wNOD
8-16wNOD
mSTZ
db/db
all studies
c
beta
imm.
mature
T2D
STZ adult
STZ+ins adult
healthy 0.1m
healthy 0.4m
healthy 0.7m
healthy 2m
healthy 4m
Ins1
Ins2
Nkx6-1
Pdx1
Rbp4
Cd81
Mafb
Mafa
Ucn3
Slc2a2
Gast
Gc
Aldh1a3
Cck
Aldob
Chgb
d
imm.
adult
db/db+mSTZ
ideal imm.
Tor3a
Prkca
Slc37a1
ideal adult
Prss53
ideal db/db+mSTZ
Sorcs2
Slc5a10
Gast
Serpina7
Tagln3
fraction of cells
20 40 60 80 100
relative
mean expression
0.0 0.5 1.0
e
f
T1D
T2D
aged
imm.
mature
NOD-D
adult
agedF
agedM
db/db+mSTZ
imm.
lFC
$10^1$
$10^0$
0
$-10^0$
$-10^1$
B2M
ALDH1A3
ALDOB
CCK
CHGB
GAST
GC
TP53BP1
CD81
MAFB
RBP4
MAFA
SLC2A2
UCN3
IDNK
IRGM
PPA1
PRSS53
B3GALT1
FBXW12
FMO1
NEB
TPH1
BMT2
DLGAP1
GAD1
IRS2
KIAA2026
GAST
SERPINA7
SLC5A10
SORCS2
TAGLN3
PRKCA
SLC37A1
TOR3A
significant
False
True
ratio of cells
in group
0.15
0.30
0.45
0.60
0.75
0.90

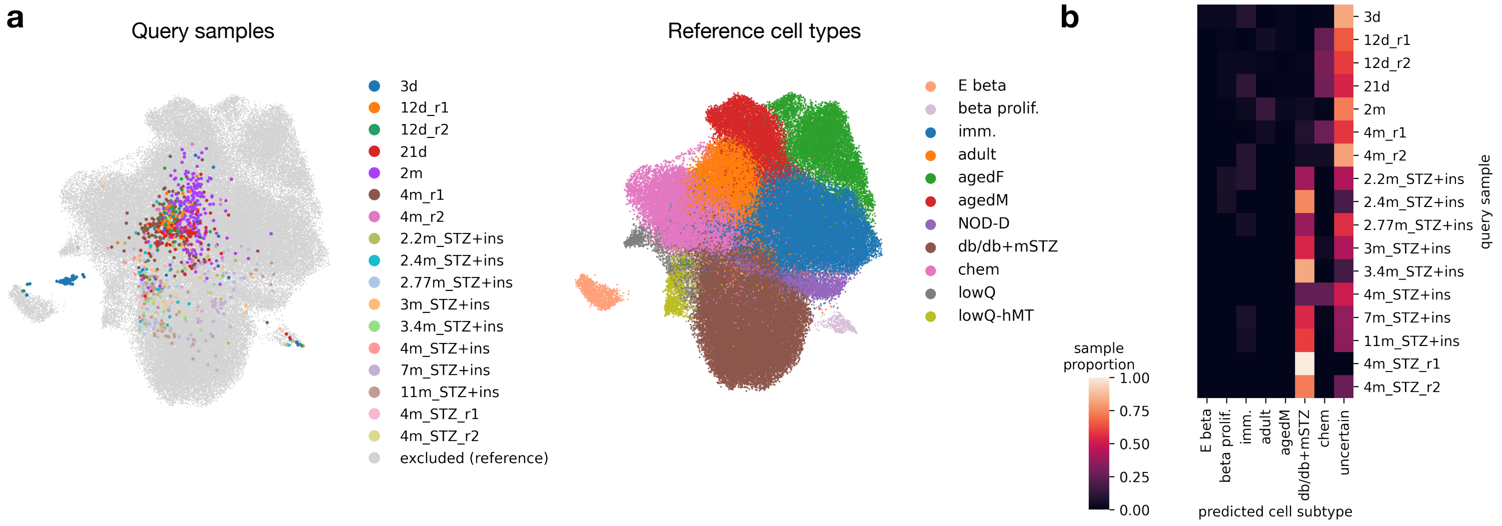
a
Query samples
3d
12d_r1
12d_r2
21d
2m
4m_r1
4m_r2
2.2m_STZ+ins
2.4m_STZ+ins
2.77m_STZ+ins
3m_STZ+ins
3.4m_STZ+ins
4m_STZ+ins
7m_STZ+ins
11m_STZ+ins
4m_STZ_r1
4m_STZ_r2
excluded (reference)
Reference cell types
E beta
beta prolif.
imm.
adult
agedF
agedM
NOD-D
db/db+mSTZ
chem
lowQ
lowQ-hMT
b
sample proportion
1.00
0.75
0.50
0.25
0.00
E beta
beta prolif.
imm.
adult
agedM
db/db+mSTZ
chem
uncertain
predicted cell subtype
query sample

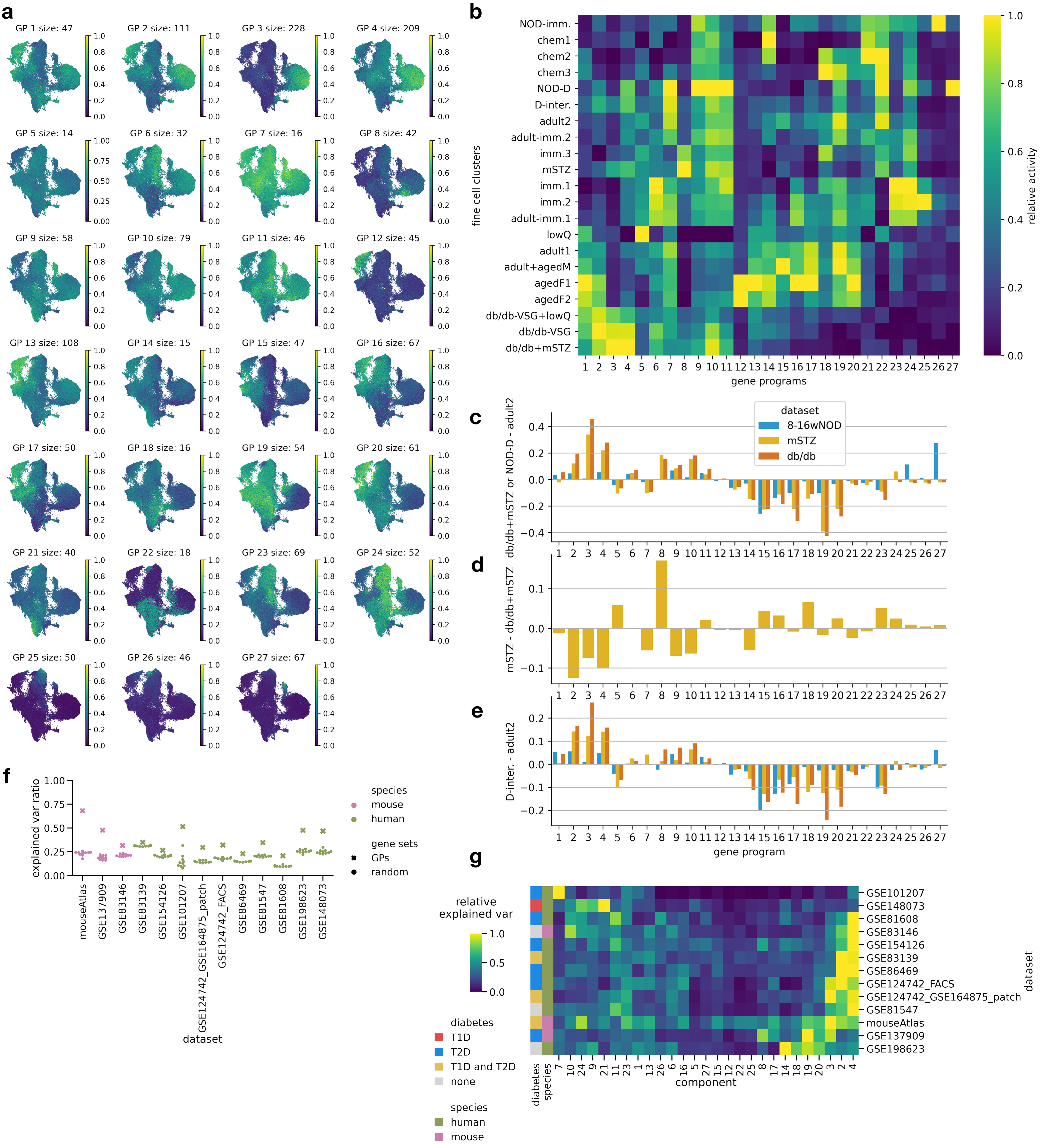
a
GP 1 size: 47
GP 2 size: 111
GP 3 size: 228
GP 4 size: 209
GP 5 size: 14
GP 6 size: 32
GP 7 size: 16
GP 8 size: 42
GP 9 size: 58
GP 10 size: 79
GP 11 size: 46
GP 12 size: 45
GP 13 size: 108
GP 14 size: 15
GP 15 size: 47
GP 16 size: 67
GP 17 size: 50
GP 18 size: 16
GP 19 size: 54
GP 20 size: 61
GP 21 size: 40
GP 22 size: 18
GP 23 size: 69
GP 24 size: 52
GP 25 size: 50
GP 26 size: 46
GP 27 size: 67
b
NOD-imm.
chem1
chem2
chem3
NOD-D
D-inter.
adult2
adult-imm.2
imm.3
mSTZ
imm.1
imm.2
adult-imm.1
lowQ
adult1
adult+agedM
agedF1
agedF2
db/db-VSG+lowQ
db/db-VSG
db/db+mSTZ
fine cell clusters
gene programs
relative activity
c
dataset
8-16wNOD
mSTZ
db/db
db/db+mSTZ or NOD-D - adult2
d
mSTZ - db/db+mSTZ
e
D-inter. - adult2
gene program
f
explained var ratio
species
mouse
human
gene sets
GPs
random
mouseAtlas
GSE137909
GSE83146
GSE83139
GSE154126
GSE101207
GSE124742_GSE164875_patch
GSE124742_FACS
GSE86469
GSE81547
GSE81608
GSE198623
GSE148073
dataset
g
relative explained var
diabetes
T1D
T2D
T1D and T2D
none
species
human
mouse
GSE101207
GSE148073
GSE81608
GSE83146
GSE154126
GSE83139
GSE86469
GSE124742_FACS
GSE124742_GSE164875_patch
GSE81547
mouseAtlas
GSE137909
GSE198623
dataset
diabetes
species
component

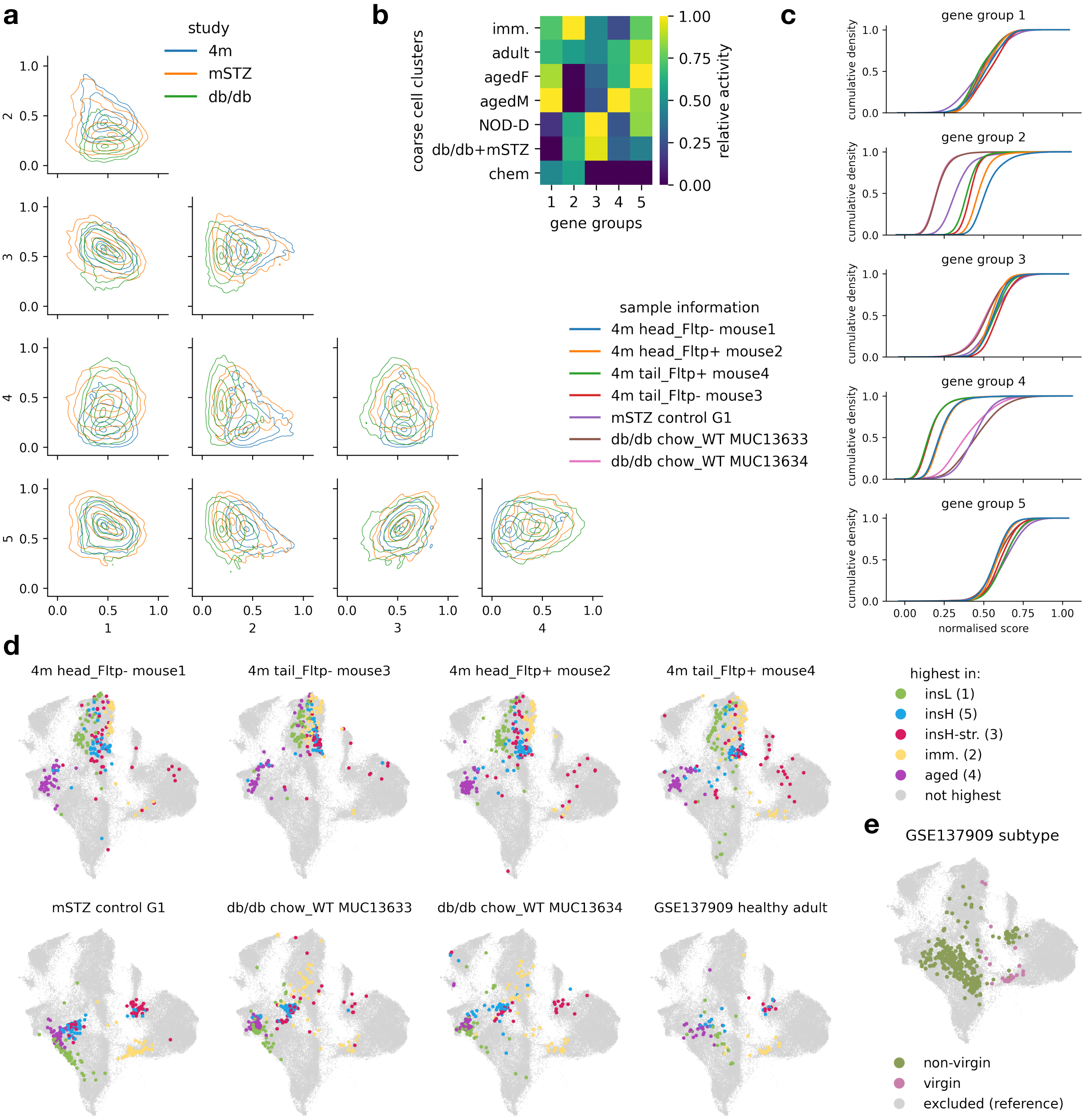

a
study
4m
mSTZ
db/db
b
coarse cell clusters
imm.
adult
agedF
agedM
NOD-D
db/db+mSTZ
chem
gene groups
relative activity
c
gene group 1
gene group 2
gene group 3
gene group 4
gene group 5
cumulative density
normalised score
sample information
4m head_Fltp- mouse1
4m head_Fltp+ mouse2
4m tail_Fltp+ mouse4
4m tail_Fltp- mouse3
mSTZ control G1
db/db chow_WT MUC13633
db/db chow_WT MUC13634
d
4m head_Fltp- mouse1
4m tail_Fltp- mouse3
4m head_Fltp+ mouse2
4m tail_Fltp+ mouse4
mSTZ control G1
db/db chow_WT MUC13633
db/db chow_WT MUC13634
GSE137909 healthy adult
highest in:
insL (1)
insH (5)
insH-str. (3)
imm. (2)
aged (4)
not highest
e
GSE137909 subtype
non-virgin
virgin
excluded (reference)

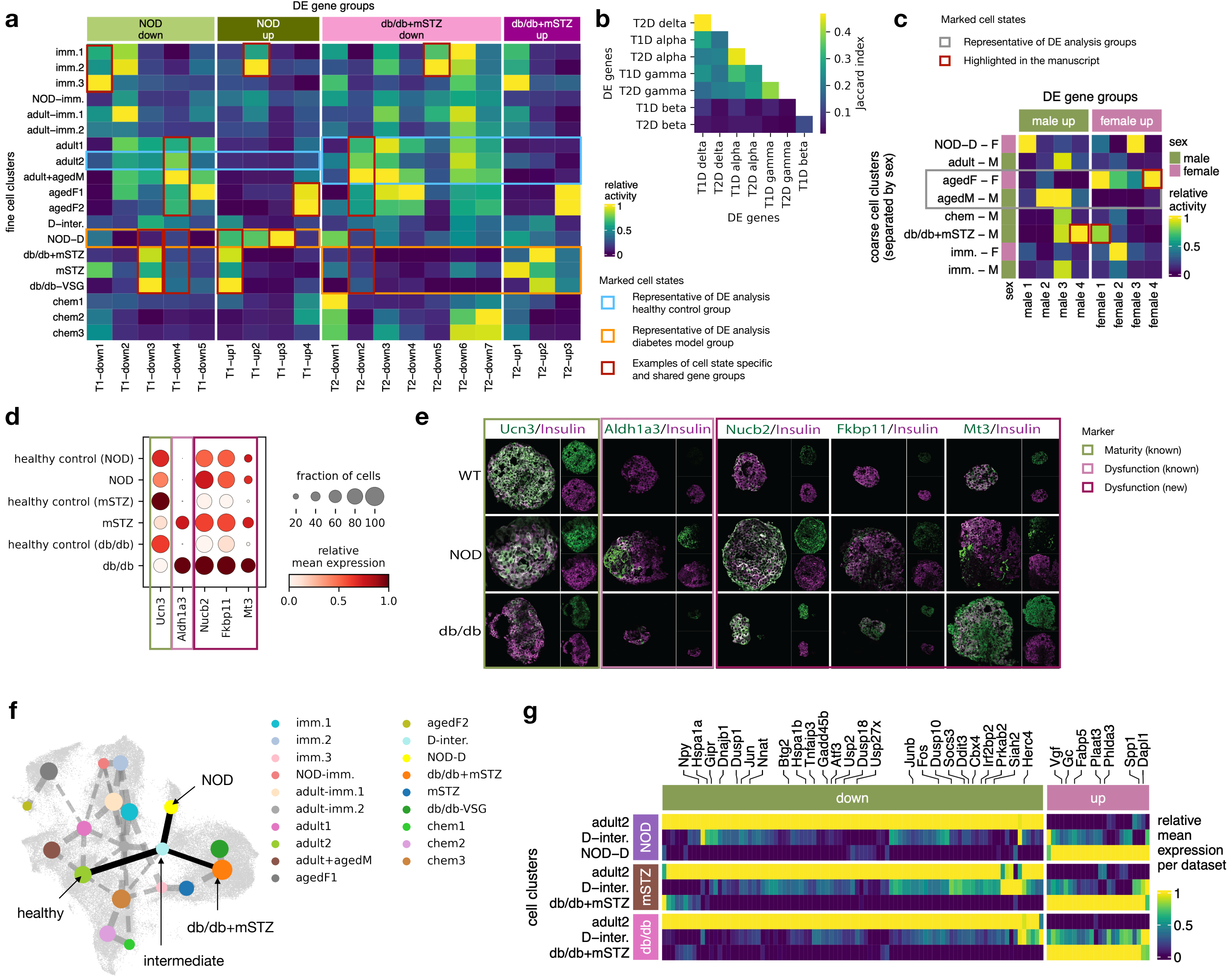
a
DE gene groups
NOD down
NOD up
db/db+mSTZ down
db/db+mSTZ up
fine cell clusters
imm.1
imm.2
imm.3
NOD–imm.
adult–imm.1
adult–imm.2
adult1
adult2
adult+agedM
agedF1
agedF2
D–inter.
NOD–D
db/db+mSTZ
mSTZ
db/db–VSG
chem1
chem2
chem3
T1–down1 T1–down2 T1–down3 T1–down4 T1–down5 T1–up1 T1–up2 T1–up3 T1–up4 T2–down1 T2–down2 T2–down3 T2–down4 T2–down5 T2–down6 T2–down7 T2–up1 T2–up2 T2–up3
b
T2D delta
T1D alpha
T2D alpha
T1D gamma
T2D gamma
T1D beta
T2D beta
DE genes
T1D delta T2D delta T1D alpha T2D alpha T1D gamma T2D gamma T1D beta
Jaccard index
0.1 0.2 0.3 0.4
relative activity
0 0.5 1
Marked cell states
Representative of DE analysis healthy control group
Representative of DE analysis diabetes model group
Examples of cell state specific and shared gene groups
c
Marked cell states
Representative of DE analysis groups
Highlighted in the manuscript
DE gene groups
male up
female up
coarse cell clusters (separated by sex)
NOD–D – F
adult – M
agedF – F
agedM – M
chem – M
db/db+mSTZ – M
imm. – F
imm. – M
sex
male 1 male 2 male 3 male 4 female 1 female 2 female 3 female 4
sex
male
female
relative activity
0 0.5 1
d
healthy control (NOD)
NOD
healthy control (mSTZ)
mSTZ
healthy control (db/db)
db/db
Ucn3 Aldh1a3 Nucb2 Fkbp11 Mt3
fraction of cells
20 40 60 80 100
relative mean expression
0.0 0.5 1.0
e
Ucn3/Insulin
Aldh1a3/Insulin
Nucb2/Insulin
Fkbp11/Insulin
Mt3/Insulin
WT
NOD
db/db
Marker
Maturity (known)
Dysfunction (known)
Dysfunction (new)
f
NOD
healthy
intermediate
db/db+mSTZ
imm.1
imm.2
imm.3
NOD-imm.
adult-imm.1
adult-imm.2
adult1
adult2
adult+agedM
agedF1
agedF2
D-inter.
NOD-D
db/db+mSTZ
mSTZ
db/db-VSG
chem1
chem2
chem3
g
Npy Hspa1a Gipr Dnajb1 Dusp1 Jun Nnat Btg2 Hspa1b Tnfaip3 Gadd45b Atf3 Usp2 Dusp18 Usp27x Junb Fos Dusp10 Socs3 Ddit3 Cbx4 Irf2bp2 Prkab2 Siah2 Herc4 Vgf Gc Fabp5 Plaat3 Phlda3 Spp1 Dapl1
down
up
cell clusters
NOD
adult2
D–inter.
NOD–D
mSTZ
adult2
D–inter.
db/db+mSTZ
db/db
adult2
D–inter.
db/db+mSTZ
relative mean expression per dataset
0 0.5 1

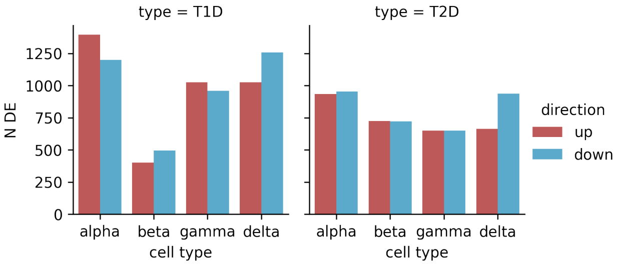
type = T1D
type = T2D
N DE
0
250
500
750
1000
1250
alpha
beta
gamma
delta
cell type
direction
up
down

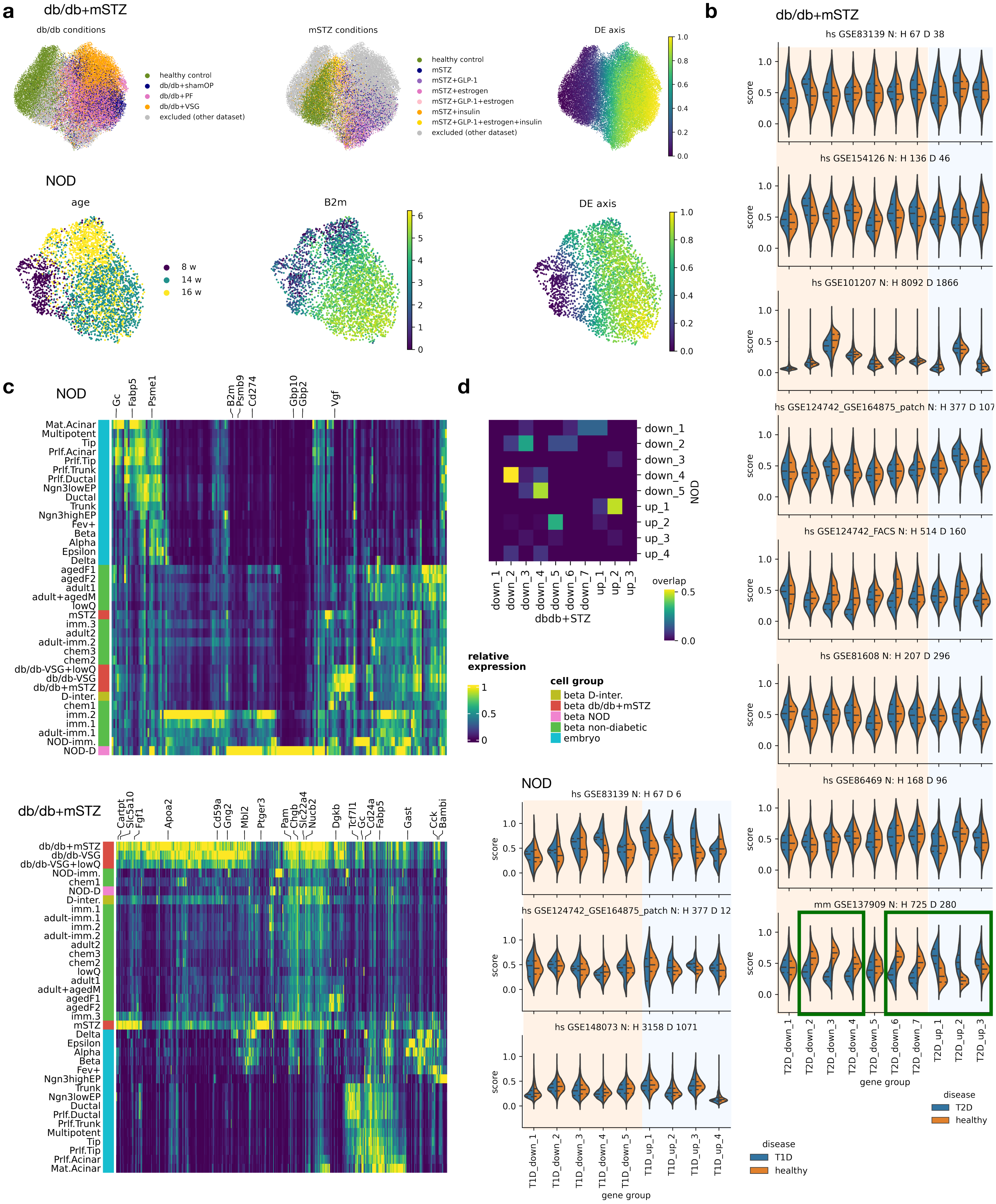
a
db/db+mSTZ
db/db conditions
healthy control
db/db+shamOP
db/db+PF
db/db+VSG
excluded (other dataset)
mSTZ conditions
healthy control
mSTZ
mSTZ+GLP-1
mSTZ+estrogen
mSTZ+GLP-1+estrogen
mSTZ+insulin
mSTZ+GLP-1+estrogen+insulin
excluded (other dataset)
DE axis
NOD
age
8 w
14 w
16 w
B2m
DE axis
b
db/db+mSTZ
hs GSE83139 N: H 67 D 38
hs GSE154126 N: H 136 D 46
hs GSE101207 N: H 8092 D 1866
hs GSE124742_GSE164875_patch N: H 377 D 107
hs GSE124742_FACS N: H 514 D 160
hs GSE81608 N: H 207 D 296
hs GSE86469 N: H 168 D 96
mm GSE137909 N: H 725 D 280
score
T2D_down_1
T2D_down_2
T2D_down_3
T2D_down_4
T2D_down_5
T2D_down_6
T2D_down_7
T2D_up_1
T2D_up_2
T2D_up_3
gene group
disease
T2D
healthy
c
NOD
Gc
Fabp5
Psme1
B2m
Psmb9
Cd274
Gbp10
Gbp2
Vgf
Mat.Acinar
Multipotent
Tip
Prlf.Acinar
Prlf.Tip
Prlf.Trunk
Prlf.Ductal
Ngn3lowEP
Ductal
Trunk
Ngn3highEP
Fev+
Beta
Alpha
Epsilon
Delta
agedF1
agedF2
adult1
adult+agedM
lowQ
mSTZ
imm.3
adult2
adult-imm.2
chem3
chem2
db/db-VSG+lowQ
db/db-VSG
db/db+mSTZ
D-inter.
chem1
imm.2
imm.1
adult-imm.1
NOD-imm.
NOD-D
db/db+mSTZ
Cartpt
Slc5a10
Fgf1
Apoa2
Cd59a
Gng2
Mbl2
Ptger3
Pam
Chgb
Slc22a4
Nucb2
Dgkb
Tcf7l1
Gc
Cd24a
Fabp5
Gast
Cck
Bambi
relative expression
cell group
beta D-inter.
beta db/db+mSTZ
beta NOD
beta non-diabetic
embryo
d
down_1
down_2
down_3
down_4
down_5
up_1
up_2
up_3
up_4
NOD
down_6
down_7
dbdb+STZ
overlap
NOD
hs GSE83139 N: H 67 D 6
hs GSE124742_GSE164875_patch N: H 377 D 12
hs GSE148073 N: H 3158 D 1071
T1D_down_1
T1D_down_2
T1D_down_3
T1D_down_4
T1D_down_5
T1D_up_1
T1D_up_2
T1D_up_3
T1D_up_4
gene group
disease
T1D
healthy

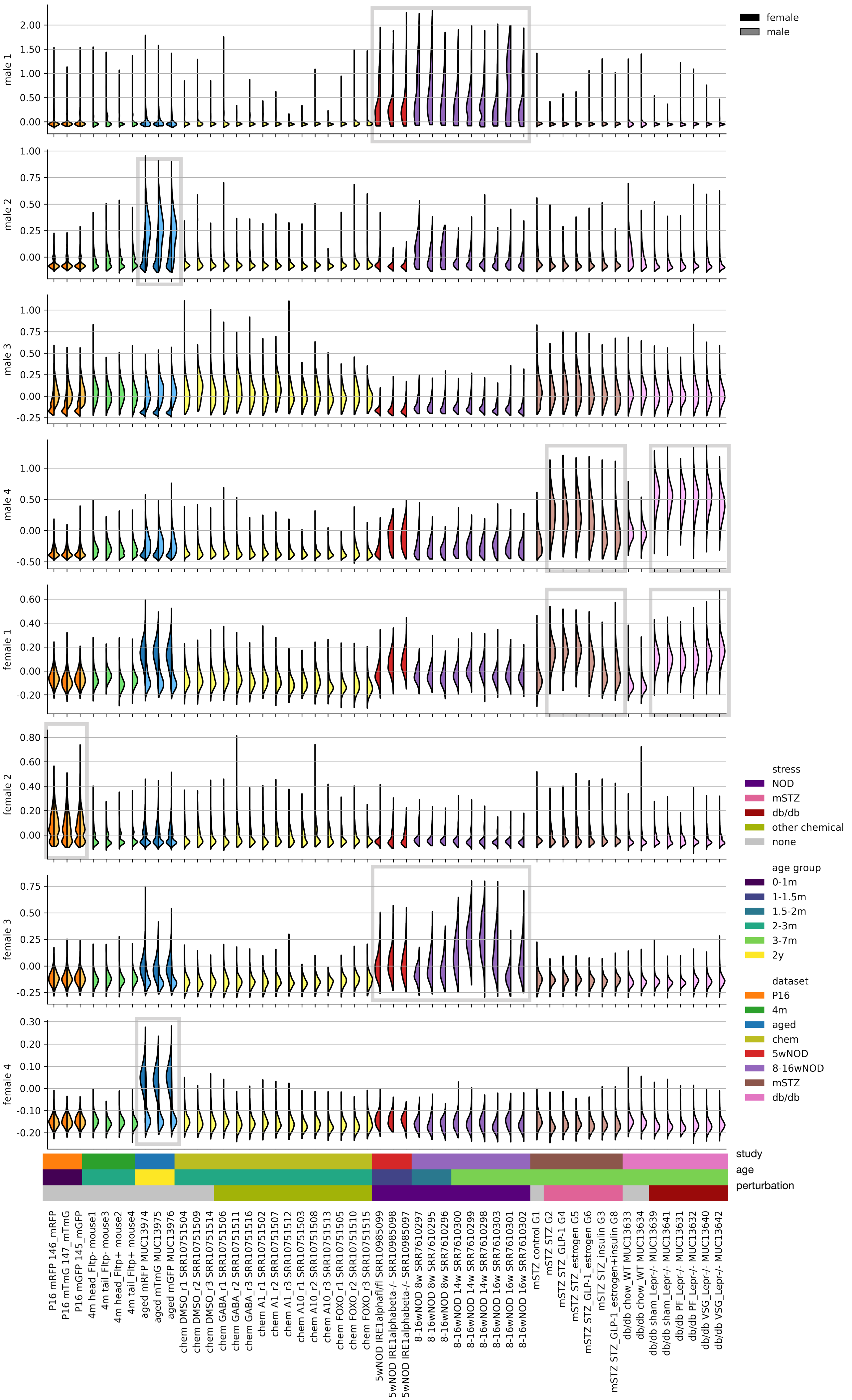

female
male
male 1
male 2
male 3
male 4
female 1
female 2
female 3
female 4
stress
NOD
mSTZ
db/db
other chemical
none
age group
0-1m
1-1.5m
1.5-2m
2-3m
3-7m
2y
dataset
P16
4m
aged
chem
5wNOD
8-16wNOD
mSTZ
db/db
study
age
perturbation
P16 mRFP 146_mRFP
P16 mTmG 147_mTmG
P16 mGFP 145_mGFP
4m head_Fltp- mouse1
4m tail_Fltp- mouse3
4m head_Fltp+ mouse2
4m tail_Fltp+ mouse4
aged mRFP MUC13974
aged mTmG MUC13975
aged mGFP MUC13976
chem DMSO_r1 SRR10751504
chem DMSO_r2 SRR10751509
chem DMSO_r3 SRR10751514
chem GABA_r1 SRR10751506
chem GABA_r2 SRR10751511
chem GABA_r3 SRR10751516
chem A1_r1 SRR10751502
chem A1_r2 SRR10751507
chem A1_r3 SRR10751512
chem A10_r1 SRR10751503
chem A10_r2 SRR10751508
chem A10_r3 SRR10751513
chem FOXO_r1 SRR10751505
chem FOXO_r2 SRR10751510
chem FOXO_r3 SRR10751515
5wNOD IRE1alphafl/fl SRR10985099
5wNOD IRE1alphabeta-/- SRR10985098
5wNOD IRE1alphabeta-/- SRR10985097
8-16wNOD 8w SRR7610297
8-16wNOD 8w SRR7610295
8-16wNOD 8w SRR7610296
8-16wNOD 14w SRR7610300
8-16wNOD 14w SRR7610299
8-16wNOD 14w SRR7610298
8-16wNOD 16w SRR7610303
8-16wNOD 16w SRR7610301
8-16wNOD 16w SRR7610302
mSTZ control G1
mSTZ STZ G2
mSTZ STZ_GLP-1 G4
mSTZ STZ_estrogen G5
mSTZ STZ_GLP-1_estrogen G6
mSTZ STZ_insulin G3
mSTZ STZ_GLP-1_estrogen+insulin G8
db/db chow_WT MUC13633
db/db chow_WT MUC13634
db/db sham_Lepr-/- MUC13639
db/db sham_Lepr-/- MUC13641
db/db PF_Lepr-/- MUC13631
db/db PF_Lepr-/- MUC13632
db/db VSG_Lepr-/- MUC13640
db/db VSG_Lepr-/- MUC13642

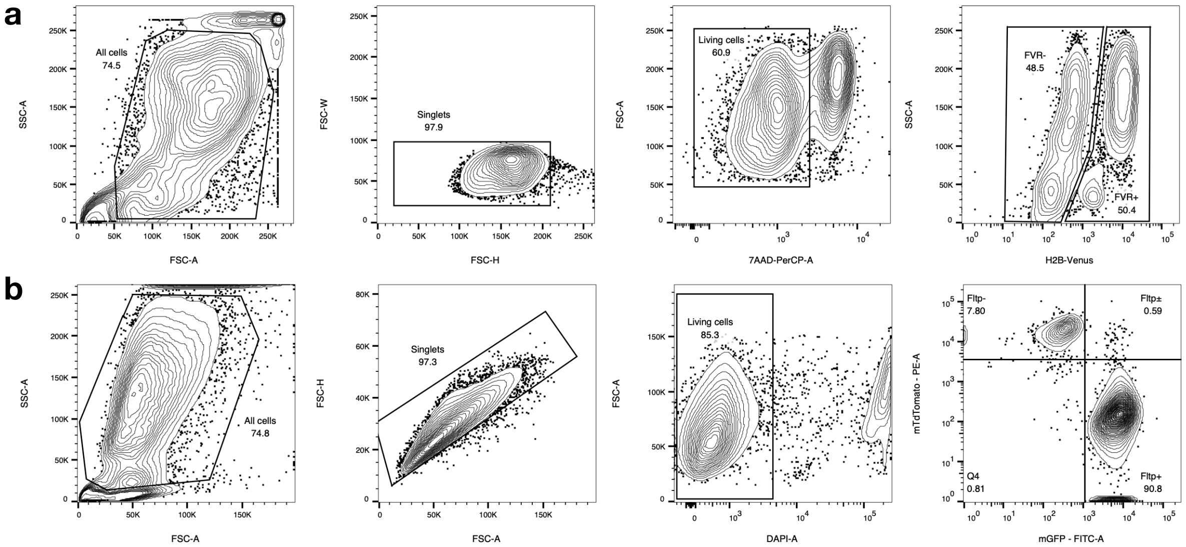
a
All cells
74.5
SSC-A
FSC-A
Singlets
97.9
FSC-W
FSC-H
Living cells
60.9
FSC-A
7AAD-PerCP-A
FVR-
48.5
FVR+
50.4
SSC-A
H2B-Venus
b
All cells
74.8
SSC-A
FSC-A
Singlets
97.3
FSC-H
FSC-A
Living cells
85.3
FSC-A
DAPI-A
Fltp-
7.80
Fltp±
0.59
Q4
0.81
Fltp+
90.8
mTdTomato - PE-A
mGFP - FITC-A
